# A spermatogonial perspective on the expansion of the mammalian brain

**DOI:** 10.1101/2025.09.15.676184

**Authors:** Stephen J. Bush, Anne Goriely

**Author notes:** Correspondence (S.J.B.), (A.G.).

## Abstract

The brain and testis share many molecular similarities, yet the evolutionary implications of this overlap remain unclear. We have previously hypothesised that, throughout evolution, some genetic variants contributing to brain size expansion first arose in spermatogonia where they conferred a selective advantage to the male germline stem cells via ‘selfish spermatogonial selection’, a process analogous to oncogenesis. Once transmitted to the next generation, these selfish variants became constitutive, disproportionately accumulating in signalling pathways active in both spermatogenesis and neurogenesis and which regulate stem cell proliferation. However, testing this hypothesis is stymied by the relative scarcity of spermatogonia and the inherent stochasticity of single-cell transcriptomic profiling. Accordingly, the molecular signatures of spermatogonia are incompletely understood, and their similarity with neural programs difficult to assess. To address this, we combine single-cell testis and brain transcriptomic datasets with multi-tissue proteomic data to show that genes associated with brain-expansion phenotypes, and neurodevelopment generally, are not only widely expressed in the male germline but particularly enriched in spermatogonia, an observation consistent across mammalian evolutionary history. We contextualise these results with an extensive literature survey and conclude that enquiry into the testis-brain connection may yield novel insight into the evolutionary processes that shaped the human condition.

## Introduction

Prominent hallmarks of primate evolution include an increased brain volume and disproportionate expansion of its executive and integrative centre, the cortex ^1,2^. After correcting for allometric scaling with body mass, primates (and humans in particular) have larger brains than almost any other mammal ^3^, due to an increased number of neurons supporting more elaborate neural architectures and cognitive abilities ^4^. A recent review of human brain evolution offers a striking perspective on the extent to which neuron numbers have increased in our species: compared to other great apes, developing humans generate around three million more neurons per hour ^5^. How this came about is an active subject of debate and likely the result of a combination of different factors. In general, the mechanisms producing interspecies differences in neuron number and specialisation implicate the duration, timing, and relative rate of neural stem and progenitor cell (NSPC) divisions ^6^ during development and in what proportion and when mitoses are symmetrical (proliferative, or self-renewing) or asymmetrical (producing differentiation-inclined progeny) ^7^. Consequently, even subtle species-specific differences in NSPC activity can have pronounced effect upon the size and composition of both the brain as a whole, or its regions ^8^. Human-specific variants in, for instance, *ARHGAP11B* ^9^, *NOVA1* ^10^ and *TKTL1* ^11^ influence NSPC proliferation and by extension the overall size of the progenitor pool, and thereby brain. Hence, by modulating the rate of cell division, or by changing developmental decisions, genetic variants in a specific set of genes can alter the number of neurons produced over time and so play causal roles in cortical (and other brain region) expansion.

We have previously drawn attention to selfish spermatogonial selection – a mechanism by which *de novo* mutations affecting stem and progenitor cell regulation in the human testis gain a proliferative advantage and are preferentially introduced into the genome ^12^ – as we believe this may offer a fresh perspective on the molecular and developmental basis of brain expansion and its close connection to other phenotypes, in particular fertility. We have outlined this hypothesis briefly elsewhere ^13^ and illustrate its core principles in **Figure 1**. As our hypothesis approaches the topic of brain expansion from a different area of biology entirely, we refer readers to recent reviews of human brain evolution and development for subject-specific detail (see, for example, ^5,14^ and references therein). Here we focus instead on evaluating the evidence that bears directly on the role of selfish spermatogonial selection. Given this emphasis, a full treatment of other evolutionary forces shaping the brain-testis relationship is beyond the scope of this work, although selected aspects are considered in the **Supplementary Text**.

**Figure 1.**
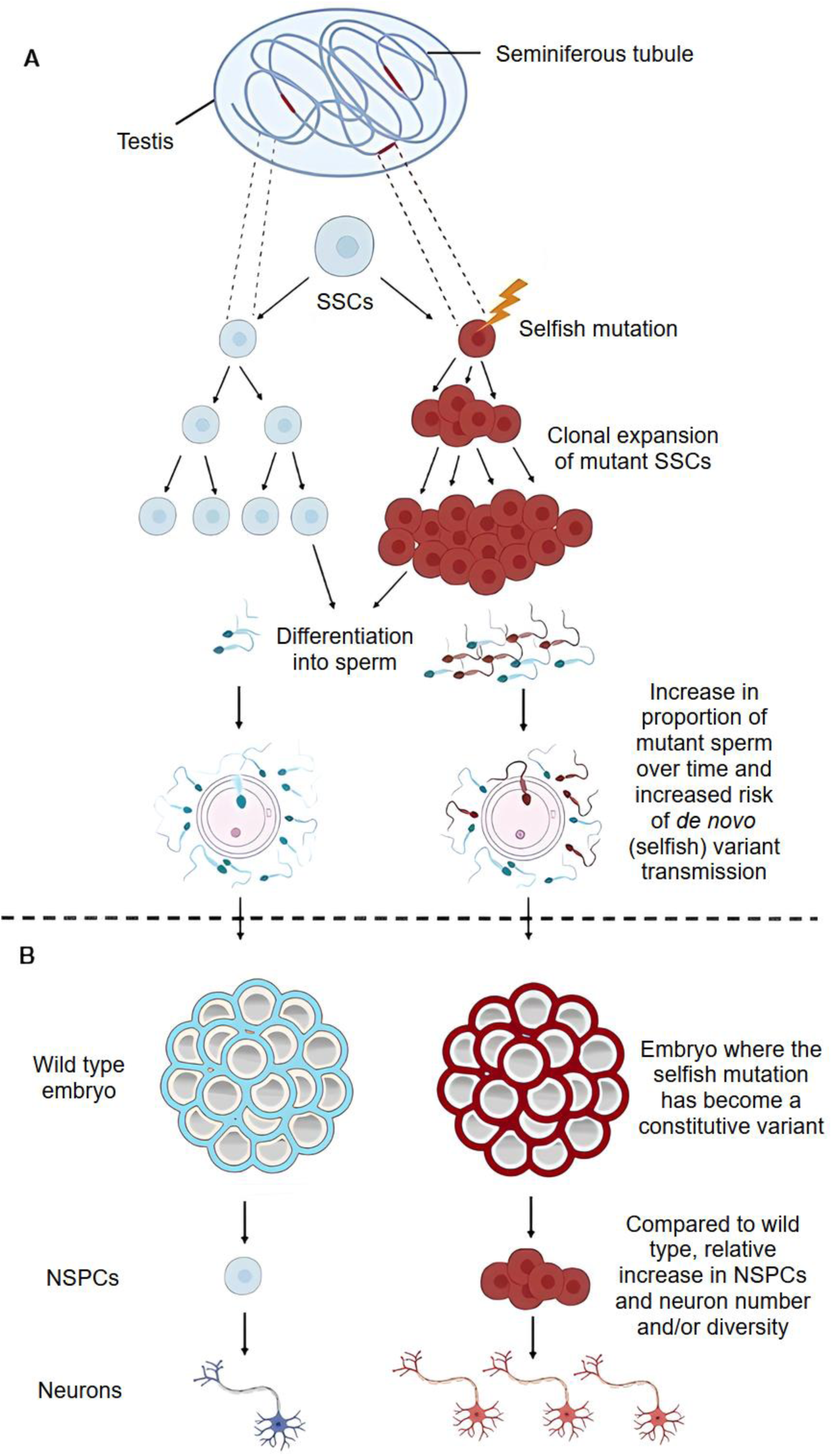
Variants underlying relative increases in brain size (through expansion of the neural progenitor pool) can be introduced into the genome by a male germline-specific evolutionary force, selfish spermatogonial selection. (A) In selfish spermatogonial selection, spontaneous oncogenic-like mutations occur in genes functionally involved in spermatogonial stem cell (SSC) regulation. These selfish mutations confer a selective advantage to SSCs over their wild-type counterparts (blue), resulting in the clonal expansion of mutant SSCs (red) within the seminiferous tubules, and increasing the relative proportion of mutant sperm over time. Sperm carrying selfish mutations often give rise to offspring with neurodevelopmental disorders, although not all selfish mutations are inherently deleterious. This panel is adapted from that originally published in ^12^. (B) As human development begins from a single zygotic cell, this early bottleneck effectively fixes any selfish variants transmitted via sperm in the resulting embryo. As such, ‘mild’ selfish mutations (ones selectively beneficial to SSCs and fertility, yet not overly detrimental when present constitutively in the offspring) may become a source of heritable material and propagate across generations. As selfish selection preferentially enriches the genome for mutations affecting SSC regulation, and given that many of the same genes are also expressed in neural stem and progenitor cells (NSPCs), we propose that this process may have introduced functional variants that expanded the NSPC pool. In principle, such variants could contribute to increased brain size, regional diversification and/or developmental complexity.

Selfish spermatogonial selection is a process taking place during adult spermatogenesis whereby a subset of spontaneously occurring oncogenic-like (‘selfish’) mutations confer a selective advantage to mutant spermatogonial stem cells (SSCs) which then clonally expand along the length of the seminiferous tubules as men age (see reviews ^12,15,16^). As a consequence, selfish mutations disproportionately accrue in the sperm of older men and, compared to classical spontaneous mutations, have a higher likelihood of being inherited as a constitutive mutation in the next generation. To date, the best characterised examples of selfish mutations are pathogenic gain-of-function single nucleotide variants (SNVs), typically (but not exclusively ^17^) occurring within components of the RTK/RAS/MAPK signalling pathway, a central regulator of testicular homeostasis ^18,19^ characteristic of the most primitive SSC state ^20^. Documented examples include variants in the 14 genes *BRAF*, *CBL*, *FGFR2*, *FGFR3*, *HRAS*, *KRAS*, *MAP2K1*, *MAP2K2*, *NRAS*, *PTPN11*, *RAF1*, *RET*, *SMAD4* and *SOS1* (see ^12,17^ and references therein, plus ^21^ for 27 additional candidates). These selfish mutations are consistently associated with severe conditions in the offspring including skeletal dysplasia and neurodevelopmental disorders, such as RASopathies ^15^. While their pathogenic phenotypes are complex and heterogenous (individuals carrying identical RTK/RAS/MAPK mutations can exhibit variable clinical presentations ^22^), cranial overgrowth or macrocephaly, a disproportionately large head, is a recurrent feature ^23^. This is plausibly linked to the roles of many of these genes in regulating cell fate decisions during neurogenesis (discussed further in the **Supplementary Text**).

These observations provide the empirical grounds for our hypothesis – namely, that the clonal dynamics of SSCs have broader implications beyond fertility, and that changes in mammalian brain volume may, at least in part, have been shaped by the selfish properties of the testis. Despite vast differences in structure and function, the testis and brain share many cellular and molecular similarities, including substantial overlap of both their proteomes and transcriptomes ^24^, larger repertoires of splice variants than any other organ ^25^, and – critically for our argument – coordinated cell fate programs, with the neural lineage and the germline co-regulated during early development ^26^. Indeed, not only is there a growing catalogue of genes with pleiotropic functions in both the germline and nervous system ^27^, but many characteristic spermatogonial markers also play functional roles in neurogenesis and brain expansion ^13^ (see also **Supplementary Text**). Moreover, both lineages exhibit similar age-dependent mutational signatures, in particular SBS5. Under a recent model, this signature arises when multiple sources of DNA damage converge into a common mutational outcome, producing similar signatures across diverse cell types. This has been interpreted to suggest the output of common, tightly coupled, endogenous processes; that is, in drawing on the same molecular machinery, the germline and neural lineages accrue the majority of their mutations the same way ^28^.

As an evolutionary force, selfish selection favours the introduction into our genome of functional variants that alter specific molecular pathways modulating SSC self-renewal ^13^. As such, when mutant SSCs differentiate, they generate disproportionate numbers of sperm carrying the selfish allele. Provided that the resulting heterozygous variant is not overly deleterious to the somatic health of the offspring, it can become heritable and persist across generations. Importantly, tight control of the self-renewal of stem cells at the onset of neurogenesis is essential for generating a large pool of progenitors (see review ^29^), the size of which predicts both mature adult brain volume and the relative size of brain regions ^30^. Consistent with this, mathematical modelling suggests that an additional round of symmetric – rather than asymmetric (neurogenic) divisions – by neuroepithelial stem cells would double the number of neurons in the cortex ^31^. Further supporting a mechanistic link between male reproduction and neurogenesis, studies of human brain organoids show that androgens, key regulators of spermatogenesis, can increase the number of proliferative divisions in cortical progenitors ^32^.

Female reproductive tissues (which, like the testis, possess unusually broad proteomic overlap with the brain ^33^) may also be relevant to brain expansion; they are enriched in genes associated with endocranial globularity, a uniquely human phenotype reflecting the timing and pattern of brain growth ^34^. Similarly, a recent study which performed *in silico* modelling of hominin brain expansion argued that increased brain size may have arisen not from direct selection on cognitive ability but by developmental constraints on fertility (specifically, affecting the number of ovarian follicles) ^35^. Although the study lacked data for males, these constraints may have diverted evolutionary change in a direction that led to brain expansion, such that this phenotype evolved as a spandrel ^36^.

While these observations point to intriguing connections between the germline and brain development, our aim here is to evaluate this relationship more systematically. The outstanding challenge in characterising molecular similarities between the brain and either spermatogonia or SSCs (a subset of undifferentiated spermatogonia ^37^) is relative paucity of data: not only are spermatogonia a rare cellular population ^38^ but compared to other human tissues, relatively few single-cell transcriptomic datasets are available for testis ^39^. Accordingly, the molecular identity of spermatogonia, and the heterogeneity of their subpopulations, remains incompletely understood ^40^. To address this limitation, we had previously created an integrated single-cell atlas of 34 publicly-available adult human testis samples, with the aim of increasing the resolution at which spermatogonial states and their developmental trajectory could be defined ^41^. Using this atlas, alongside other datasets discussed below, here we assess the extent to which genes functionally associated with brain expansion phenotypes and neurodevelopment are also expressed in the testis. Our data support the existence of common molecular programs to both spermato- and neurogenesis and we argue that, on balance, there is a plausible role for the male germline, in at least some capacity, in brain expansion.

More specifically, we combine transcriptomic data from multiple single-cell brain ^42,43^ and testis atlases (our own ^41^, that of a previous study ^44^ and the HumanTestisDB ^45^) with whole-body protein-level expression data from the Human Protein Atlas ^46–48^ and find that of thousands of brain-associated proteins, the majority are expressed in male germ cells, with particular enrichment in spermatogonia. Furthermore, we also find the converse to be true: genes enriched in spermatogonia, implicated in human-specific aspects of spermatogenesis, or thought part of the conserved metazoan spermatogenic program, are not only widely expressed throughout the brain but often actively expressed throughout the critical early stages of neurodevelopment, when neurogenesis is at its peak. Finally, we demonstrate that genes with human-specific variants ^49^, which are disproportionately associated with neurodevelopment ^50^, are also enriched in the male germline, with more only detectable at the protein-level in spermatogonia than any other male germ cell type. We contextualise these results with an extensive discussion of the literature and argue that, taken together, they support further enquiry into the testis-brain relationship as a potential source of insight into the evolutionary processes that shaped the human condition. To support this work, we provide extensive datasets of, among others, brain-associated genes and testicular cell markers (**Supplementary Tables 1 to 10**) as resources for the community.

## Materials and Methods

### Human proteomic and single-cell testis and brain transcriptomic data

Our analyses draw on several publicly accessible proteomic and transcriptomic datasets. Firstly, from the Human Protein Atlas (HPA) v23 ^46–48^, we obtained the files ‘normal_tissue.tsv’ and ‘rna_tissue_hpa.tsv’ which represent, respectively, protein-level expression for 56 tissues and 122 distinct cell types (of which 8 cell types were characterised for the testis) and transcript-level expression for 40 tissue types (https://v23.proteinatlas.org/download/normal_tissue.tsv.zip and https://v23.proteinatlas.org/download/rna_tissue_hpa.tsv.zip, downloaded 17^th^ June 2023). Secondly, we obtained an integrated single-cell atlas of 60,427 adult human testicular cells (representing data from 29 individuals across 9 different studies ^51–59^ and an age range of 14-66 years), the bioinformatic methods for which we described in previous work ^41^ and briefly recapitulate here. To create this atlas, we generated cell x gene count matrices using the kallisto/bustools v0.26.3 ‘count’ workflow ^60^, which were then processed using Seurat v4.0.3 ^61^ with SCTransform normalisation ^62^ and integrated using Seurat’s rPCA ‘anchor’ technique. Using an unsupervised clustering approach (the Smart Local Moving algorithm ^63^), we then identified 10 distinct cell clusters, which were annotated using both a global (all-against-all) differential gene expression analysis and established biomarkers for the major somatic and germ cell types (sourced from ^51,53,58,64–67^ and as previously described ^41^). These comprised 3 somatic cell clusters (‘myoid and Leydig cells’, ‘Sertoli cells’, and ‘endothelia and macrophages’, collectively 10,788 cells) and 7 germ cell clusters (‘undifferentiated spermatogonia’ (undiff SPG), ‘differentiating spermatogonia/early meiosis’ (diff SPG), ‘spermatocyte’, ‘early spermatid 1’, ‘early spermatid 2’, ‘late spermatid 1’ and ‘late spermatid 2’, collectively 49,639 germ cells). As a validation that these clusters followed the expected developmental trajectory, the relative proportions of spermatogonia, spermatocytes, and spermatids approximated the ratio 1:2:4. The undiff SPG cluster contained 3298 cells, the spermatocyte cluster 6714 cells, and the four spermatid clusters collectively 36,897 cells (a ratio of 1 to 2.03 to 5.49); note that the germline clusters form a continuum, as shown in **Figure 1B**, and so do not necessarily represent discreet cell types. Differentially expressed genes in each cluster were identified using Seurat’s FindAllMarkers function with parameters min.pct = 0.25, logfc.threshold = 0.25 and only.pos = TRUE which, respectively, require that a gene is expressed in >25% of the cells in a given cluster, that it has a log_2_ fold-change difference in expression relative to all other clusters of > 0.25 (that is, > 2-fold average expression) and that it is on average more highly expressed in that cluster compared to all others.

We also obtained comparable sets of Seurat differential expression analysis results from four previously published human single-cell whole-testis and male-germline atlases, one from a previous study ^44^ (which integrated data from ^51,53,57,58^) and three from the HumanTestisDB ^45^ (which integrated data from ^51,53,56–58,68,69^). Data sources, and details of both the testicular expression atlas contents and analyses performed, are given in **Supplementary Table 4**.

To determine to what extent male germline-associated genes were expressed in the brain both spatially and temporally, we obtained two additional brain transcriptomic datasets. To assess to what extent male germline-associated genes showed preferential expression in particular neural cell types, we used the Karolinska single-cell ‘superset’, a pre-computed cell-type specificity matrix derived from mouse scRNA-seq and comprising 24 ‘level 1’ cell types from the neocortex, hippocampus, hypothalamus, striatum and midbrain ^42^. To assess to what extent male germline-associated genes were expressed throughout the time course of human neurodevelopment, we obtained the Allen BrainSpan Atlas ^43^ Developmental Transcriptome dataset, specifically the file of normalised expression level estimates “RNA-seq Gencode v10 summarised to genes” (https://www.brainspan.org/api/v2/well_known_file_download/267666525, downloaded 26^th^ September 2025). This dataset reports RPKM (reads per kilobase per million mapped reads) values per gene for up to each of 26 brain structures and 31 developmental time points, from embryonic to 40 years. Importantly for our analysis, it provides comprehensive coverage of early human neurodevelopment, including 13 time points from 8 to 37 weeks post-conception. As not all anatomical structures were obtained at all time points, we restricted our analysis to the cerebellar cortex, the most consistently sampled region and, in mammals, the structure containing the vast majority of neurons ^70^.

### Bulk RNA-seq of undifferentiated human spermatogonia

Two previous studies used FACS (fluorescence-activated cell sorting) to purify undifferentiated human spermatogonia from adult testicular biopsies, with a view to enriching their samples for SSCs prior to bulk RNA-seq. We obtained libraries purified using FSD1 ^20^ (four biological replicates with GEO accessions GSM9001401, GSM9001402, GSM9001403, and GSM9001404; 85-88 million 100bp paired-end reads sequenced using an Illumina NovaSeq X Plus) and PLPPR3 ^66^ (four biological replicates with GEO accessions GSM4279699, GSM4279700, GSM4279701, and GSM4279702; 32-35 million 75bp single-end reads sequenced using an Illumina HiSeq 4000). Expression was quantified as transcripts per million (TPM) using Kallisto v0.51.1 ^71^ with the ‘quant’ option and default parameters, and as a transcriptomic index the set of Ensembl v115 ^72^ cDNAs from human genome GRCh38.p14 (http://ftp.ensembl.org/pub/release-115/fasta/homo_sapiens/cdna/Homo_sapiens.GRCh38.cdna.all.fa.gz, accessed 5^th^ May 2026). For the paired-end reads, Kallisto estimates the fragment length from the reads so this parameter does not need to be user-specified. However, for single-end reads, fragment length cannot be empirically derived by the software and is instead assumed to follow a truncated Gaussian distribution with user-specified mean and standard deviation. As such, for these samples, we considered the mean fragment length to be 1.2 × the median read length and the standard deviation to be 0.1 × the median fragment length, obtaining the median read length with fastp v0.23.4 ^73^.

### Protein-coding genes functionally associated with the human brain

To assess the degree of overlap between genes expressed in the testis and functionally associated with the brain, we first compiled an inclusive list of ‘brain genes’ on the basis of either an explicit or suggestive phenotype (**Supplementary Table 1**). This comprised 7193 human brain-associated protein-coding genes (36% of the 19,871 in total in the GRCh38 primary assembly), collated from previous studies of genes associated with macrocephaly ^74^, megalencephaly (i.e. abnormal brain size) ^75^, postmortem brain weight ^76,77^, cognitive ability (using intelligence quotient ^78^ and educational attainment ^79^ as provisional proxies, and defining cognitive ability broadly as the acquisition, retention, processing and use of information ^80^), and human-accelerated regions ^81^ (otherwise conserved genomic loci with human-specific substitutions, the expression of which spatially correlates with patterns of cortical expansion ^50^). We also collated genes from previous reviews of the molecular factors governing brain development ^14^, human cognition ^82^, neocortical evolution ^83,84^, autism spectrum disorder ^85–89^, schizophrenia ^90–92^, genetic epilepsies ^93–96^, and indirect neurogenesis, a process thought to underlie the evolution of the gyrencephalic cortex ^97^.

The overall list is presented in **Supplementary Table 1**, alongside notes on the sources, summary statistics, and the classification of each gene by the nature of its association. As a resource to support further enquiry, genes are assigned, where possible, to one or more of the seven categories of macrocephaly/megalencephaly (n=379 genes), brain weight (n=879), autism (n=2562), schizophrenia (n=345), epilepsy (n=3372), intelligence/educational attainment (n=2303), or human-acceleration (n=1608). We consider the first five of these seven categories to be explicit rather than suggestive phenotypes and to collectively constitute a conservative set of 5212 brain-associated genes.

### Spermatogenesis and human male infertility-associated genes

We compiled a combined, non-redundant, set of 218 genes considered to be either a human spermatogonial marker ^98–100^, to have a human-specific role in spermatogenesis ^101^, or to be a core (conserved) component of the metazoan spermatogenic program ^102^, alongside a set of 154 genes moderately, strongly or definitively linked to a human male infertility phenotype, sourced from two systematic reviews of monogenic gene-disease relationships ^103,104^. Genes, and their sources, are detailed more fully in **Supplementary Table 5**.

### Non-human testicular marker genes

Prospective marker genes for the testicular cell types of humans, 9 non-human mammals (bonobo, chimpanzee, gibbon, gorilla, rhesus macaque, marmoset, mouse, opossum, and platypus) and one bird (chicken) were obtained from a previous study ^105^. Marker genes were considered those differentially expressed in that cell type after snRNA-seq profiling of whole adult testes, as previously described ^105^ (in brief, using Seurat’s FindAllMarkers function ^61^ and retaining only genes with Bonferroni-corrected two-sided Wilcoxon rank-sum test p < 0.05). To ensure a meaningful cross-species comparison of cell types, the authors first grouped the germ cells into 20 evenly distributed bins along the pseudotime trajectory for each species ^105^. Owing to differences in resolution, the cell types for each species were annotated at different levels of granularity. For the seven primates (bonobo, chimpanzee, gibbon, gorilla, human, rhesus macaque, marmoset), 8 germ and 2 somatic cell types were defined: undifferentiated spermatogonia, differentiated spermatogonia, leptotene spermatocytes, pachytene spermatocytes, zygotene spermatocytes, early round spermatids, late round spermatids, elongated spermatids, Sertoli, and ‘other somatic’. By contrast, for four non-primate animals (chicken, mouse, opossum, platypus), only 4 germ and up to 2 somatic cell types were defined, at a lower level of granularity: spermatogonia, spermatocytes, round spermatids, elongated spermatids, Sertoli (platypus excepted), and ‘other somatic’. Accordingly, we processed the ‘7 primate’ and ‘11 animal’ subsets separately. When incorporating the 7 primates into the ‘11 animal’ subset, we renamed their cell clusters for consistency. Thus, a gene differentially expressed in either the early or late round spermatids of gorillas would, when analysed as part of the ‘11 animal’ set, be differentially expressed only in round spermatids.

Finally, to compare gene names across species, we obtained orthology data for each pair of species from Ensembl BioMart v106 ^106^ (i.e. in order to use the same Ensembl stable gene IDs as the source data ^105^). The Ensembl comparative genomics workflow, Compara ^107^, classifies orthologues by type (one-to-one, one-to-many, or many-to-many) and as either high or low confidence on the basis of gene order conservation and pairwise whole genome alignment (detailed further at https://mart.ensembl.org/info/genome/compara/Ortholog_qc_manual.html, accessed 14^th^ June 2026). Using this data, we defined orthogroups as sets of genes whereby every member had a high-confidence one-to-one orthologue with at least one other member, but not necessarily all of them (because for some of the genes in the orthogroup, the orthology relationship could be one-to-many or one-to-one but low-confidence). Orthogroups were named for their human gene name, if a human gene was present and had a name, otherwise they were named for the most commonly occurring gene name within them. Orthogroups and their contents are detailed in **Supplementary Table 8**.

In total, we parsed 219,835 distinct Ensembl IDs across the 11 species, producing 30,964 orthogroups. Of the set of 30,964 orthogroups, 15,269 (49%) had no shortfall between the number of possible high-confidence one-to-one relationships (i.e. number of species represented in the orthogroup x (number of species-1)) and the number observed whereas 15,695 orthogroups (51%) had a shortfall (i.e. difference between maximum number of possible high-confidence one-to-one relationships and the number observed > 0), although as indicated in **Supplementary Table 8** this shortfall was often minimal. A further consequence of this more inclusive definition of orthogroups is that of the total set of 219,835 Ensembl IDs, 846 (0.4%) appeared in more than one orthogroup (often because one or more of the genes had a one-to-many relationship with some of the members). However, in the majority of cases, these orthogroups could be assigned the same gene name regardless (**Supplementary Table 8**). Of these 846 Ensembl IDs, only 185 (22%) appeared in orthogroups with different names; these were excluded from subsequent analyses. Given these considerations, we reasoned that a strict definition of orthogroups, restricted only to high-confidence one-to-one orthologues, would scale poorly with number of species and lead to the unnecessary exclusion of too great a volume of data.

## Results

### Genes associated with brain growth and development are widely expressed in the human male germline

Multiple genes have been functionally implicated with increased brain mass or identified as positively selected along the primate phylogeny or the lineage leading to humans (see reviews ^9,14,83,84^). Given the extensive overlap between testicular and neural transcriptomes ^24^, we hypothesised that a substantial fraction of these brain-growth-associated genes would be active in the male germline. To assess this, we draw upon both the Human Protein Atlas v23 ^46–48^ and an integrated single-cell transcriptomic atlas of 60,427 human testicular cells, which we have previously described ^41^.

We first examined the expression levels of a list of 7193 brain-associated protein-coding genes using the single-cell atlas (**Supplementary Table 1**). This list is intentionally broad with no filters applied beyond those of the original authors (see notes to **Supplementary Table 1**); rather, our aim was simply to determine whether even an unselected list of ‘brain genes’ would show significant overlap with expression in the testis. We summarise the testicular expression of each gene at the protein-level in **Supplementary Table 1** and at the transcript-level in **Supplementary Tables 2 and 3**.

Of the 7193 genes in **Supplementary Table 1**, 2392 (33%) lacked usable protein expression data (either no data or data marked by the HPA as ‘uncertain’) and were excluded from summary statistics. Of the remaining 4801 genes, experimental antibody staining was available; for 2467 (51%) genes staining had been performed for three somatic (peritubular, Leydig, and Sertoli) and five germ cell types, including spermatogonia (detailed further in the notes to **Supplementary Table 1**). For the other 2334 genes (49%), staining was only performed for two cell types: Leydig cells and the broader category of ‘cells in seminiferous ducts’ (that is, seminiferous tubules, comprising all germ cells and Sertoli cells). The HPA only contains the more specific level of antibody staining data (eight cell types, rather than two) for genes with testis-elevated mRNA expression (detailed at https://www.proteinatlas.org/humanproteome/tissue/testis, accessed 12^th^ May 2024). We considered protein expression in seminiferous ducts to represent highly probable germline expression as Sertoli cells, the only somatic cells within them, represent only a few percent of their total ^108^. Although we cannot exclude the possibility that some signals in seminiferous ducts reflect Sertoli cell-specific proteins, this seems unlikely to affect our conclusions as of the 2467 genes stained for eight cell types, only 76 (3.1%) were exclusively detected in Sertoli cells (**Supplementary Table 1**).

Overall, we found that 73% of brain-associated genes had either experimentally-supported or highly probable germline expression (that is, 3512 of 4801 genes for which data was available, irrespective of whether antibody staining was performed only for seminiferous ducts or for specific germ cells; **Supplementary Table 1**). Furthermore, of the 2467 genes where antibody staining data was available for specific germ cells, 2049 brain-associated genes (83%) had detectable protein-level expression in at least one germ cell type, more of which (n = 1476, 59%) were expressed before meiosis (in spermatogonia) than either during (in preleptotene or pachytene spermatocytes; n = 1258, 51%) or after (in round or elongated spermatids; n = 1328, 54%) (**Supplementary Table 1**). The number of brain-associated genes is summarised in **Figure 2A** and the distribution of their proteins in the testis in **Figure 2B**.

**Figure 2.**
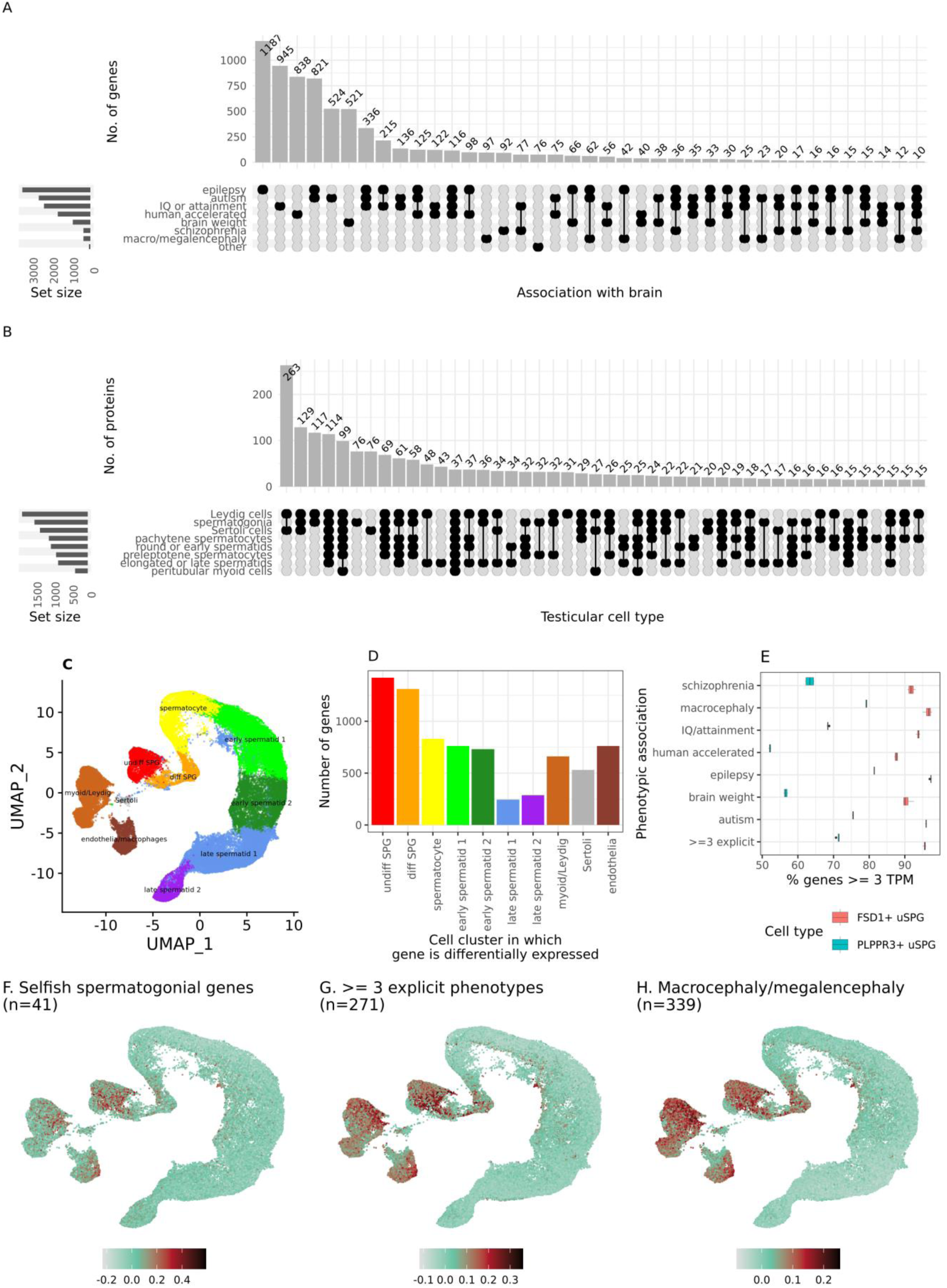
Brain-associated genes and their expression in the human testis. **(A)** Distribution of 7193 genes among an inclusive set of brain-associated phenotypes, both explicit (such as macro- or megalencephaly) and suggestive (such as educational attainment, a proxy for cognitive ability), detailed further in notes to **Supplementary Table 1**. Sets containing < 10 genes are not shown. **(B)** Protein-level expression of 2049 brain-associated genes for which antibody staining data was available for each of 8 testicular cell types (3 somatic, 5 germline) and with a protein detected in at least one (raw data from https://v23.proteinatlas.org/about/download/normal_tissue.tsv, accessed 17^th^ June 2023). Sets represent detectable protein-level expression (i.e. classification into either of the HPA categories of low, medium or high expression) in each cell type (raw data in **Supplementary Table 1**). Sets containing < 15 proteins are not shown. **(C)** Single-cell expression atlas of the adult human testis (n = 60,427 cells), annotated into ten cell clusters (3 somatic, 7 germline). **(D)** Number of genes differentially expressed, at the transcript level, in at least one of these ten clusters (raw data in **Supplementary Tables 2 and 3**). **(E)** Percentage of genes with a brain-associated phenotype detectably expressed (≥ 3 transcripts per million, TPM) in four independent sets of FSD1 or PLPPR3-enriched adult human testicular cells; that is, undifferentiated human spermatogonia. Average expression of genes that **(F)** harbour selfish spermatogonial mutations (n=41), **(G)** have three or more explicit brain phenotypes (n=271), and **(H)**, are associated with either macrocephaly or megalencephaly (n=339), all of which show similarly pronounced expression peaks at the onset of spermatogenesis. A version of panels A to D of this figure restricted to 5212 genes with five explicit brain-associated phenotypes (postmortem brain weight, macrocephaly/megalencephaly, epilepsy, autism, and schizophrenia) is presented as **Supplementary Figure 1** and shows quantitatively similar results.

To further assess whether genes associated with brain growth and development have enriched expression in spermatogonia, we next considered transcript-level expression (**Supplementary Tables 2 and 3**). The testicular transcriptome and proteome are surprisingly dissimilar ^109^: while the testis is abundant in mRNAs, many proteins with high testicular mRNA expression remain undetected by antibody-based or mass spectrometry approaches, with no obvious technical explanation available. One compelling suggestion is that these ‘missing’ proteins are enriched for spermatogenesis-related processes and may have transient, specialised, functions, being rapidly degraded ^109^.

Using data from the testicular single-cell RNA-seq atlas ^41^ (**Figure 2C**), we found that 6293 of the 7193 brain-associated genes (87%) were detectably expressed in the germline (having one or more sequenced reads in >1% of the germ cells; **Supplementary Table 3**). The number of genes differentially expressed in a germ cell cluster declined across the spermatogenic trajectory, being at its highest for ‘undiff SPG’ (1420 genes, or 20% of the total, including *TMEM14B*, a primate-specific gene which promotes cortical expansion ^110^, and both *NOVA1* ^10^ and *TKTL1* ^11^, two genes in which human-specific substitutions have been implicated in neocortical expansion), then decreasing to 1312 (‘diff SPG’), 831 (‘spermatocyte’), 763 (‘early spermatid 1’), 731 (‘early spermatid 2’), 245 (‘late spermatid 1’) and 286 (‘late spermatid 2’) (**Figure 2D** and **Supplementary Table 2**).

Overall, these findings indicate that many genes functionally associated with brain growth are expressed not only in the testis but show particularly enriched expression in undifferentiated spermatogonia, more so than either meiotic or post-meiotic adult germ cells (**Figure 2C**). We note, however, that some of the gene categories in **Figure 2A** represent suggestive associations (such as IQ ^78^ or educational attainment ^79^, broad proxies for cognitive ability) or indirect associations (such as human-accelerated sequence ^81^) with the brain, rather than discreet phenotypes. Nevertheless, repeating the analysis after excluding genes found only in these categories, we obtain quantitively similar results (**Supplementary Figure 1**). Moreover, brain-associated genes were significantly more highly expressed in undifferentiated spermatogonia than all other germ cell types, except spermatocytes (Mann-Whitney U p < 0.05; **Supplementary Figure 2**). To complement this analysis, we also computed the degree to which these genes were expressed across the spermatogenic trajectory using the AddModuleScore function in Seurat ^61^, which calculates the average expression levels of a given gene set for each cell cluster. As expected, the expression of 41 genes known to harbour selfish spermatogonial mutations (14 from ^12,17^ and 27 from ^21^) peaked at the onset of spermatogenesis, in the spermatogonial cell cluster (**Figure 2F**), as did the sets of 271 genes with three or more explicit brain phenotypes (**Figure 2G**) and 339 genes associated with either macrocephaly or megalencephaly (**Figure 2H**).

In terms of single cell expression profiles, the male germline resembles a continuum of ‘states’ (**Figure 2C**) with its division into discrete cell types imposed by a clustering algorithm. As such, our conclusions about the relative enrichment of brain-associated genes across germline populations (i.e. their differential expression; **Figure 2D**) could be sensitive to methodological choices in clustering and annotation. To ensure robustness, we repeated the analysis using comparable Seurat clustering results from four previously published human single-cell whole-testis and male-germline atlases ^44,45^, each of which annotated their cell types independently and varied in the cells used, age range, granularity, and methodology (described further in **Supplementary Table 4**). Across all datasets, we found that brain-associated genes were more frequently differentially expressed in germ than somatic cells (**Supplementary Figures 3 and 4**). Consistent with our own expression atlas ^41^, differentially expressed genes were also particularly abundant towards the start of the spermatogenic trajectory (**Supplementary Figures 3 and 4**). However, unlike our own atlas, which only included adult samples, these alternative datasets also comprised foetal, pre-, and peri-pubertal cells (**Supplementary Table 4**), and hence the highest proportion of differentially expressed genes often appeared in germ cells ‘adjacent’ to spermatogonia, namely leptotene spermatocytes and primordial germ cells (PGCs; **Supplementary Figures 3 and 4**). Notably, PGCs give rise to foetal spermatogonia ^111^ and are transcriptomically similar to ‘state 0’ adult spermatogonia, the most undifferentiated spermatogonial state ^69^.

To test whether our results were driven by specific brain phenotypes, we repeated our analysis for each of the seven phenotypic categories separately. We consistently found differentially expressed genes enriched in the spermatogonia of our own atlas (**Supplementary Figure 5**) and, with few exceptions, around the start of the spermatogenic trajectory in the others (**Supplementary Figures 6, 7, 8 and 9**; the major exceptions are genes associated with human-accelerated regions, which show greater enrichment in spermatids). Finally, to assess whether the observed testis-brain overlap simply reflected a broader enrichment for developmental or housekeeping genes (given their disproportionate association with cell proliferation), we examined, for each phenotypic category, the number of proteins detected in each of the 56 tissues comprising the HPA. As expected, the greatest proportion of each gene (protein) set was expressed in cerebral cortex, cerebellum, and testis (**Figure 3**). This suggests that the overlap is not a generic consequence of ubiquitous protein expression but instead reflects shared molecular programs among these tissues.

**Figure 3.**
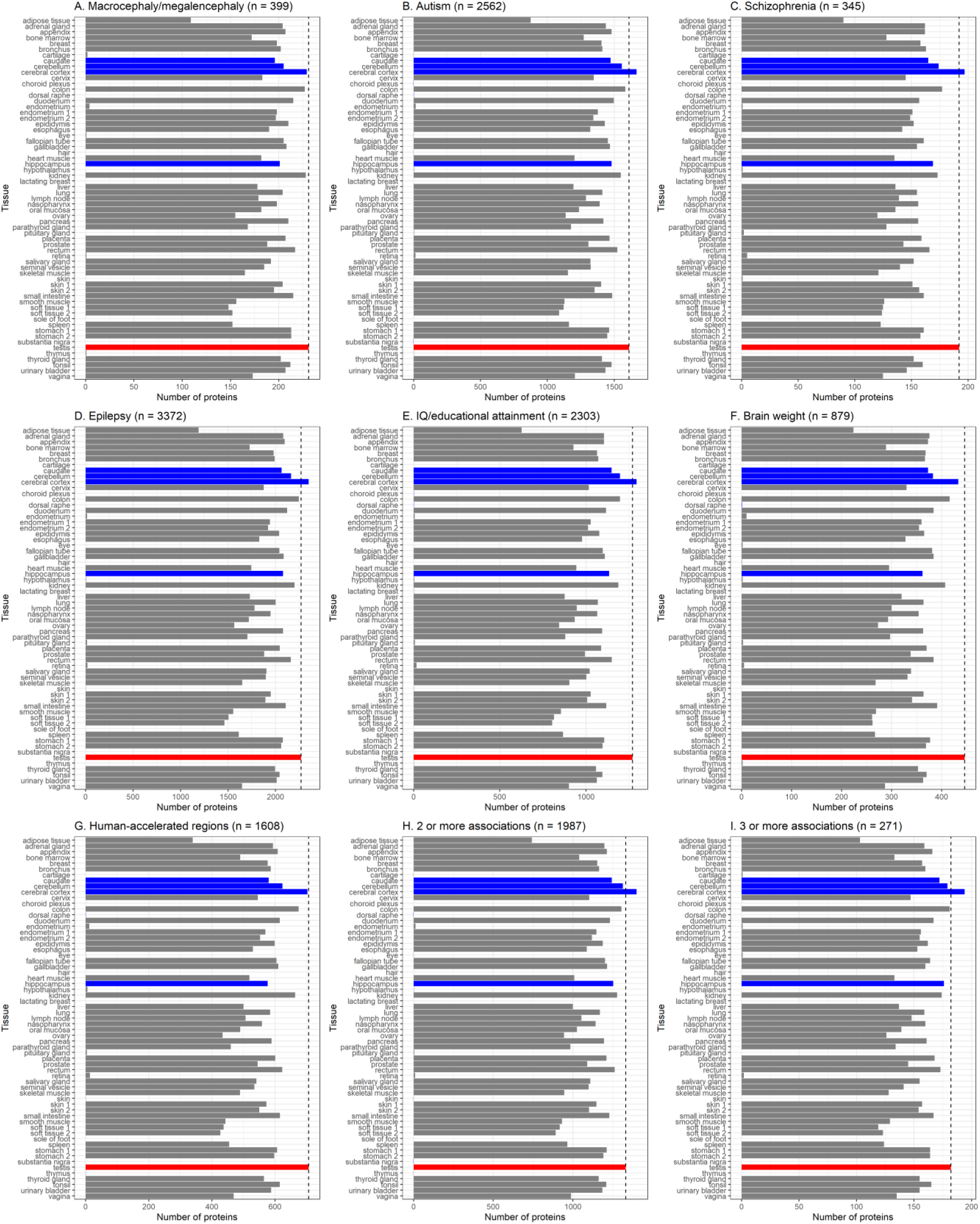
Number of brain-associated genes whose proteins are detectably expressed in human tissues. **(A-G)** Distribution of brain-associated genes for seven different phenotypic categories defined in **Supplementary Table 1**. **(H-I)** Subsets of genes associated with at least 2 (H) or 3 (I) different phenotypic categories, excluding the two ‘indirect’ categories of IQ/attainment and human-acceleration. Raw data for this figure, representing protein expression level per cell type per tissue, was obtained from the Human Protein Atlas v23.0 (https://v23.proteinatlas.org/about/download/normal_tissue.tsv, downloaded 17^th^ June 2023). A protein was considered expressed in a given tissue if, in any cell type of that tissue, ‘level’ was either ‘low’, ‘medium’ or ‘high’ and ‘reliability’ was not ‘uncertain’. Cell types per tissue are detailed in **Supplementary Table 6**. For ease of comparison, brain tissues are shown in blue, testis in red, and all others in grey, with the black dashed line indicating the number of proteins detected in the testis.

Further supporting a bidirectional link between genes functional in the brain and male germline, we also found evidence for the converse: genes considered human spermatogonial markers ^98–100^, having human-specific roles in spermatogenesis ^101^, associated with monogenic male infertility ^103,104^, or thought to be core components of the metazoan spermatogenic program ^102^, not only have broad spatial expression throughout the brain but are often actively expressed throughout the critical early stages of neurodevelopment, when neurogenesis is at its peak (discussed further in the **Supplementary Text** and drawing on analyses of both the ‘Karolinska superset’ of neural cell-type expression data ^42^ and the Allen BrainSpan Atlas Developmental Transcriptome ^43^; see also **Supplementary Tables 5 and 6** and **Supplementary Figures 10, 11, 12, 13, 14 and 15**).

### Genes associated with brain growth and development are abundant in purified human undifferentiated spermatogonia

Thus far, we have used single-cell transcriptomics to show that brain-associated genes are more highly expressed in undifferentiated spermatogonia (uSPG) than all other germ cell types, although it should be noted that the capture rate of scRNA-seq is comparatively low (around 6-30% ^112^), and so in these analyses we were only able to consider a fraction of the total transcriptome. Moreover, the uSPG pool, in which SSCs reside, is characteristically heterogenous such that identifying exclusive SSC cell-surface markers (to enable their isolation and culture for fertility treatments, for instance) has proven challenging ^113^. Nevertheless, both PLPPR3 ^66^ and FSD1 ^20^ have been proposed as ‘most primitive uSPG’ and ‘pan-uSPG’ markers, respectively, and used to produce bulk RNA-seq libraries of the purified human uSPG population. As bulk RNA-seq offers over a hundred-fold higher sequencing depth than scRNA-seq ^20^, we used these libraries to examine the expression of brain-associated genes with greater sensitivity. Corroborating our single-cell results, we found that the majority of each set of brain-associated genes was detectably expressed (≥ 3 transcripts per million, TPM) in purified uSPG, including approx. 80-92% of genes associated with either macrocephaly or megalencephaly (**Figure 2E**).

### Proteins only expressed in the human testis and brain can be associated with brain-expansion phenotypes

The breadth of the HPA antibody staining data offers an additional line of evidence for a functional connection between the male germline and brain: genes whose protein expression is restricted to the two tissues. Although we found only 16 genes out of 5379 (0.3%) whose proteins were only detectable in both a male germ and brain cell, and no other cell type in any other (healthy) tissue (**Supplementary Table 6**), these represent potential entry points for further exploring the molecular parallels between the two organs (discussed further in the **Supplementary Text**). Notably, several of these have been associated with brain-expansion phenotypes. Three have been associated with increased postmortem brain weight ^76^ – *ERMN* (which promotes oligodendroglial differentiation ^114^), *HTR2A* (ectopic expression of which increases the number of basal progenitor cells in embryonic mice ^115^), and *OPALIN* (which promotes oligodendrocyte terminal differentiation ^116^) – one with macrocephaly ^74^ (*HYLS1*, a conserved centriole protein ^117^), and another (*OMG*, an oligodendrocyte glycoprotein which inhibits cell proliferation ^118^) has a co-evolutionary relationship with primate brain mass ^77^ (**Supplementary Table 1**). We note, however, that some of these proteins may also be expressed at low levels or in tissues not assayed by the HPA, and cannot be assumed to be strictly brain-testis specific.

### Proteins with human-specific amino acid changes are simultaneously enriched for neurodevelopmental functions and widely expressed in male germ cells

Having shown that many genes expressed in spermatogonia are functionally associated with the brain, including the brain-expansion phenotypes of macrocephaly and megalencephaly, we next considered whether the germline was also enriched for genes harbouring human-specific variants, as these are disproportionately associated with neurodevelopment ^50^.

Advances in genome sequencing have enabled detailed comparisons of modern humans with archaic hominins, and identified genetic variants that define human-specific traits (see review ^119^). Although the precise catalogue of these variants depends on archaic sample availability, reference genome annotation and coverage, as well as genotype calling and filtering methods, there is nevertheless a broad consensus on the strongest candidates – protein-coding changes in cell cycle-related genes to which the brain is especially sensitive ^49^. A study comparing modern human, Denisovan, and Neanderthal genomes identified 571 genes with non-synonymous amino acid changes in humans (all present at high variant allele frequencies), and an additional 36 genes in which 42 amino acid changes were fixed in modern humans ^49^. Enrichment analyses suggested that these genes mostly affect cell division and early brain growth trajectories, with notable expression in the infant frontal cortex ^49^. However, this same gene set also shows a strong connection to the testis: it is significantly enriched for the GO term ‘spermatoproteasome complex’. Furthermore, 3 of the 42 fixed amino acid changes occur in *SPAG5* (sperm-associated antigen 5), a gene crucial for meiosis and spermatid morphogenesis ^120^ (yet dispensable for fertility ^121^) and which is also expressed in the foetal ventricular zone during cortical neurogenesis ^122^. *SPAG5* encodes a mitotic spindle protein, with mutant phenotypes consistent with a critical role in whether a cell divides symmetrically or asymmetrically ^119^. This is particularly relevant to NSPCs as the orientation of the mitotic cleavage plane can impact cell fate decisions, and thereby the composition of the progenitor pool ^123^.

Motivated by these observations, we hypothesised that genes containing human-specific amino acid changes would also be detectably expressed (and putatively functional) in the male germline, and spermatogonia in particular. To test this, we assessed protein-level expression in the male germline for 309 of the 607 genes in the human-specific substitution set (genes with no usable data were excluded; **Supplementary Table 7**). Of these, 70% had either experimentally-supported or highly probable (that is, in seminiferous ducts) germline expression. Furthermore, of the 152 genes where protein expression data was available for five germ cell types, 22 were only detectable in spermatogonia (compared to 4 in spermatocytes and 15 in spermatids). These include *ASCC1*, mutations in which downregulate genes associated with neurogenesis and neuronal migration ^124^, *LRRTM1*, an imprinted gene expressed throughout forebrain development and associated paternally with cerebral asymmetry ^125^, and *TCF3*, which represses neuronal differentiation and increases the self-renewal of NSPCs during neocortical development ^126^.

### Genes associated with brain growth and development are also enriched in the male germline of other metazoans

The majority of our results have centred on humans, due to the wealth of data available for our species. Nevertheless, our hypothesis that many genetic variants contributing to brain size expansion first arose in spermatogonia predicts a general principle and so we also expect to see an enrichment of brain-associated genes in the spermatogonia of other mammals. To assess this, we obtained data from a previous study using whole-testes snRNA-seq profiling to identify prospective markers for the testicular cell types of humans, 9 non-human mammals (bonobo, chimpanzee, gibbon, gorilla, rhesus macaque, marmoset, mouse, opossum, and platypus) and one bird (chicken) ^105^. To ensure meaningful cross-species comparisons, we reprocessed this data to produce both ‘7 primate’ and ‘11 animal’ subsets, each with a common set of cell types, and ensured each species’ set of marker genes had a common name derived on the basis of orthology relationships (see Methods). The total number of marker genes per cell type per species are shown in **Supplementary Figures 16 and 17** for the primate and animal sets, respectively. These figures show that although many marker genes are species-specific there is nevertheless a substantial overlap between them, likely reflecting components of the conserved (ancestral) spermatogenic program (orthogroups are defined in **Supplementary Table 8** with their differential expression in each testicular cell type per species detailed in **Supplementary Table 9**). This ancestral program comprises genes differentially expressed in the same cell type in multiple species, and notably includes those in which selfish spermatogonial mutations have been characterised, such as *ARID1B* (differentially expressed in spermatogonia in 9 species of the ‘11 animal’ dataset; **Supplementary Table 9**), *DYRK1A*, *MAP2K1*, *NF1*, *SOS1* and *PACS1* (each differentially expressed in uSPG in 5 species of the ‘7 primate’ dataset; **Supplementary Table 9**).

We next considered to what extent the number of brain-associated genes differentially expressed in each metazoan germ cell cluster declined across the spermatogenic trajectory, an analysis comparable to that shown only for humans in **Figure 2C**. As expected, for both primate and animal subsets, the number of brain-associated genes typically peaked at the start of the trajectory, in the uSPG or spermatogonia clusters, respectively. This pattern was more apparent for primates, especially human, and consistent across both inclusive and exclusive sets of brain-associated genes: the inclusive set of 5212 genes with either of five explicit brain-associated phenotypes (**Supplementary Figure 18**) and the exclusive sets of 271 genes with three or more explicit brain phenotypes (**Figure 4**) and 339 genes associated with either macrocephaly or megalencephaly (**Supplementary Figure 20**). Nevertheless, unlike our analyses with scRNA-seq data, we also found a high enrichment of brain- and macrocephaly-associated genes in Sertoli cells, at levels comparable to or higher than that in spermatogonia. The reasons for this are unclear, although may in part be technical (unlike snRNA-seq, scRNA-seq requires tissue dissociation which disproportionately loses somatic cells) or reflect the fact that Sertoli cells are the active coordinators of spermatogenesis and, indeed, the only somatic population synchronized with it ^127^.

**Figure 4.**
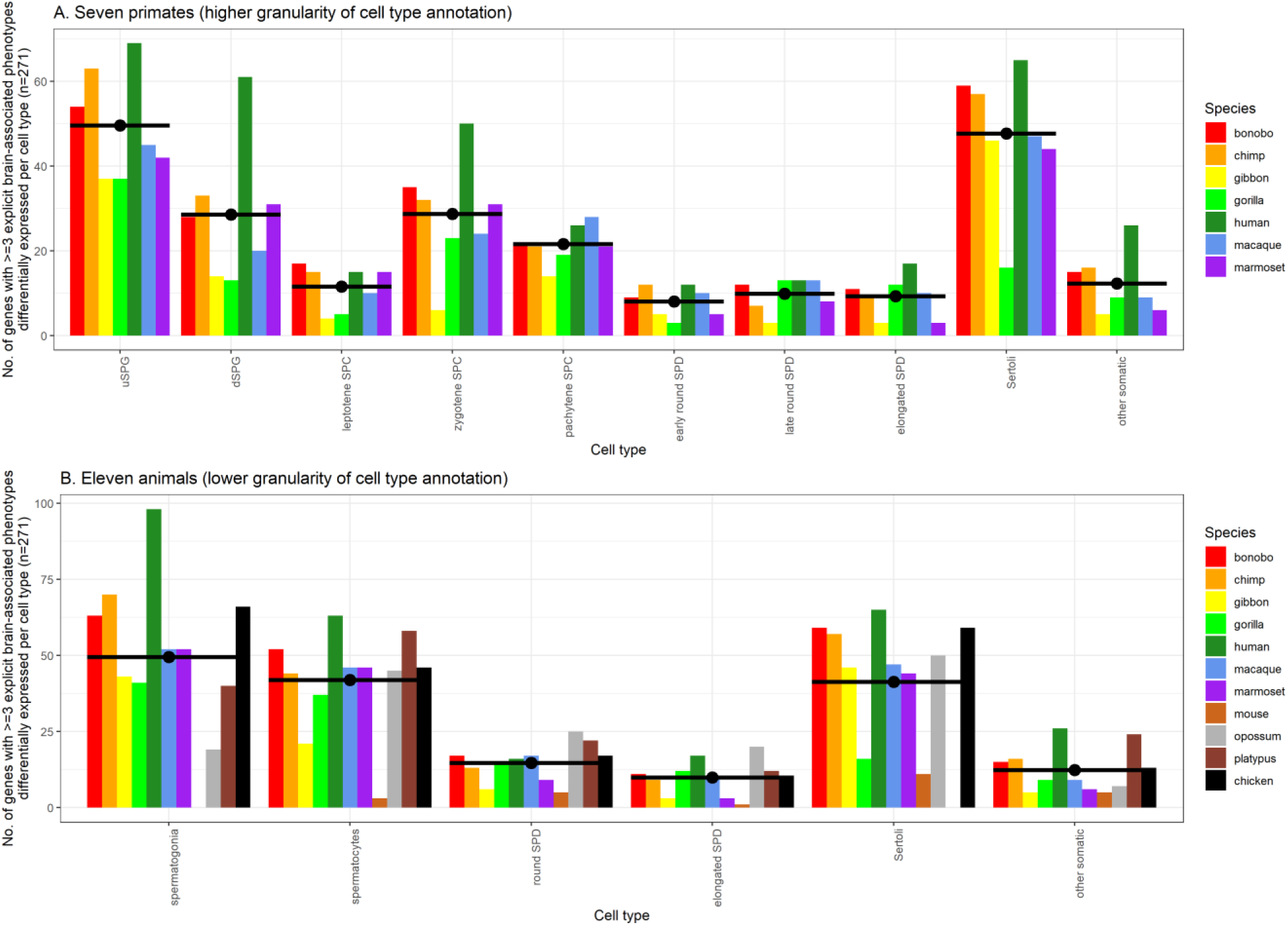
Number of genes with ≥ 3 explicit brain-associated phenotypes that are also testicular cell type markers in (A) 7 primate, and (B) 11 animal species. This figure represents a re-analysis of data from ^105^. To compare gene names across species, we only plotted those that could be assigned to an orthogroup, defined as a set of genes whereby every member had a high-confidence one-to-one orthologue with at least one other member. Orthogroups are defined in **Supplementary Table 8** with their differential expression in each testicular cell type per species detailed in **Supplementary Table 9**. The contents of the brain-associated gene set are detailed in **Supplementary Table 1**. Black bars for each group denote the mean number of genes across species. uSPG, undifferentiated spermatogonia; dSPG, differentiated spermatogonia; SPC, spermatocyte; SPD, spermatid.

## Discussion

We have previously hypothesised that the dysregulation of male germline homeostasis provides a novel perspective on the mechanisms underlying mammalian brain expansion ^13^. In the present work we further test the validity of this proposal by systematically examining the extent to which genes with functional roles in brain growth, development or evolution are also expressed in the male germline, and vice versa. In conjunction with numerous observations from the literature, we summarise our results, alongside other evidence supporting our hypothesis, in **Table 1**.

**Table 1.**
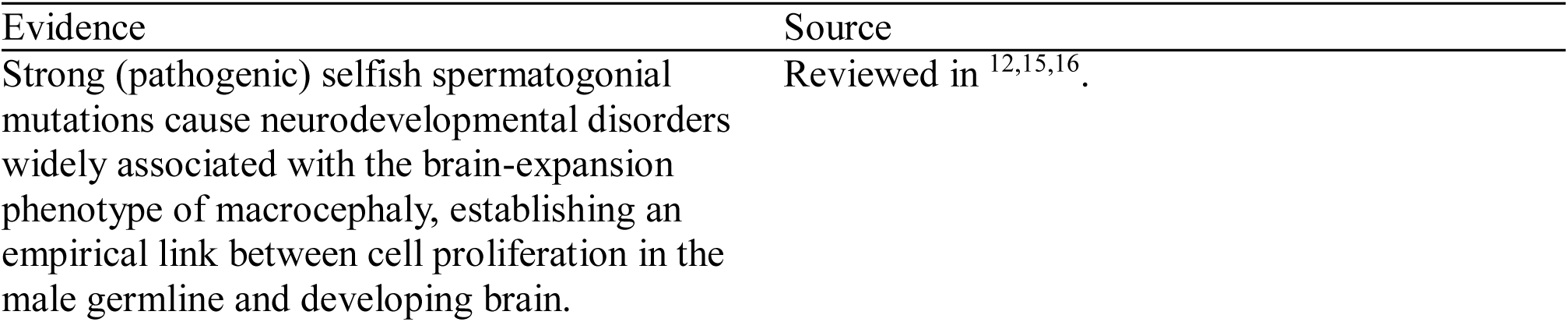
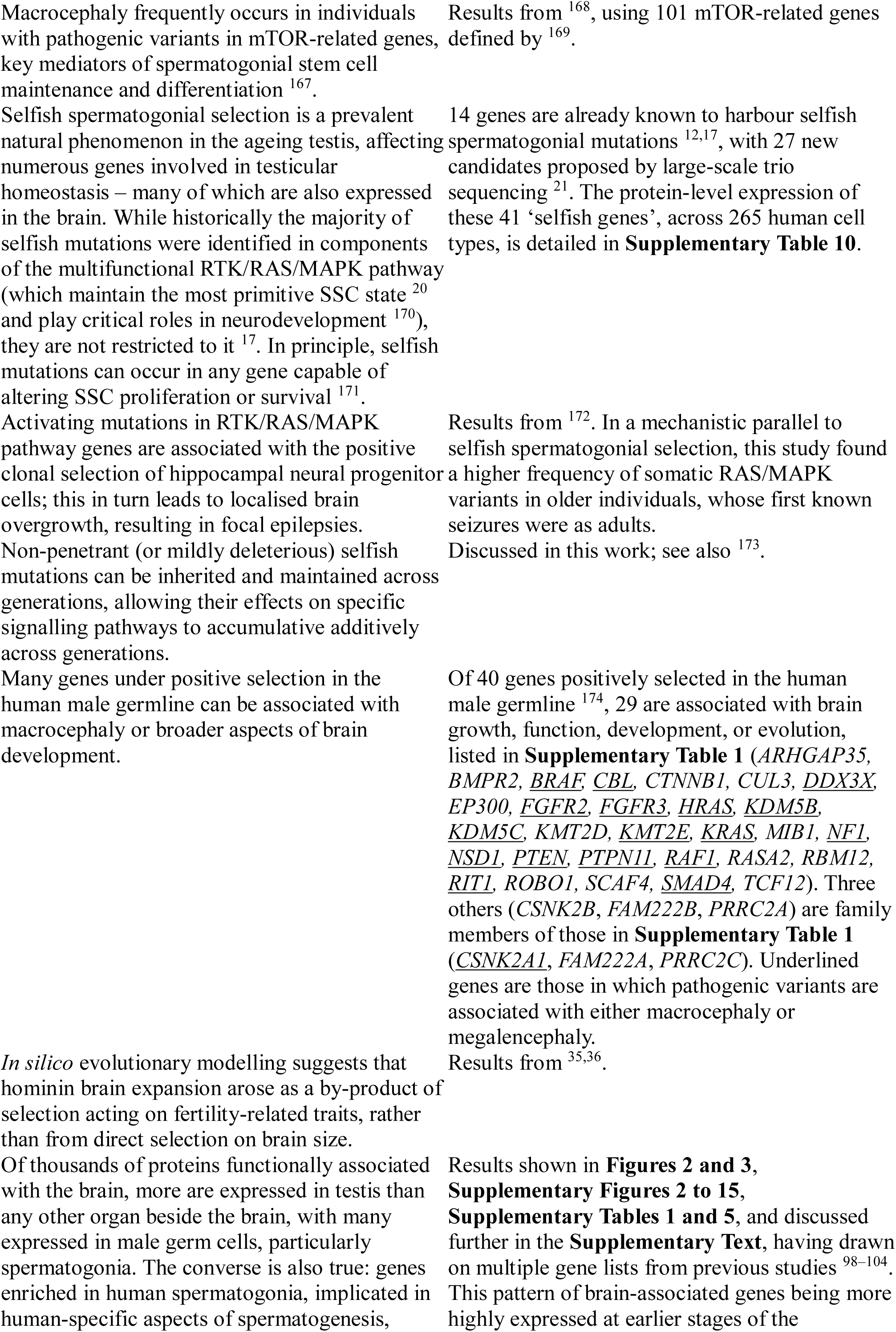
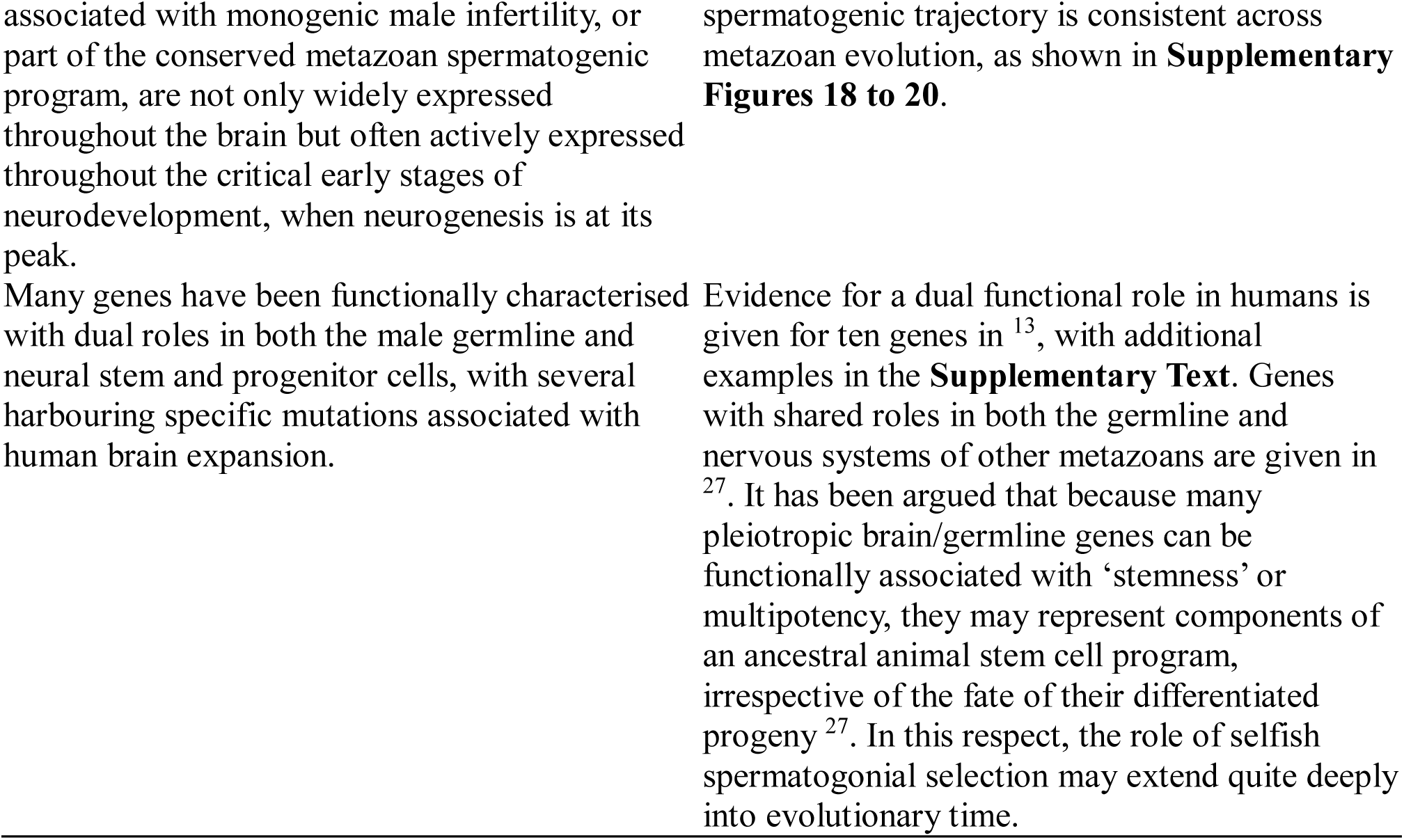
Overview of the lines of evidence implicating a link between the male germline, and spermatogonia in particular, and brain expansion.

Our entryway into this topic was the identification of a mechanism of enrichment of spontaneous oncogenic-like mutations in spermatogonial stem cells, in genes that are functionally critical to brain development. This mechanism, selfish selection, provides a causal link between the otherwise disparate fields of spermatogenesis, neurogenesis, and oncogenesis, and offers a novel perspective on a number of observations. For example, a genome-wide association study of the genetic variants associated with human head size ^128^ (a highly heritable trait and strong correlate of brain size) identified 67 candidate loci. These loci were disproportionately located near genes preferentially expressed in intermediate neural progenitors, whose increased proliferation has been linked to human brain expansion ^129^, as well as being enriched for pathways involved in both macrocephaly and cancer. Consistent with our hypothesis, many proteins encoded by these genes are also abundant in spermatogonia and have functional roles in male fertility, including FOXO3 ^130^, HMGA2 ^131^, IGF2BP1 ^132^, NFIX ^133^, and PPP6R3 ^134^ (**Supplementary Table 1**).

Similarly, another genome-wide association study of human brain MRI data ^135^ identified 199 loci significantly associated with either cortical surface area or thickness, traits linked to the NSPC proliferation rate and their number of neurogenic divisions, respectively. Many of these loci lie in genes involved with the establishment and maintenance of the male germline, or in regions functionally linked to Wnt signalling, a key regulator of spermatogenic cell fate ^136^. Examples include variants in *PAX7* (a marker of spermatogonial stem cells in mice ^137^), *MOV10* (knockdown of which affects spermatogonial cell fate decisions in mice ^138^), *DAAM1* (which regulates the actin cytoskeleton of sperm ^139^), and *HDAC9* (which, in chickens, promotes the differentiation of embryonic stem cells into male germ cells ^140^).

Our hypothesis of a testis-brain relationship also intersects with the biology of human-accelerated regions (HARs), many of which act as neurodevelopmental enhancers ^141^. The evolutionary changes underpinning human cognitive abilities have been conceptualised as “Achilles’ heels” as they may predispose to the high burden of psychiatric disorders in our species – that is, they may represent maladaptive by-products of adaptations in brain development ^142^. Supporting this view, many HARs with enhancer activity in NSPCs show evidence of compensatory evolution to maintain ancestral activity. That is, variants may have initially induced large shifts in regulatory activity in the brain which were then moderated by nearby substitutions ^142^. Although the underlying forces responsible for accelerated substitution rates in HARs remain poorly understood, this ‘back and forth’ pattern is compatible with a scenario in which variants were initially adaptive in another context (i.e. male germline) and only indirectly affected the brain, with phenotypic consequences that may be mildly deleterious but tolerated.

As discussed above, these testis-brain associations reflect the shared developmental and regulatory origins of the neural lineage and the germline. In humans, this coordination centres on the early specification of primordial germ cells (PGCs), mediated by the core transcriptional network of *OCT4* (*POU5F1*), *PAX5,* and *PRDM1* (*BLIMP1*). This regulatory ‘master switch’ simultaneously activates germline and represses somatic differentiation programs, thereby committing cells to a germline fate (reviewed in ^98,143^), the establishment and maintenance of which requires extensive chromatin modelling ^144^. Disruption of this process can lead to reprogramming, with PGCs aberrantly expressing somatic genes. Tellingly, when germ cells reprogram as soma in both *C. elegans* ^145^ and mice ^146^, they often express pan-neuronal markers, extrude neurite-like projections, and differentiate into neurons. It has been suggested that, in metazoans in general, the default differentiation program for cells that lose their germ line identity is neuronal ^147^.

Although our focus here has been on the conceptual links between SSCs and NSPCs, the genetic architecture of brain size is multifaceted and complex ^148^. It includes, among others, gene duplication ^149,150^, gene family size variation ^151–153^, and changes to coding ^154,155^, non-coding ^156^, and regulatory ^157^ sequences, these mechanisms collectively giving rise to relatively large brains through multiple independent evolutionary trajectories ^158^. Moreover, larger brains are not only defined by a greater number of neurons: they also differ in neuronal size, regional specialisation, and packing density, as well as their number and proportion of somatic support cells (glia) ^159^. Importantly, not all genes implicated in brain size expansion are functionally active in the testis as the overlap between testis and brain proteomes, although substantial, is not exhaustive. For example, human-specific *NOTCH2NL* genes, which arose from recent gene duplication events to play critical roles in cortical neurogenesis ^160^, have no known association with spermatogenesis. Taken together, we have drawn upon the biology of the testis to offer a fresh perspective on an old question: how did the human brain become so large? Looking through the lens of the germline can provide new insight into this fundamental question. A male germline-specific evolutionary force, selfish selection, enriches genomes for oncogenic-like mutations in pathways involved in the regulation of spermatogonial stem cells – and which are also functional in neural stem and progenitor cells. In this respect, we can conceive of the male germline as both a generator of mutations and a filter, disproportionately passing to the next generation those of a particular (‘selfish’) nature – ones benefiting reproductive success and robust sperm production whilst ageing. It follows that if selfish mutations disproportionately accumulate within pathways essential for SSC proliferation and fertility, they may also influence the phenotype of other (somatic) cells in which the same genes are expressed. Scaled up to the population level, the long-term effect of this additive, non-random, germline mutagenesis may have been to expose to selection variants that modulate baseline rates of cell division, potentially exerting second-order effects in related lineages, such as neural progenitors (consistent with this, protein-coding changes in genes involved in cell cycling rank among the strongest candidates for human-specific sequence variation ^49^). This would also align with current theories of human brain evolution through the concept of bradychrony (‘slowed time’), which refers to the relative slowing of equivalent developmental processes between species ^161^. Notably, differences in developmental tempo correlate with differences in brain size. In humans, brain maturation is markedly protracted (bradychronic) compared to other primates, taking a longer proportion of the lifespan to reach adult brain weight ^161^. Early human neurodevelopment is particularly prolonged, with the period of tangential cortical expansion, driven by symmetrically-dividing radial glial cells, lasting approximately a month in humans but a fortnight in macaques ^161^. The mechanisms driving differences in neurodevelopmental tempo are an area of active enquiry with calcium regulation (a process also fundamental to both SSC regulation ^162,163^ and sperm function ^24^) emerging as a key candidate ^164^. Enrichment of variants selfishly selected in the testis could also contribute by subtly modulating stem cell turnover rates, thereby influencing the duration and extent of neurogenic output. Although in support of this hypothesis we have summarised a large body of work on male germline mutations as they relate to brain expansion, experimental work to establish a causal relationship will be necessary. Future work could consider to what extent genes and mutations associated with sperm production rate (such as number or turnover rate of spermatogonial stem cells) are associated with brain phenotypes. This is an underexplored research area, with few obvious models in which to address it, although some tentative directions are discussed in the **Supplementary Text**.

To conclude, we highlight a pronounced, yet largely unexplored, role for the male germline – and spermatogonia in particular – in shaping both the development and evolution of the brain. Beyond demonstrating a substantial overlap between genes expressed in spermatogonia and those functionally associated with neurodevelopment, we draw conceptual parallels between the neural and spermatogonial stem cell lineages noting that the timing and balance of self-renewing and differentiating cell divisions are not only critical to both but especially so for longer-lived species (such as humans), who reproduce over many decades. Variants selfishly selected in the testis may bias this balance towards increasing cellular output (i.e. more sperm and/or neurons), thereby influencing both fertility and cognitive ability. These variants, in affecting neuron production, may have been especially influential throughout vertebrate evolutionary history as the strongest neuroanatomical correlate of their cognitive abilities is, to date, absolute cortical neuron number ^165,166^. We hope our analyses will encourage further enquiry into this topic as we find it particularly compelling to consider how mutations selectively advantageous to human spermatogonia may have been co-opted by other tissues, contributing to the genomic innovations underlying our unusually large, neuron-rich, brains.

## Authors’ contributions

**Stephen J. Bush**: conceptualisation, methodology, formal analysis, data curation, investigation, validation, visualisation, software, writing – original draft, writing – review & editing.

**Anne Goriely**: conceptualisation, validation, funding acquisition, writing – review & editing.

## Data availability

The Seurat object for our integrated single-cell atlas of 60,427 adult human testicular cells, described in ^41^, is available from FigShare at https://doi.org/10.6084/m9.figshare.31443727. All other data supporting this work is available either in the article itself or in its electronic supplementary material.

## Code availability

All scripts used to perform the analyses and create both figures and tables are available at www.github.com/sjbush/testis_brain.

## Supplementary Figures

**Supplementary Figure 1. A conservative set of brain-associated genes and their expression in the human testis.**

**(A)** Distribution of 5212 genes among an explicit set of brain-associated phenotypes, detailed further in notes to **Supplementary Table 1**. Sets containing < 10 genes are not shown. **(B)** Single-cell expression atlas of the adult human testis (n = 60,427 cells), annotated into ten cell clusters (3 somatic, 7 germline). **(C)** Number of genes differentially expressed, at the transcript level, in at least one of these ten clusters (raw data in **Supplementary Tables 2 and 3**). **(D)** Protein-level expression of 1573 brain-associated genes for which antibody staining data was available for each of 8 testicular cell types (3 somatic, 5 germline) and with a protein detected in at least one (raw data from https://v23.proteinatlas.org/about/download/normal_tissue.tsv, accessed 17^th^ June 2023). Sets represent detectable protein-level expression (i.e. classification into either of the HPA categories of low, medium or high expression) in each cell type (raw data in **Supplementary Table 1**). Sets containing < 15 proteins are not shown.

**Supplementary Figure 2. Average expression across, respectively, an (A) inclusive and (B) conservative set of 7193 and 5212 brain-associated genes for each of ten cell clusters in an adult human testis expression atlas.**

The brain-associated gene sets are detailed in **Supplementary Table 1** with expression level per testicular cell type in **Supplementary Table 2**. Statistical comparisons are Mann-Whitney U tests, implemented using the R package ‘ggpubr.’

**Supplementary Figure 3. Number of differentially expressed brain-associated genes per cell cluster, for each of four testis and male germline single-cell atlases, and using an inclusive set of brain-associated genes.**

For each of the four single-cell atlases, barplots show the number of genes significantly differentially expressed within an inclusive set of brain-associated genes (n=7193). The cell atlases and their respective annotations are described in **Supplementary Table 4**, alongside the differential expression analysis results, which were filtered according to a common set of criteria. The brain-associated gene sets are detailed in **Supplementary Table 1**.

**Supplementary Figure 4. Number of differentially expressed brain-associated genes per cell cluster, for each of four testis and male germline single-cell atlases, and using a conservative set of brain-associated genes.**

For each of the four single-cell atlases, barplots show the number of genes significantly differentially expressed within an inclusive set of brain-associated genes (n=5212). The cell atlases and their respective annotations are described in **Supplementary Table 4**, alongside the differential expression analysis results, which were filtered according to a common set of criteria. The brain-associated gene sets are detailed in **Supplementary Table 1**.

**Supplementary Figure 5. Number of brain-associated genes differentially expressed, at the transcript level, in one or more of ten testicular cell clusters.**

The list of brain-associated genes is given in **Supplementary Table 1**, with raw data for this figure in **Supplementary Tables 2 and 3**. Panels A to G show the distribution of differentially expressed genes for seven different categories of phenotype, whereas panels H and I require that each gene is associated with at least 2 or 3 different categories, respectively, not including the two ‘indirect’ phenotypes of IQ/attainment and human-acceleration.

**Supplementary Figure 6. Number of brain-associated genes differentially expressed, at the transcript level, in one or more of cell clusters from the ‘Salehi 2023’ single-cell expression atlas.**

The ‘Salehi 2023’ expression atlas, its respective annotations, and the differential expression analysis results used for this figure, are detailed in **Supplementary Table 4**. The brain-associated gene sets are detailed in **Supplementary Table 1**. Panels A to G show the distribution of differentially expressed genes for seven different categories of phenotype, whereas panels H and I require that each gene is associated with at least 2 or 3 different categories, respectively, not including the two ‘indirect’ phenotypes of IQ/attainment and human-acceleration.

**Supplementary Figure 7. Number of brain-associated genes differentially expressed, at the transcript level, in one or more of cell clusters from the ‘Wang 2025 (all cells)’ single-cell expression atlas.**

The ‘Wang 2025 (all cells)’ expression atlas, its respective annotations, and the differential expression analysis results used for this figure, are detailed in **Supplementary Table 4**. The brain-associated gene sets are detailed in **Supplementary Table 1**. Panels A to G show the distribution of differentially expressed genes for seven different categories of phenotype, whereas panels H and I require that each gene is associated with at least 2 or 3 different categories, respectively, not including the two ‘indirect’ phenotypes of IQ/attainment and human-acceleration.

**Supplementary Figure 8. Number of brain-associated genes differentially expressed, at the transcript level, in one or more of cell clusters from the ‘Wang 2025 (germ cells)’ single-cell expression atlas.**

The ‘Wang 2025 (germ cells)’ expression atlas, its respective annotations, and the differential expression analysis results used for this figure, are detailed in **Supplementary Table 4**. The brain-associated gene sets are detailed in **Supplementary Table 1**. Panels A to G show the distribution of differentially expressed genes for seven different categories of phenotype, whereas panels H and I require that each gene is associated with at least 2 or 3 different categories, respectively, not including the two ‘indirect’ phenotypes of IQ/attainment and human-acceleration.

**Supplementary Figure 9. Number of brain-associated genes differentially expressed, at the transcript level, in one or more of cell clusters from the ‘Wang 2025 (germ cells – part 1)’ single-cell expression atlas.**

The ‘Wang 2025 (germ cells – part 1)’ expression atlas, its respective annotations, and the differential expression analysis results used for this figure, are detailed in **Supplementary Table 4**. The brain-associated gene sets are detailed in **Supplementary Table 1**. Panels A to G show the distribution of differentially expressed genes for seven different categories of phenotype, whereas panels H and I require that each gene is associated with at least 2 or 3 different categories, respectively, not including the two ‘indirect’ phenotypes of IQ/attainment and human-acceleration.

**Supplementary Figure 10. Beeswarm (violin scatter) plot of the cellular specificity, in the mouse brain, of 14 genes known to harbour selfish spermatogonial mutations.**

Cellular specificity values, from the Karolinska superset ^42^, represent the proportion of the total expression of a gene attributable to one of 24 brain cell types (a specificity of 0 means that the gene is not expressed in that cell type and a value of 1 that it is only expressed in that cell type). The set of 14 genes known to harbour one or more selfish spermatogonial mutations have been identified in humans ^12,17^ although for the purpose of visualisation here, specificity data from the one-to-one mouse orthologue is used.

**Supplementary Figure 11. Violin plot of the cellular specificity of 218 spermatogonia and spermatogenesis-associated genes in the mouse brain.**

Cellular specificity values, from the Karolinska superset ^42^, represent the proportion of the total expression of a gene attributable to one of 24 brain cell types (a specificity of 0 means that the gene is not expressed in that cell type and a value of 1 that it is only expressed in that cell type). Violins are sorted from left to right in ascending order of median specificity. The genes used for this plot are listed in **Supplementary Table 5** and drawn from previous studies ^98–102^. The genes in this table are human and for the purpose of visualisation here, the one-to-one mouse orthologue (if extant) is used.

**Supplementary Figure 12. Violin plot of the cellular specificity of 154 male infertility-associated genes in the mouse brain.**

Cellular specificity values, from the Karolinska superset ^42^, represent the proportion of the total expression of a gene attributable to one of 24 brain cell types (a specificity of 0 means that the gene is not expressed in that cell type and a value of 1 that it is only expressed in that cell type). Violins are sorted from left to right in ascending order of median specificity. The genes used for this plot are listed in **Supplementary Table 5** and drawn from two previous systematic reviews ^103,104^. The genes in this table are human and for the purpose of visualisation here, the one-to-one mouse orthologue (if extant) is used.

**Supplementary Figure 13. Boxplot of the expression in the cerebellar cortex over time of 14 genes known to harbour selfish spermatogonial mutations, using the Allen BrainSpan Atlas.**

The set of 14 genes known to harbour one or more selfish spermatogonial mutations have been described in previous studies ^12,17^.

**Supplementary Figure 14. Violin plot of the expression in the cerebellar cortex over time of 218 spermatogonia and spermatogenesis-associated genes, using the Allen BrainSpan Atlas.**

The genes used for this plot are listed in **Supplementary Table 5** and drawn from previous studies^98–102^.

**Supplementary Figure 15. Violin plot of the expression in the cerebellar cortex over time of 154 male infertility-associated genes, using the Allen BrainSpan Atlas.**

The genes used for this plot are listed in **Supplementary Table 5** and drawn from two previous systematic reviews ^103,104^.

**Supplementary Figure 16. Degree of overlap between marker genes for each of 9 testicular cell types in each of 7 primate species.**

This figure represents a re-analysis of data from ^105^. To compare gene names across species, we only plotted those that could be assigned to an orthogroup, defined as a set of genes whereby every member had a high-confidence one-to-one orthologue with at least one other member. Orthogroups are defined in **Supplementary Table 8** with their differential expression in each testicular cell type per species detailed in **Supplementary Table 9**. uSPG, undifferentiated spermatogonia; dSPG, differentiated spermatogonia; SPC, spermatocyte; SPD, spermatid.

**Supplementary Figure 17. Degree of overlap between marker genes for each of 6 testicular cell types in each of 11 animal species.**

This figure represents a re-analysis of data from ^105^. To compare gene names across species, we only plotted those that could be assigned to an orthogroup, defined as a set of genes whereby every member had a high-confidence one-to-one orthologue with at least one other member. Orthogroups are defined in **Supplementary Table 8** with their differential expression in each testicular cell type per species detailed in **Supplementary Table 9**. SPD, spermatid.

**Supplementary Figure 18. Number of brain-associated genes that are also testicular cell type markers in (A) 7 primate, and (B) 11 animal species, using an inclusive set of 5212 brain-associated genes.**

This figure represents a re-analysis of data from ^105^. To compare gene names across species, we only plotted those that could be assigned to an orthogroup, defined as a set of genes whereby every member had a high-confidence one-to-one orthologue with at least one other member. Orthogroups are defined in **Supplementary Table 8** with their differential expression in each testicular cell type per species detailed in **Supplementary Table 9**. The contents of the brain-associated gene set are detailed in **Supplementary Table 1**. Black bars for each group denote the mean number of genes across species. uSPG, undifferentiated spermatogonia; dSPG, differentiated spermatogonia; SPC, spermatocyte; SPD, spermatid.

**Supplementary Figure 19. Number of macrocephaly/megalencephaly-associated genes that are also testicular cell type markers in (A) 7 primate, and (B) 11 animal species.**

This figure represents a re-analysis of data from ^105^. To compare gene names across species, we only plotted those that could be assigned to an orthogroup, defined as a set of genes whereby every member had a high-confidence one-to-one orthologue with at least one other member. Orthogroups are defined in **Supplementary Table 8** with their differential expression in each testicular cell type per species detailed in **Supplementary Table 9**. The contents of the gene set are detailed in **Supplementary Table 1**. Black bars for each group denote the mean number of genes across species. uSPG, undifferentiated spermatogonia; dSPG, differentiated spermatogonia; SPC, spermatocyte; SPD, spermatid.

## Supplementary Tables

**Supplementary Table 1.** Summary of the testicular expression, at the protein-level, of 7193 genes associated with brain growth, function, development, or evolution.

**Supplementary Table 2.** Transcript-level expression of genes associated with brain growth, function, development, or evolution, using an adult human testis single-cell expression atlas.

**Supplementary Table 3.** Summary of the testicular expression, at the transcript-level, of genes associated with brain growth, function, development, or evolution.

**Supplementary Table 4.** Sources of cluster (cell type) annotations, and Seurat differential gene expression analysis results, for four previously published single-cell human testis atlases.

**Supplementary Table 5.** Transcript-level expression in the brain of spermatogonia-enriched genes, human-specific germline-associated genes, genes associated with human male infertility, and core components of the metazoan spermatogenic program.

**Supplementary Table 6.** Proportion of male germline-expressed proteins detectably expressed in other tissues.

**Supplementary Table 7.** Summary of the testicular expression, at the protein-level, of protein-coding genes containing human-specific high-frequency missense changes.

**Supplementary Table 8.** Gene orthogroups across 11 metazoans.

**Supplementary Table 9.** Testicular cell marker genes in metazoan testis single-nucleus expression atlases (adapted from data originally published by Murat, *et al.* 2023).

**Supplementary Table 10.** Protein-level expression, across 265 human cell types, of 41 genes in which selfish spermatogonial mutations have been characterised.

