## Supplementary Text and Figures S1 to S19 for "A spermatogonial perspective on the expansion of the mammalian brain"

**This file includes:**

Supplementary Text

Supplementary Figures 1 to 19

References

**Other supplementary materials for this manuscript include:**

Supplementary Tables 1 to 10

**SUPPLEMENTARY TEXT**

**(A) Selfish genes and their role in neural stem and progenitor cell (NSPC) regulation**

In the main text, we wrote that: *“These selfish mutations are consistently associated with severe conditions in the offspring […] This is plausibly linked to the roles of many of these genes in regulating cell fate decisions during neurogenesis (discussed further in the Supplementary Text)”.* We elaborate on this point here.

Many of the genes expressed in undifferentiated spermatogonia and known to harbour selfish mutations (namely, *BRAF*, *CBL*, *FGFR2*, *FGFR3*, *HRAS*, *KRAS*, *MAP2K1*, *MAP2K2*, *NRAS*, *PTPN11*, *RAF1*, *RET*, *SMAD4* and *SOS1* ^1–3^) belong to or intersect with the RTK/RAS/MAPK signalling cascade and play roles in proliferation and cell fate decisions during neurogenesis. Given their ability to promote clonal expansion in the testis, functional mutations in these genes could also exert substantial downstream effects on brain size by altering neurogenic output. This can be demonstrated by experimentally manipulating these genes and observing the impact upon phenotype. For example, *in vitro* cell counting experiments in mice show that conditional activation of *KRAS* increases neural stem cell proliferation rate, without altering cell fate decisions ^4^. Constitutive activation of *MAP2K1* also increases the production of radial glial cells (RGCs) – a major class of neural stem cells – in the outer subventricular zone, a cortical germinal layer and key neurogenic niche ^5^. Ventricular injection of FGF2 similarly increases the growth of the rostral ventricular zone and number of progenitors within it, and correlates with cortical folding ^6^; conversely, *FGF2* knockout almost halves the number of cortical neurons by the end of the neurogenic period ^7^. In addition, mice engineered with gain-of-function mutations in the terminal MAPK gene, *Mapk1* (encoding ERK2, which by phosphorylation directly regulates the expression of many genes ^8^), recapitulate the neurodevelopmental features of human RASopathies ^9^. These gain-of-function mutations, equivalent to *de novo* human missense variants, enhance ERK2 signalling and are thought to disrupt the balance between self-renewing (proliferative) and asymmetric (neurogenic) progenitor divisions, producing an increased number of oligodendrocyte precursors ^9^. Similarly, ERK2-knockout mice present with microcephaly, the result of an elongated neural progenitor cell cycle that shifts the balance towards neurogenic division, thereby prematurely depleting the progenitor pool and affecting the overall number and distribution of pyramidal neurons ^10^.

In rats, co-activation of *FGFR1* and *FGFR3* promotes symmetric division of RGCs, while their inactivation shifts the balance towards asymmetric division and differentiation, ultimately depleting the RGC pool and resulting in a smaller cortex ^11,12^. Furthermore, in ferrets, a dominant-negative mutant allele of *FGFR3*, when expressed at a critical developmental timepoint (embryonic day 33), alters the relative proportions of RGCs leading to a reduction in cortical folding ^13^. By contrast, the induced expression of a constitutively active *FGFR3* mutant allele (p.K664E) in mice did not affect cortical folding, although did increase cortical thickness ^14^. Indeed, the mutational spectrum of *FGFR3* is considered by ^15^ to be “probably the strongest genetic evidence” supporting the hypothesis that mammalian cortical expansion was driven largely by FGF/RAS/MAPK signalling in RGCs. The author proposed that the activation of MAPK (ERK) signalling caused by *FGFR3* gain-of-function induces *BMP7* and *GLI3R* expression, the latter acting to repress *SHH*, a known antagonist of ERK ^15,16^. This establishes a positive feedback loop that sustains ERK activity during development, thereby extending the period of neural progenitor proliferation and increasing the size of the cortical NSPC pool through enhanced self-renewal ^15,16^. It was also proposed that in smaller-brained species such as mice (approx. 14 million cortical neurons), higher concentrations of SHH in the cerebrospinal fluid intrinsically suppress ERK signalling, so maintaining a smaller cortical progenitor pool. By contrast, larger-brained species such as humans (approx. 16 billion cortical neurons), by virtue of being larger, have naturally lower SHH concentrations, which enables prolonged FGF/MAPK/ERK activity that may increase brain size further ^15,16^. This hypothesised mechanism of cortical expansion over evolutionary time is predicated on the nature, strength, and frequency of gain-of-function mutation(s) in FGF signalling components and/or their downstream effectors, particularly those in the RAS/MAPK pathway, in which selfish spermatogonial mutations have been abundantly documented ^1^.

**(B) Many cell markers used to phenotype spermatogonia can be functionally associated with neurogenesis and brain expansion.**

In the main text, we wrote that: *“many characteristic spermatogonial markers also play functional roles in neurogenesis and brain expansion (see also Supplementary Text)”.* We elaborate on this point here.

With advances in single-cell RNA sequencing technology, many studies have characterised the expression profiles of testicular cells across the human and mouse lifespan, both in health and disease (see review ^17^). These studies have yielded a broadly accepted set of markers for distinguishing cell types along the spermatogenic trajectory (reviewed in ^18^). Notably, a number of the most widely used spermatogonial markers, including *FGFR3*, play critical roles in neurodevelopment. For example, *ZBTB16* (*PLZF*) is essential for spermatogonial stem cell (SSC) self-renewal ^19,20^ but has also been associated with increased encephalisation quotients (a measure of relative brain size) across the primate phylogeny ^21^ and directly activates *PAX6*, a transcription factor central to multiple aspects of mammalian cortical development ^22^. Of note, *PAX6* is highly expressed in the radial glial cells (RGCs, the NSPCs of the cortex ^23,24^) of primates compared to mice, and that its overexpression in the mouse embryonic neocortex increases RGC proliferation rate, and thereby overall number ^12^. In addition, *PAX6* activates *TBR2* (*EOMES*), a marker of both mouse spermatogonia and the intermediate progenitors that drive indirect neurogenesis in amniotes ^12^. Lineage tracing studies in mice indicate that a subset of EOMES-expressing spermatogonia are long-term self-renewing SSCs ^25^, highlighting the requirement – shared with the neurogenic lineage – to tightly balance proliferation and self-renewal to avoid exhaustion.

This convergence is not limited to a handful of well-characterised genes. Many commonly used markers of undifferentiated human spermatogonia (from a recent review ^18^) can be functionally associated with neurodevelopmental or neuronal functions. Examples include *EGR4*, which regulates cell-fate decisions in geniculate ganglion neurons ^26^; *GFRA1*, required for synaptogenesis ^27^; *PIWIL4*, which modules neurogenic gene expression programs ^28^; and *TSPAN33*, which regulates the metalloprotease *ADAM10* ^29^, itself central to neuronal differentiation ^30^*.* Other markers such as *MAGEA4* and *UTF1* are ‘cancer/testis genes’, better known for their overexpression in the context of neural tumours such as glioblastomas ^31^, germinomas ^32^ and neuroblastomas ^33^. These links highlight that the gene markers used to define undifferentiated spermatogonia overlap extensively with those implicated in neuronal lineage specification and maintenance.

SSCs constitute a rare subpopulation within the undifferentiated spermatogonial compartment ^34^, estimated to represent only approx. 35,000 cells per adult mouse testis, equivalent to 1.25% of all spermatogonia, and just 0.03% of all germ cells ^35^. Although genes exclusive to SSCs have not yet been identified, there is nevertheless a broad consensus on those enriched in SSCs and critical to their self-renewal and maintenance. These genes include the transcription factors *BCL6B*, *ETV5*, *ID4*, *LHX1*, *NANOS2*, *PDX1*, and *TBR2* (*EOMES*), as well as the surface markers *GFRA1* and *THY1* ^34^. Many of these genes are also associated with neural development or in neural stem cell dynamics, including *ETV5*, which promotes glial progenitor specification in the developing brain ^36^; *ID4*, which regulates neural progenitor proliferation ^37^ and quiescence in adult mouse hippocampal stem cells ^38^; *LHX1*, which synchronises neuronal activity underlying circadian rhythms ^39^; and *PDX1*, which regulates calcium homeostasis in sensory neurons ^40^. In addition, associations with expression profiles are also noticeable, such as for *BCL6B*, which has a multi-regional signature of sex-biased expression in the human brain ^41^, and *THY1*, which is a marker of oligodendrocyte precursors in fetal human brains ^42^. Again, these overlaps are not just anecdotal, as many of these genes act as core components of stem cell maintenance programs in both the germline and nervous system.

We have previously described a set of genes whose protein products are abundant in spermatogonia and which also contribute to brain development, namely *BNIP3*, *CTNNB1*, *DNMT1*, *EZH2*, *FMR1*, *FXR1*, *MCPH1*, *NSD1*, *OGA*, *PAFAH1B1*, *PPP1R17*, and *TKTL1* ^43^. Beyond these, many other spermatogonial proteins play established roles in the brain, a selection of which is given in the table below.

Although the following examples are illustrative of this general point and have not been selected systematically, the broader testis-brain overlap can be evaluated quantitatively. To that end, using the Human Protein Atlas v23 ^44^, we identified 5380 proteins for which expression level was assessed for eight testicular cell types (not counting those where the reliability of the antibody profiling was ‘uncertain’). Of those proteins only detected in one of the eight cell types, the majority were unique to spermatogonia (n=184, or 3.4% of the total); for context, this was followed by Sertoli cells (n=150) and elongated spermatids (n=134), with the fewest proteins unique to preleptotene spermatocytes (n=9). Focusing on these 184 spermatogonia-specific proteins (which are plausible candidates for selfish spermatogonial selection), we note that a substantial proportion are associated with neurodevelopmental phenotypes: indeed, 76 of the 184 genes (41%) are listed in **Supplementary Table 1**, compared to 43 of the 134 elongated spermatid-specific genes (32%). In the table below, we mark genes with an asterisk (*) when their protein expression is confined to spermatogonia and no other of eight testicular cell types, and with a dagger (†) when their protein expression is confined to spermatogonia and no other of five germ cell types (but still detectable in one or more testicular somatic cell types).

| Gene/protein | Evidence supporting a functional role in the brain |
| --- | --- |
| AAK1 * | Serine/threonine protein kinase ubiquitously expressed in the central nervous system, participating in multiple nervous system-related signalling pathways, and required for neuropathic pain responses ^45^. |
| ADD3 | Cytoskeletal protein with tumour suppressive properties, the loss of which promotes cell proliferation in glioblastoma multiforme ^46^. |
| ADNP | Transcription factor, the haploinsufficiency of which causes Helsmoortel-Van der Aa syndrome, a common form of syndromic autism ^47^. Promotes neuronal differentiation via modulation of Wnt signalling ^48^; supports neurite outgrowth and maturation ^49^ and neuronal survival ^50^. Enriched for recurrent *de novo* mutations associated with neurodevelopmental disorders ^51^. |
| ALKBH5 | RNA demethylase widely expressed throughout the developing mouse brain, predominantly in NeuN-expressing neurons ^52^. Influences cell proliferation and differentiation in the cerebellum by modulating levels of RNA m6A methylation in cell fate determination genes ^53^. Mutations in *ALKBH5* are associated with increased testis size across tetrapod evolution ^54^, with *Alkbh5* a leading QTL candidate for testis weights in mice ^55^ (accordingly, *Alkbh5* knockout mice present impaired fertility and reduced testis size ^56^). |
| ANKS1B * | Encodes AIDA-1, a synapse-enriched protein which regulates synaptic plasticity ^57^. Heterozygous microdeletions in *ANKS1B* result in a broad spectrum of neurodevelopmental phenotypes ^58^. |
| AUTS2 | Regulates the cell fate of human cortical progenitors and promotes cerebral cortex expansion ^59^. |
| BAZ1B † | Chromatin modifier regulating a number of neural crest-associated genes showing human-specific signs of selection ^60^. Although knockout also results in ectopic epigenetic signals during meiosis, BAZ1B is dispensable for male fertility in mice ^61^. |
| CAPRIN1 | Regulates proliferation and migration in multiple cell types, including neurons; haploinsufficiency results in neuronal loss and a neurodevelopmental disorder characterised by language impairment, attention deficit hyperactivity, and autism ^62^. |
| CNTN1 * | Encodes contactin-1, a neuronal cell surface receptor involved in the formation and maintenance of perineuronal nets, key regulators of neural plasticity ^63^. |
| CRLF3 | Evolutionarily ancient gene which emerged with the metazoan nervous system ^64^ as a central regulator of neuron differentiation, maturation and survival ^65^. |
| DYRK1A | Haploinsufficiency results in microcephaly in both mice ^66^ and humans ^67^, whereas, in humans, gain-of-function increases brain volume ^66^. Enriched for *de novo* pathogenic variants in a large rare-disease cohort, consistent with clonal expansions in spermatogonia ^3^. |
| FOXP1 † | Critical regulator of self-renewing cell divisions in both apical ^68^ and basal ^69^ radial glia, and widely associated with autism spectrum disorders ^70^. |
| KDM6B | Histone demethylase acting on the promoters of neuron-specific genes in mice and required for neuronal differentiation ^71^. Haploinsufficiency produces autistic-like behavioural traits in mice ^72^ and conditional knockout reduced synaptic activity ^73^. Enriched for *de novo* variants in large independent cohorts of individuals with neurodevelopmental disorders ^74^ and autism spectrum disorder ^75^. Pathogenic *KDM6B* variants result in complex phenotypes, although developmental delay and cognitive defects are the most consistent clinical features ^76^. |
| MACROD2 | Expressed in cortical and hippocampal neurons in the developing mouse brain ^77^. Human intragenic deletions cause multiple congenital anomalies including microcephaly and intellectual disability ^78^. Identified as the top genome-wide significant locus for temporal lobe volume in a neuroimaging GWAS ^79^. |
| MTOR | Heterozygous mutations cause Smith-Kingsmore syndrome, characterized by macrocephaly or megalencephaly ^80^. Enriched for *de novo* gain-of-function mutations in a large rare-disease cohort, consistent with clonal expansions in spermatogonia ^3^. |
| PARD6A | Core member of an apical protein complex which plays a critical role in cell polarity ^81^ and in radial glial cell divisions: overexpression biases divisions towards self-renewal while knockdown promotes neurogenic differentiation ^81^. |
| PENK * | Neuropeptide gene marking a neuronal subpopulation constituting the most prominent component of a long-term memory engram in mice ^82^. |
| POGZ | Cerebrocortical and hippocampal neuron-enriched chromatin regulator ^83^ promoting the transcription of synaptic genes during early development ^84^; conditional knockout in mice causes microcephaly ^85^. In humans, *de novo* mutations are associated with developmental delay and microcephaly ^86^. |
| PPP1CB | Heterozygous mutations cause a Noonan syndrome-like disorder, characterized by relative or absolute macrocephaly ^87^. Component of the RTK/RAS/MAPK signalling pathway, a central regulator of testicular homeostasis ^88,89^, and enriched for *de novo* gain-of-function mutations in a large rare-disease cohort, consistent with clonal expansions in spermatogonia ^3^. |
| RBFOX2 | Splicing regulator required both for murine cerebellar development and the maintenance of mature neuron physiology ^90^, as well as spermatogonial stem cell differentiation and meiotic initiation ^91^. |
| STAG1 | Regulates the G2/M transition and apoptosis of murine hippocampal NSPCs in response to *Gli1* activation, expression of which induces cell cycle arrest as a protective mechanism against hyperproliferation ^92^. *De novo* heterozygous mutations in humans cause intellectual disability, epilepsy, and microcephaly ^93^. |
| STIP1 * | Highly expressed in glioblastomas; downregulation in glioma cells slows proliferation by lengthening G1 phase ^94^. |
| TLE1 | Transcription factor promoting glioblastoma growth by supporting the maintenance of stem-like tumour-initiating cells in the brain ^95^. |
| TRIO | Essential for late embryonic mouse development including neural tissue organization ^96^. Highly expressed in glioblastomas ^97^; depletion reduces glioma cell proliferation rate ^98^. |
| VIM | Encodes vimentin, a cytoskeletal filament protein involved in cell migration, shape, division, and plasticity ^99^, as well as protecting differentiating stem cells from stress by directing the asymmetric partitioning of stress granules during mitosis ^100^. Implicated in multiple nervous system disorders ^101^. Expressed in neural stem and progenitor cells, where mutations at serine sites phosphorylated during mitosis result in increased levels of neuronal differentiation ^102^. |

**Genes with protein-level expression in either spermatogonia or the seminiferous ducts (assessed by antibody profiling) and with functional associations with neuronal proliferation, differentiation, or neurodevelopment in general.**

According to the Human Protein Atlas ^44^, each gene shows protein-level expression either specifically in spermatogonia or, more broadly, in seminiferous ducts (see https://www.proteinatlas.org/humanproteome/tissue/testis). Genes marked * were exclusively detected in spermatogonia and no other of eight testicular cell types; genes marked † could only be detected in spermatogonia and no other of five germ cell types. For the vast majority of entries, protein-level expression was explicitly detected in spermatogonia. The three exceptions are *ALKBH5*, *MTOR*, and *RBFOX2*, for which the Human Protein Atlas only assessed expression for ‘cells in seminiferous ducts’ (raw data available in **Supplementary Table 1**). Nevertheless, there is abundant support for spermatogonial expression from other species: immunofluorescent staining shows intense ALKBH5 immunoreactivity in the spermatogonia, primary spermatocytes and round spermatids of cattle and yak testis ^103^, and functional experiments in mice show that *Mtor* knockout impairs spermatogonial proliferation ^104^, and that *Rbfox2* knockdown inhibits meiotic initiation ^91,105^.

**(C) Genes enriched in spermatogonia, involved in human-specific aspects of spermatogenesis, associated with human male infertility, or representing components of the core metazoan spermatogenic program, are also widely expressed throughout the brain.**

In the main text, we wrote that: *“Further supporting a bidirectional link between genes functional in the brain and male germline, we also found evidence for the converse: genes considered human spermatogonial markers, having human-specific roles in spermatogenesis, associated with monogenic male infertility, or thought to be core components of the metazoan spermatogenic program, not only have broad spatial expression throughout the brain but are often actively expressed throughout the critical early stages of neurodevelopment, when neurogenesis is at its peak (discussed further in the Supplementary Text)”.* We elaborate on this point here.

To complement our observation that many genes influencing brain growth, function and evolution are detectable in spermatogonia (**Supplementary Table 1**), we used data from the Human Protein Atlas (HPA) to evaluate whether genes enriched in the male germline are similarly expressed in the brain. To do so, we first obtained a set of 44 genes only detected at the protein level in human spermatogonia, and no other germ cell type ^106^. 35 of these genes (80%) were expressed at the mRNA level in at least one of 13 brain regions, including 27 (61%) transcribed in all of them (**Supplementary Table 5**; anatomical regions are described in more detail at https://www.proteinatlas.org/humanproteome/brain, accessed 15^th^ May 2024). Notably, this set of 44 genes includes *DMRT1*, *FGFR3*, *GFRA1*, *ID4*, *KIT*, *MAGEA4, NANOS2*, *NANOS3*, *PAX7, SALL4*, *UCHL1*, *UTF1* and *ZBTB16*, each of which are widely used undifferentiated, differentiating, and differentiated spermatogonial biomarkers in single-cell transcriptomic studies ^18,107^, alongside *EXOSC10*, a marker of undifferentiated spermatogonia in humans ^108^. With the exceptions of *DMRT1*, *MAGEA4* and *NANOS2*, all of these genes are expressed in at least one brain brain, and most were expressed in all of them (**Supplementary Table 5**).

We next supplemented this set of spermatogonia-enriched genes in two ways. First we added 4 additional spermatogonial markers compiled by ^107^ (*MKI67*, *RET*, *SOHLH1*, and *STRA8*). Second, we incorporated 44 transcription factors whose expression was enriched in spermatogonia at any point in human development (from embryogenesis to adulthood) ^109^; these lists draw from multiple single-cell transcriptomic studies ^110–117^ and are used non-redundantly here. Consistent with the above finding, 46 of these genes (96%) were transcribed in at least one brain region and 39 (81%) in all of them (**Supplementary Table 5**).

In addition, we considered a distinct set of genes proposed to underlie human-specific roles in spermatogenesis. A recent study integrating epigenetic and expression data identified 23 genes with germ cell-restricted activity in humans, and which were absent from comparable rhesus macaque, mouse, and opossum datasets ^118^. Of these 23 genes, 17 (74%) were transcribed in at least one brain region, and 14 (61%) in every region (**Supplementary Table 5**).

Finally, another recent study performed network analysis on the spermatocyte transcriptomes of human, mice and flies to identify 104 genes comprising the putative core metazoan spermatogenic program ^119^. Of note, surprisingly, this gene set shows almost no overlap with any of the aforementioned gene sets with one exception, the transcription factor *CLOCK*, considered by ^118^ to have a human-specific role in spermatogenesis. Strikingly, despite this minimal overlap, 96 of these genes (92%) were expressed in all regions of the human brain (**Supplementary Table 5**), with 31 (30%) also directly associated with brain growth or development (that is, it also appears in **Supplementary Table 1**).

Aggregating these datasets yields a total of 218 genes enriched in human spermatogonia or otherwise implicated in either the metazoan or human-specific spermatogenic program (i.e., the combined gene list from ^106,107,109,118,119^). Overall, we found that 198 of these genes (91%) had detectable mRNA expression in at least one region of the human brain and 175 (80%) in all regions (**Supplementary Table 5**). Complementing these lists of spermatogenesis-associated genes, we also assessed the brain expression of 154 genes with moderate, strong or definitive evidence for association with monogenic male infertility phenotypes, compiled from two systematic reviews ^120,121^ (**Supplementary Table 5**). Here again the overlap with brain expression was extensive: 141 of these genes (92%) had detectable mRNA expression in the human brain, with 102 (66%) detected in every region. Taken together, the HPA expression data consistently shows that genes associated with spermatogenesis and male fertility are broadly expressed across the brain.

To assess whether these germline-associated genes show preferential expression in particular neural cell types, we used the Karolinska single-cell ‘superset’, a pre-computed cell-type specificity matrix derived from mouse scRNA-seq and comprising 24 ‘level 1’ cell types from the neocortex, hippocampus, hypothalamus, striatum and midbrain ^122^. Its original purpose was to sample a broad range of brain regions relevant to the neurobiology of schizophrenia and accordingly, its authors note relative sparsity of coverage for cortical and striatal development (that is, under-representation from gestation, early post-natal, and adolescent periods) ^122^. Furthermore, although this dataset has been widely used to implicate certain cell types as enriched for a given (human) gene list (for instance, by ^123^), there remain cells for which data is unavailable or (despite the generally high degree of mouse-human conservation for genes expressed in brain) where the genes differ in function or selective pressure between the two species. These caveats in mind, we plotted the distribution of cell-type specificity scores for (i) the set of 14 genes known to harbour selfish spermatogonial mutations ^1,2^ (**Supplementary Figure 10**), (ii) the combined sets of 218 spermatogonia and spermatogenesis genes (from ^106,107,109,118,119^; **Supplementary Figure 11**), and (iii) 154 male infertility genes (from ^120,121^; **Supplementary Figure 12**).

Overall, the specificity distributions were broad, consistent with widespread expression across cell types rather than concentration in particular neural progenitor classes. There is a notable exception in **Supplementary Figure 10**, however: *Fgfr3* expression was more specific to radial glial-like cells (specificity = 0.275; discussed in section A of this supplementary text) and ependymal astrocytes (a cell type which can reversibly switch from a mature to a stem/progenitor-like identity ^124^; specificity = 0.350). Only one gene showed any substantial degree of specificity to a differentiated population: *Ret*, to adult dopaminergic neurons (specificity = 0.753). Otherwise, genes with selfish mutations showed no consistent enrichment in neural progenitors; instead, consistent with their fundamental roles in regulating cell proliferation, these genes were expressed ubiquitously across neural cell types.

Having established that spermatogenesis and male fertility-associated genes (**Supplementary Table 5**) exhibit broad spatial expression throughout the brain, we next asked whether these genes were also broadly expressed across the temporal course of neurodevelopment. Under our proposed hypothesis of a germline-brain relationship, we expect that these genes – irrespective of whether they have documented roles in neural proliferation, or harbour selfish mutations – should at least be detectably expressed throughout the critical early stages of neurodevelopment.

To assess this, we obtained the Allen BrainSpan Atlas ^125^ Developmental Transcriptome dataset, specifically the file of normalised expression level estimates “RNA-seq Gencode v10 summarised to genes” (https://www.brainspan.org/api/v2/well_known_file_download/267666525, downloaded 26^th^ September 2025). This dataset reports RPKM (reads per kilobase per million mapped reads) values per gene for up to each of 26 brain structures and 31 developmental time points, from embryonic to 40 years. Importantly, it provides comprehensive coverage of early neurodevelopment, including 13 time points from 8 to 37 weeks post-conception. As not all anatomical structures were obtained at all time points, we restricted our analysis to the cerebellar cortex, the most consistently sampled region and, in mammals, the structure containing the vast majority of neurons ^126^. This allowed us to ask whether germline-associated genes show sustained transcription during these periods of active neurogenesis.

For each of our three gene sets (14 genes with selfish spermatogonial mutations, 218 spermatogenesis-associated genes, and 154 male infertility-associated genes), we calculated the mean RPKM across the cerebellar cortex samples (**Supplementary Figures 13, 14, and 15**, respectively). As anticipated, all known selfish genes were detectably expressed (RPKM > 1) throughout embryonic development of the cerebellar cortex, often markedly so (RPKM > 10), as were a clear majority of the spermatogonia- and spermatogenesis-associated gene set. In contrast, many genes associated with adult male infertility – an inherently heterogenous category, including genes not necessarily directly linked to spermatogenesis ^120,121^ – showed minimal expression in the cerebellar cortex, with a median RPKM approximating 1.

Taken together, these observations indicate a consistent transcriptomic association between the male germline and brain development, although it remains important to emphasise that transcription alone is not indicative of function and these correlations should not be interpreted as direct evidence of shared mechanistic roles. It is also worth noting that alternative splicing, widespread in both the male germline and brain ^127^, may mitigate pleiotropic effects for genes transcribed in both organs. In principle, distinct splice isoforms could support organ-specific roles, allowing a gene to perform different cellular roles without functional conflict.

We next considered the extent to which protein-level expression was shared between cells of the brain and the testis. The Human Protein Atlas (HPA) dataset contains expression profiles for the protein products of 11,167 genes in 144 cell types from 63 tissues based on immunohistochemistry (although not every protein was assessed in each cell type; data from https://v23.proteinatlas.org/about/download/normal_tissue.tsv, accessed 17^th^ June 2023). We excluded all data with a reliability score of ‘uncertain’ (https://www.proteinatlas.org/about/assays+annotation#ih_reliability). Our initial aim was to determine the proportion of genes whose proteins were detectable in one of five germ cell types (as discussed in the main text) and which were also found in other tissues.

As expected, the strongest overlap was between protein expression in male germ cells and the brain. Of the 4543 genes with detectable expression in at least one germ cell type, 3589 (79%) were also found in at least one of the seven brain regions surveyed by HPA (caudate, cerebellum, cerebral cortex, choroid plexus, dorsal raphe, hippocampus, or substantia nigra; note that the HPA immunohistochemistry dataset covers a different set of brain regions as those assessed for mRNA expression); 3461 of these were expressed in the cerebral cortex (**Supplementary Table 6**). We note, however, that this is a restricted subset of the spermatogonial proteome and that accordingly there were also substantial overlaps with other tissues (78% of the 4543 germline proteins could also be found in colon, for instance; **Supplementary Table 6**). Nevertheless, this observation supports the transcriptomic data (**Supplementary Table 5**) and indicates that the proteins encoded by ‘spermatogenesis genes’ (more specifically, those enriched in spermatogonia ^106,107,109^, involved in human-specific roles in spermatogenesis ^118^, or part of the conserved metazoan spermatogenic program ^119^), are widely expressed throughout the brain.

Finally, we have also noted in the main text that brain-associated genes (listed in **Supplementary Table 1**) are typically transcribed earlier in the spermatogenic trajectory (illustrated in **Figure 2C**). Corroborating this, among the genes with detectable protein expression in both the male germline and brain, a higher proportion were expressed in spermatogonia than in either spermatocytes or spermatids. For instance, 2611 of 3589 proteins (58%) are found in both brain cells and spermatogonia compared to 1519 (33%) found in both brain cells and round spermatids (**Supplementary Table 6**). This enrichment in the earliest germ-cell population is consistent with the broader argument that gene programs operating during early neurodevelopment and early spermatogenesis overlap to a substantial extent.

**(D) Functional roles of proteins only detectable in male germ cells and the brain.**

In the main text, we wrote that: *“Although we found only 16 genes out of 5379 (0.3%) whose proteins were only detectable in both a male germ and brain cell, and no other cell type in any other (healthy) tissue (Supplementary Table 6), these represent potential entry points for further exploring the molecular parallels between the two organs (discussed further in the Supplementary Text).”* We elaborate on this point here.

We have discussed in the main text the functional associations with brain expansion for five of these 16 genes (*ERMN*, *HTR2A*, *HYLS1*, *OMG*, and *OPALIN*, each of which could be linked to either macrocephaly or relative increases in brain weight) and here outline the roles played by many of the remainder. We do not exclude the possibility that these proteins may also be expressed in unsampled tissues or at levels below the detection thresholds of the Human Protein Atlas dataset.

| Gene/protein | Evidence supporting a functional role in the brain |
| --- | --- |
| ASIC3 | Knockout in mice perturbs synaptic plasticity in the corticostriatal circuit, resulting in increased self-grooming behaviour (a repetitive behavioural phenotype common to mouse models of autism spectrum disorder) ^128^. (Aside from the brain, in the HPA v23, the protein encoded by this gene is only detected in spermatids). |
| CCDC136 | A vocal production-associated gene associated with multiple language-related phenotypes ^129^, and showing positive selection along the mammalian lineage ^130^. In vertebrates, including humans, the evolution of vocal learning has also been associated with specific neuroanatomical features, including cortical long-range projection neurons, in which this gene may play a role ^130^. (Aside from the brain, in the HPA v23, the protein encoded by this gene is only detected in spermatocytes and spermatids). |
| CDH7 | Cadherin-7, a cell adhesion molecule and member of the cadherin superfamily, all of which show spatially and temporally restricted expression profiles throughout the developing mouse ^131^ and marmoset ^132^ cortex. It is thought that because single neurons express multiple cadherins/protocadherins in a combinational manner throughout all layers of the cerebral cortex, and that because this results in a huge number of possible expression profiles, that combinatorial cadherin expression is a ‘neural code’, effectively providing a cell-surface molecular identity tag for individual neurons to enable self/non-self discrimination; this could facilitate the correct assembly of immensely complex neuronal circuits ^133^. (Aside from the brain, in the HPA v23, the protein encoded by this gene is only detected in spermatids). |
| GPR37L1 | G-protein coupled receptor that regulates the organisation of cortical astrocytes during development; in humans, mutations in GPR37L1 result in epilepsy/seizure phenotypes ^134^. (Aside from the brain, in the HPA v23, the protein encoded by this gene is only detected in spermatogonia and spermatids). |
| GRID2 | Highly conserved gene in mammals and, in mice, particularly expressed in the cerebellum throughout prenatal development; variants widely associated with cerebellar atrophy and, in humans, implicated in both bipedalism and speech development ^135^. (Aside from the brain, in the HPA v23, the protein encoded by this gene is only detected in spermatids). |
| GRM1 | Glutamate receptor primarily expressed in the cerebellum and functionally associated with synapse formation and synaptic plasticity; in humans, mutations in this gene have been associated with congential cerebellar ataxia ^136^. (Aside from the brain, in the HPA v23, the protein encoded by this gene is only detected in spermatids). |
| KCNH6 | Oncogene encoding a potassium ion channel and associated, bioinformatically, with the prognosis of glioblastoma ^137^. (Aside from the brain, in the HPA v23, the protein encoded by this gene is only detected in spermatogonia). |
| SH3GL3 | Tumour suppressor broadly associated with the invasive phenotypes of malignant gliomas ^138^ but more specifically appears to mediate the self-renewal activity of glioblastoma stem cells ^139^. (Aside from the brain, in the HPA v23, the protein encoded by this gene is only detected in spermatids). |

**(E) Evolutionary considerations on the brain-testis relationship.**

In the main text, we wrote that: *“Here we focus instead on evaluating the evidence that bears directly on the role of selfish spermatogonial selection. Given this emphasis, a full treatment of other evolutionary forces shaping the brain-testis relationship is beyond the scope of this work, although selected aspects are considered in the Supplementary Text.”* We elaborate on this point here.

Although this manuscript focuses on the potential influence of selfish spermatogonial selection on the evolutionary trajectory of the human condition, it is important to acknowledge that it represents only one of several evolutionary forces acting within the testis.

It is well established that sexual selection (arising when multiple males compete to fertilize female gametes) is a major force driving both behavioural and morphological diversification, including to many aspects of spermatogenesis (see reviews ^140,141^). Although sexual selection operates both before and after mating (in the former case by, for instance, males fighting for breeding territories) we concern ourselves here with post-copulatory sexual selection, or sperm competition, which arises when sperm from multiple males compete to fertilise female gametes. A well-established consequence is variation in relative testis size, which is routinely used as a proxy for sperm competition ^141^ as it strongly correlates with the total number of sperm in the ejaculate ^142^.

Comparative studies across diverse taxa consistently support this relationship. Species with multi-male mating systems – that is, a higher degree of sperm competition – typically have larger testes than those with single-male mating, including in anurans (frogs and toads) ^143,144^, birds ^145^, butterflies ^146^, fish ^147^, primates ^148,149^, and mammals more broadly ^150,151^. This association is sufficiently robust that relative testis size has been used to predict mating system in understudied species including nocturnal lemurs ^152^, New Zealand geckos ^153^, and West African riverine cichlids ^154^.

Testis size also varies with other elements of a species’ mating system. For instance, shorter mating seasons are associated with relatively larger testes in carnivorous mammals ^155^, an observation interpreted to reflect intensity of competition and a ‘lottery effect’ whereby producing more sperm increases the probability of reproductive success ^156^. Similarly, environmental and social cues can also modulate testis size within species. For example, in naturally promiscuous male deer mice (*Peromyscus maniculatus*), males reared in litters with more brothers develop larger testes as adults; this suggests that early development can signal future levels of competition in the wider population ^157^ but also, more broadly, that testis size shows adaptive plasticity in response to competitive context ^158^.

Nevertheless, while testis size is a comparatively accessible measurement, the trait ultimately selected under sperm competition is sperm production rate ^159^. In mammals, sperm competition selects for a faster rate of spermatogenesis, ostensibly enabling more rapid replenishment of reserves such that larger ejaculate volumes, or mating frequencies, can be maintained ^159^. Inter-specific variation in mammalian sperm competition (that is, relative testis size) thus conflates multiple sources of variation affecting production rate, both to testicular architecture (such as the proportion of seminiferous tubules in the testis) and the kinetics of spermatogenesis (such as seminiferous epithelium cycle length, or the relative efficiency of Sertoli cells) ^160^. In affecting spermatogonial stem cell divisions, selfish selection ultimately affects the kinetics of spermatogenesis, and in that respect it shares a point of connection with sperm competition.

Over the course of evolution, sexual selection has had widespread effects on body size and morphology. Polygynous species (single-male/multi-female mating) often exhibit pronounced male-biased size dimorphism ^161^ and both armaments and/or ornaments (antlers, plumes, and so on) ^162,163^. This implicitly positions testis size in a relationship with brain size – because the latter scales with body size too, in a log-curvilinear manner ^164^. Accordingly, variation in mammalian brain size correlates with many life-history traits related to reproduction: larger-brained species generally take longer to mature sexually, have longer gestation periods, and experience greater levels of parental investment ^165,166^. These trends are often explained in terms of developmental cost (larger brains take longer to grow) and a complex suite of evolutionary trade-offs reflecting myriad metabolic allocations and constraints. As such, brain size, testis size, body size, and mating system (among others) comprise an inter-dependent set of traits, each correlated to an extent with the other. For example, a comparative analysis of bats could relate mating system to both brain and testis size: promiscuous species had relatively smaller brains than monogamous species, potentially reflecting higher levels of sperm competition and comparatively greater investment in testis ^167^ (but see also ^168^, who argue this instead reflects differences in ecology). Conversely, across the avian phylogeny, there have been repeated rapid reductions in (expensive) testis mass for monogamous species, consistent with the idea that decreased sperm competition relaxes investment in an energetically costly organ, enabling investment elsewhere ^169^.

In each of the above cases, interpretation of the findings was framed with the long-standing ‘expensive tissue’ hypothesis in mind (see review ^170^), which proposed that because the brain is metabolically costly, increases in its size must be offset by energy reductions in other organs (chiefly the gut) ^171–173^. However, an extensive study of 100 mammals challenged this hypothesis by finding no negative correlation between brain size and the size of other organs, albeit not assessing testis ^174^. The study argued instead that energetic trade-offs to subsidise the brain were likely numerous and indirect. In humans specifically, the authors proposed that increased brain size was facilitated not by reducing a single costly organ but by a more diffuse reallocation of energy from the processes of growth, locomotion, and reproduction ^174^.

Given this, although we posit a link between brain expansion and oncogenic-like driver mutations in male germ cells, we would not expect to observe a direct cross-species relationship between relative brain and testis size. In other words, variation in brain size (beyond that predicted by allometry) would not necessarily track variation in testis size (beyond that predicted by allometry). Rather, we conjecture only that an evolutionary force acting specifically on the testis (but not sexual selection) could disproportionately enrich the genome for particular variants, some of which may subsequently prove advantageous in the brain. Any macroevolutionary signal linking testis biology to brain size would therefore be subtle and easily masked by multiple other variables. Indeed, a comparative study of 30 primates found no evidence of a significant correlation between relative brain and testis size ^175^, and nor did studies of 13 pinnipeds (seals, sea lions, and walruses) ^176^ and 68 shorebirds ^177^. An additional complication is that although extensive datasets of brain ^164,178^ and testis size ^54,150^ are available, when looking across a broad phylogenetic spectrum there is often little overlap between them, with measurements rarely obtained from the same individual or (due to the challenges of data collection) with replicates; this is especially the case for birds, where the testes are internal and vary in size seasonally ^179^. For example, in ^54^, there are 2036 species for which testis mass data is available, and in ^164^ (another publication from the same research group), 2807 species for which brain mass data is available, although only 996 species are common to both lists (that is, 26% of the combined set of 3847 species). This limited overlap reduces the power to detect subtle evolutionary associations and cautions against drawing strong conclusions from cross-species correlations alone.

As discussed above, mating system is particularly central to the convoluted relationship of brain, testis, and body sizes, with polygamous species (multi-male/multi-female breeding) expected to have stronger selection pressure for larger testes (higher sperm competition) and monogamous species (single-male/single-female breeding) stronger selection pressure for larger, more complex, brains (as sustaining pair bonds requires greater social acuity) ^175^. However, this relationship is further complicated by behavioural trade-offs: any link between brain size and mating system interacts with species-specific trends to either invest in mating opportunities (which promotes sperm competition) or parental care (which opposes it) ^180^. Further entwining brain development and mating system, a previous study comparing the neural transcriptomes of reproductive males in 10 monogamous and non-monogamous species pairs (from mice, voles, songbirds, frogs, and fishes) identified a conserved ‘vertebrate monogamy signature’ of 24 genes, all up-regulated in monogamous males and enriched in neurodevelopmental functions ^181^. Notably, many of these genes also have roles in male fertility. Examples include *ANK2* (involved in sperm motility and aberrantly hypermethylated in the sperm of infertile men ^182^), *HIP1* (required for spermatogonial differentiation ^183^), *NOTCH1* (conditional activation of which ablates the male germline in mice ^184^), *RBL1* (which regulates germ cell proliferation ^185^) and *SMAD9* (member of a protein family central to primordial germ cell specification ^186^, one of which, *SMAD4*, is a *bona fide* selfish gene ^2^). In this respect, these observations further support the idea of a bidirectional link between genes functional in both the brain and male germline.

**(F) From correlation to causation: do mutations in genes involved in spermatogonial activity have a demonstrable effect upon brain size?**

In the main text we wrote that: *“Enrichment of variants selfishly selected in the testis could also contribute by subtly modulating stem cell turnover rates, thereby influencing the duration and extent of neurogenic output. Although in support of this hypothesis we have summarised a large body of work on male germline mutations as they relate to brain expansion, experimental work to establish a causal relationship will be necessary. Future approaches could consider to what extent genes and mutations associated with sperm production rate (such as number or turnover rate of spermatogonial stem cells) are associated with brain phenotypes. This is an underexplored research area, with few obvious models in which to address it, although some tentative directions are discussed in the Supplementary Text.”* We elaborate on this point here.

To experimentally demonstrate the effects of an evolutionary change in spermatogonial activity (or other male fertility or sexual traits) upon brain size, one of the most powerful approaches is to use artificial selection on one trait and then to measure the response in other traits ^187^. To the best of our knowledge, this approach has yet to be applied specifically to testis size (or sperm count) and brain size, although we think it is not without promise (assuming it is experimentally possible). This is because a number of studies already support a positive genetic correlation between male sexual traits and brain size, albeit on a species-by-species basis (such that the results may not necessarily generalise) and only for pre- rather than post-copulatory male traits, mediated by mating behaviour (that is, to our knowledge, no significant association with brain size has been found for relative testis size or sperm count, although as discussed in section E such a signal, if it existed, would likely be subtle and easily masked). In this section, we survey a number of experimental models which establish associations between the brain and male fertility (or other sexual) traits, and discuss their suitability for assessing our own hypothesis – namely, that ‘selfish’ mutations arising in spermatogonial stem cells not only promote their own propagation in the testis but influence neural progenitor biology (and thereby neuron production rate and, by extension, brain size) once inherited.

One model to consider is mosquitofish (*Gambusia holbrooki*). Interestingly, mosquitofish artificially selected for increased gonopodium (intromittent male genital) length were found to produce offspring with larger brains – but only for daughters ^188^. To interpret this observation the authors note that male mosquitofish do not court but instead attempt up to a thousand coercive copulations a day ^189^ (sneaking up on females from behind) such that a longer gonopodium, in facilitating sperm transfer, is associated with greater male reproductive success ^190^. Increased brain size in females conceivably evolved in response to this heightened sexual conflict – as females suffer more from being forced to mate than males do by losing an opportunity, they should experience stronger selection for cognitive abilities to evade harassment, detect concealed males quicker, or otherwise better predict their environment. The broader point here, however, is that in mosquitofish the genetic correlation between male sexual traits and brain size could be explained by the (pre-copulatory) behavioural interactions of the sexes, thereby rendering this species an unsuitable model for addressing our specific hypothesis (which, by centring on spermatogonia, implicates a stronger role for post- rather than pre-copulatory selection).

By contrast to a coercive mating strategy, courting behaviours are thought more cognitively demanding (because to make an adaptive mate choice, one must remember and compare past encounters) and so species which engage in them should demonstrate stronger associations between brain size and sexual traits. This is supported by evidence from guppies which, unlike mosquitofish, engage in both mating tactics, coercing and courting ^187^. Guppies artificially selected for relatively larger brains – and which have greater cognitive ability as a consequence ^191–193^ – were found to have significantly higher expression of several male sexual traits, including gonopodium length and both tail fin length and bodily iridescence (which are not only attractive to females but more so for larger-brained females ^194^), although it should be noted that – as with the positive correlation of male sexual traits and brain size in mosquitofish – relative testis size and total sperm number were unaffected ^187^, diminishing the promise of guppies as a model. Possible explanations for this pattern include a common genetic architecture (through pleiotropy or linkage) underlying phenotypic variation in both traits, or that directional selection for brain size affects another variable correlated both with brain size and sexual traits (the authors suggest body condition as a candidate ‘third variable’ ^187^). Despite the unsuitability of these species for addressing our hypothesis experimentally, it remains encouraging to note these observations support the more general point that reproductive and cognitive phenotypes can co-evolve.

Another experimental system to consider is the mouse. However, although globally over a thousand mouse models have fertility phenotypes, around 99% of these show decreased fertility ^195^. An exception presents itself in the form of outbred Dummerstorf FL1 (‘fertility line 1’) mice, developed at the Leibniz Institute for Farm Animal Biology, which have been continuously selected for female index traits (large litter size with high average birth weight) since the 1970s ^196^. Although fertility selection was female-focused, relative to a random-mated control line male characteristics were also apparent and which contribute to the ‘high fertility’ phenotype ^196^. While this may be expected given that many genes necessary for reproductive success are shared between females and males ^197^, dedicated breeding experiments involving these mice concluded that the fertility phenotype depends largely on the female genetic background, with only a marginal (but nevertheless, still significant) contribution by the male ^198^. Male traits of the F1 line include a significantly increased proportion of diploid cells in the testicular parenchyma that were actively synthesising DNA ^199^, with subsequent flow cytometric analysis characterising this population primarily as Leydig cells – although, of particular interest to our hypothesis, it also contained a significantly increased number of mitotically-dividing spermatogonia ^200^. To the best of our knowledge, it is unknown what genes or mutations underlie this increase, or whether they have any pleiotropic association with brain expansion. Yet, by virtue of selection not only for fertility (larger litters) but for heavier litters, the average birth weight of FL1 mice is double that of the controls ^201^ and so by extension, they must also grow larger brains (produce more neurons) in the same gestational time.

We have previously discussed (in section C) a bidirectional link between genes functional in both the human brain and male germline, and can also find support for this in FL1 mice. Microarray profiling of the FL1 testis identified 358 genes differentially expressed relative to a random-mated control line – although, surprisingly, only 12 could be associated with a known mouse reproductive phenotype ^201^. However, with one exception, all 12 could also be functionally associated with some aspect of neurodevelopment (with the human orthologues of five of these genes – *CNBD2*, *CCND2*, *DNAH11*, *NR2C2*, and *NSUN2* – listed in **Supplementary Table 1**). Genes upregulated in the FL1 testis include *Adamts5* (associated with both neuroplasticity ^202^ and glioma invasiveness ^203^), *Cnbd2* (the exception, having no explicit functional link with neurodevelopment, although nevertheless associated with IQ in humans ^204^), *Ccnd2* (associated with both macrocephaly ^205,206^ and megalencephaly in humans ^207^), *Mfge8* (expressed by both astrocytes and microglia in cortical grey matter and with a role in synaptic pruning ^208^), and *Ttll1* (a polyglutamylase which regulates neuronal polarity ^209^, disruption of which produces neurodegenerative phenotypes ^210^). Genes downregulated in the FL1 testis include *Adgrd1* (necessary for glioblastoma growth ^211^), *Agt* (involved in the regulation of cerebral blood flow and synaptic transmission ^212^), *Dnah11* (involved in synaptic development and hearing loss ^213^), *Insl5* (neuroendocrine calcium regulator expressed in the mouse hypothalamus ^214^), *Nr2c2*/*Tr4* (knockout of which results in abnormal cerebellar cytoarchitecture in mice ^215^), *Nsun2* (neuronal deficiency of which is associated with impaired synaptic signalling ^216^), and *Sult1e1* (associated with both ischemic stroke ^217^ and meningioma ^218^).

Unfortunately, there is limited antibody staining data in the Human Protein Atlas ^44^ that allows conclusions to be drawn – assuming results in humans are analogous to mice – about whether the proteins encoded by these genes are specifically expressed in spermatogonia. This is because for 10 of the 12 genes, experiments were only performed for the non-specific category of ‘cells in seminiferous ducts’; the exceptions were NSUN2 and TTLL1, both of which were indeed detected in spermatogonia. Nevertheless, protein-level expression in Leydig cells – which, noted above, is the cell population most substantially increased in the FL1 testis ^201^ – was also found for 6 of these genes (ADGRD1, CNBD2, MFGE8, NR2C2, NSUN2, and TTLL1; no Leydig cell expression was found for ADAMTS5, AGT, CCND2, DNAH11, INSL5 or SULT1E1).

As such, an equivocal case could be made for using FL1 mice to explore how factors resulting in higher numbers of spermatogonia may also play causal roles in brain growth – because they show increased expression in the testis (if not unambiguously in spermatogonia themselves) for genes associated with brain development. Nevertheless, it remains the case that selection of the FL1 line was not male-focused, and that it did not select exclusively for increased fertility. An experimental model that specifically isolates genes involved in increased spermatogonial activity thus remains both elusive and desirable.

**SUPPLEMENTARY FIGURES**

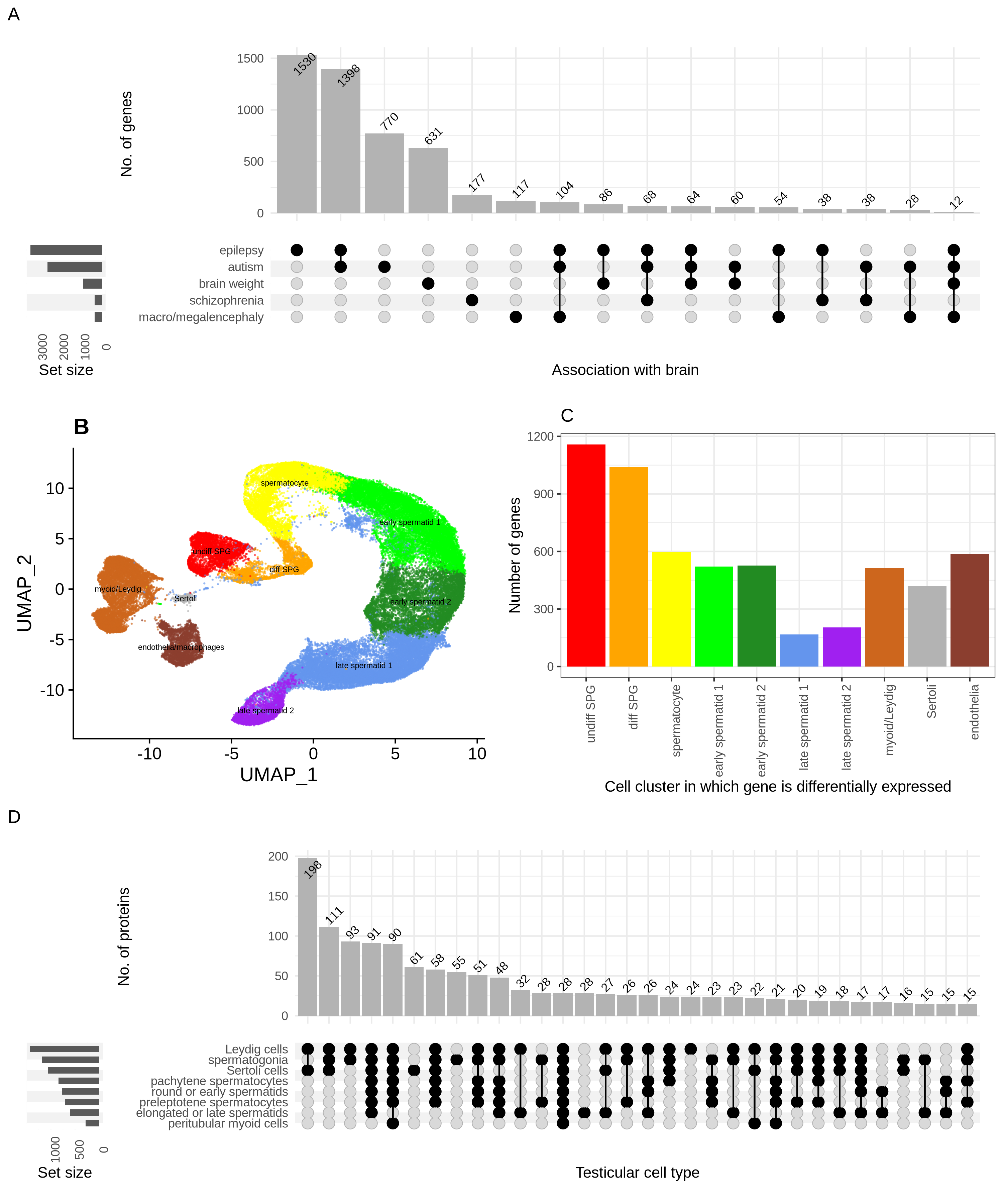

**Supplementary Figure 1. A conservative set of brain-associated genes and their expression in the human testis.**

**(A)** Distribution of 5212 genes among an explicit set of brain-associated phenotypes, detailed further in notes to **Supplementary Table 1**. Sets containing < 10 genes are not shown. **(B)** Single-cell expression atlas of the adult human testis (n = 60,427 cells), annotated into ten cell clusters (3 somatic, 7 germline). **(C)** Number of genes differentially expressed, at the transcript level, in at least one of these ten clusters (raw data in **Supplementary Tables 2 and 3**). **(D)** Protein-level expression of 1573 brain-associated genes for which antibody staining data was available for each of 8 testicular cell types (3 somatic, 5 germline) and with a protein detected in at least one (raw data from https://v23.proteinatlas.org/about/download/normal_tissue.tsv, accessed 17^th^ June 2023). Sets represent detectable protein-level expression (i.e. classification into either of the HPA categories of low, medium or high expression) in each cell type (raw data in **Supplementary Table 1**). Sets containing < 15 proteins are not shown.

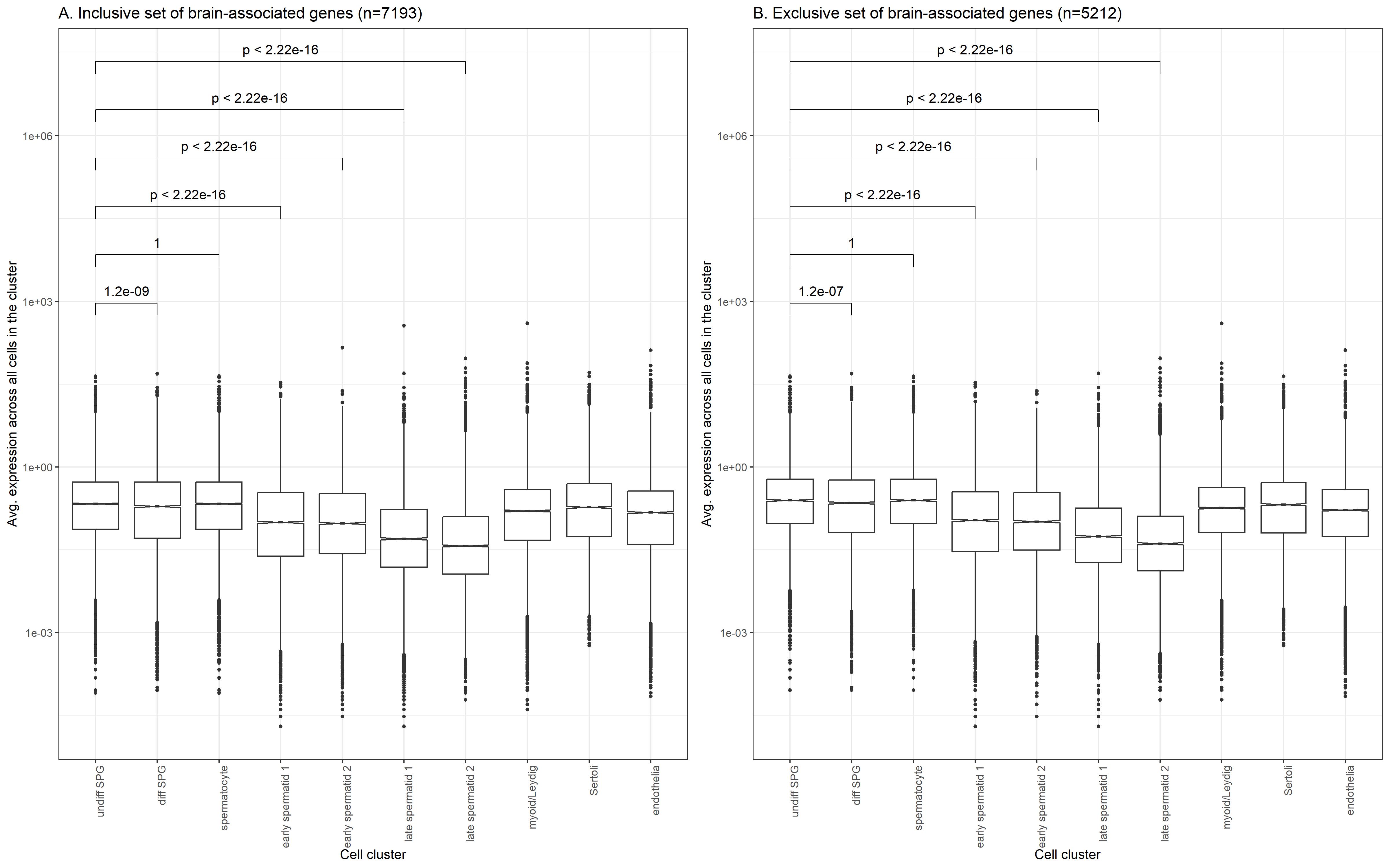

**Supplementary Figure 2. Average expression across, respectively, an (A) inclusive and (B) conservative set of 7193 and 5212 brain-associated genes for each of ten cell clusters in an adult human testis expression atlas.**

The brain-associated gene sets are detailed in **Supplementary Table 1** with expression level per testicular cell type in **Supplementary Table 2**. Statistical comparisons are Mann-Whitney U tests, implemented using the R package ‘ggpubr.’

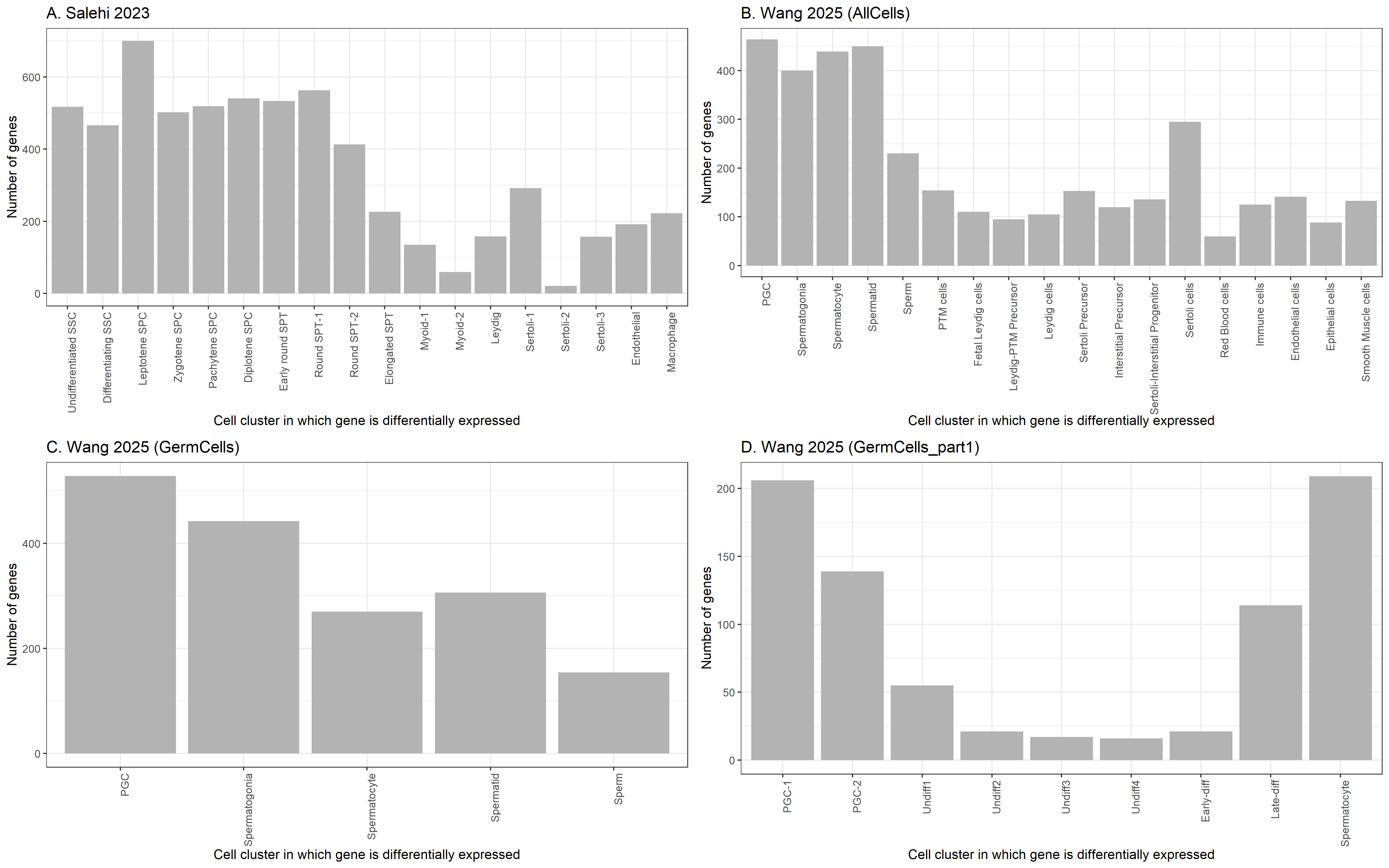

**Supplementary Figure 3. Number of differentially expressed brain-associated genes per cell cluster, for each of four testis and male germline single-cell atlases, and using an inclusive set of brain-associated genes.**

For each of the four single-cell atlases, barplots show the number of genes significantly differentially expressed within an inclusive set of brain-associated genes (n=7193). The cell atlases and their respective annotations are described in **Supplementary Table 4**, alongside the differential expression analysis results, which were filtered according to a common set of criteria. The brain-associated gene sets are detailed in **Supplementary Table 1**.

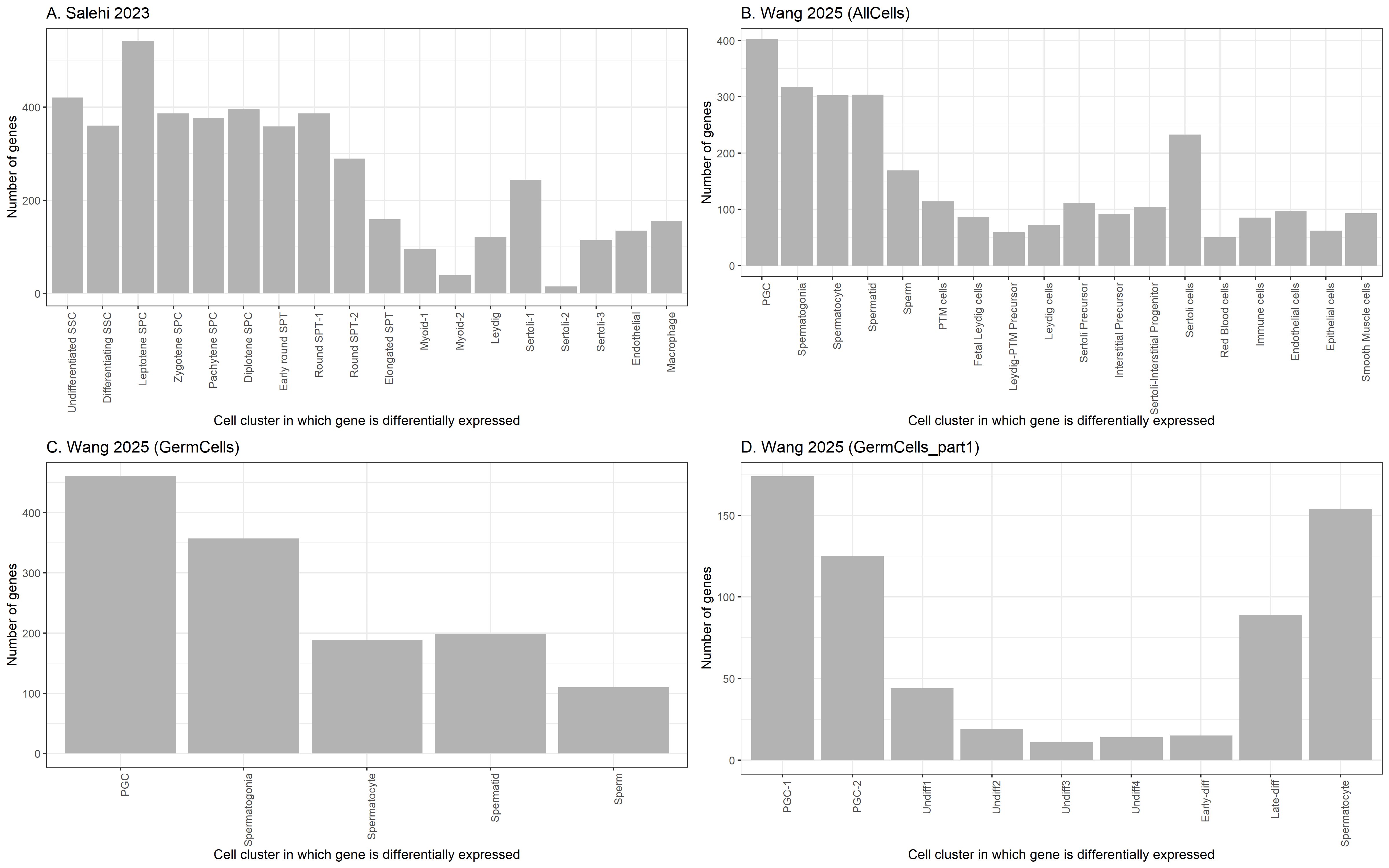

**Supplementary Figure 4. Number of differentially expressed brain-associated genes per cell cluster, for each of four testis and male germline single-cell atlases, and using a conservative set of brain-associated genes.**

For each of the four single-cell atlases, barplots show the number of genes significantly differentially expressed within an inclusive set of brain-associated genes (n=5212). The cell atlases and their respective annotations are described in **Supplementary Table 4**, alongside the differential expression analysis results, which were filtered according to a common set of criteria. The brain-associated gene sets are detailed in **Supplementary Table 1**.

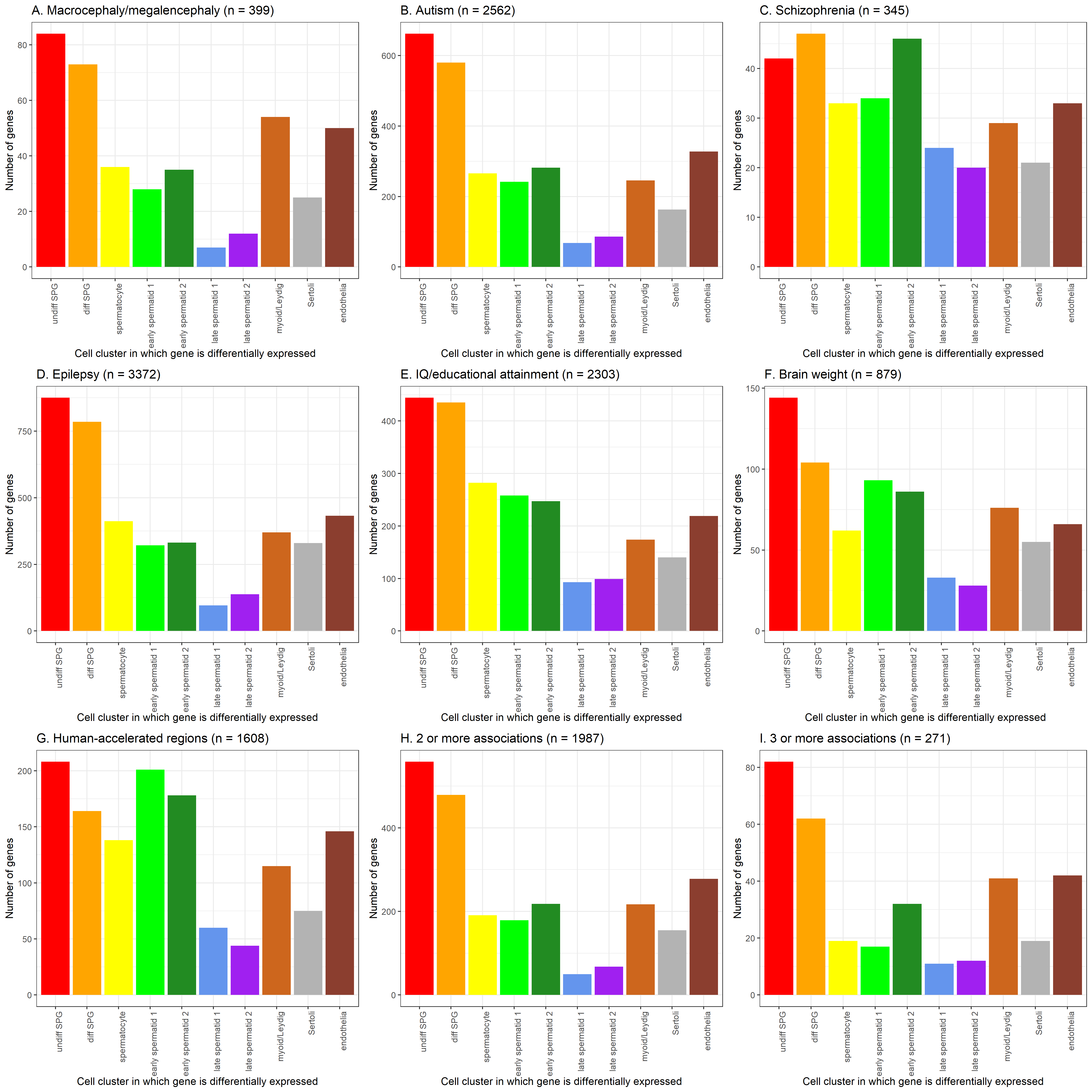

**Supplementary Figure 5. Number of brain-associated genes differentially expressed, at the transcript level, in one or more of ten testicular cell clusters.**

The list of brain-associated genes is given in **Supplementary Table 1**, with raw data for this figure in **Supplementary Tables 2 and 3**. Panels A to G show the distribution of differentially expressed genes for seven different categories of phenotype, whereas panels H and I require that each gene is associated with at least 2 or 3 different categories, respectively, not including the two ‘indirect’ phenotypes of IQ/attainment and human-acceleration.

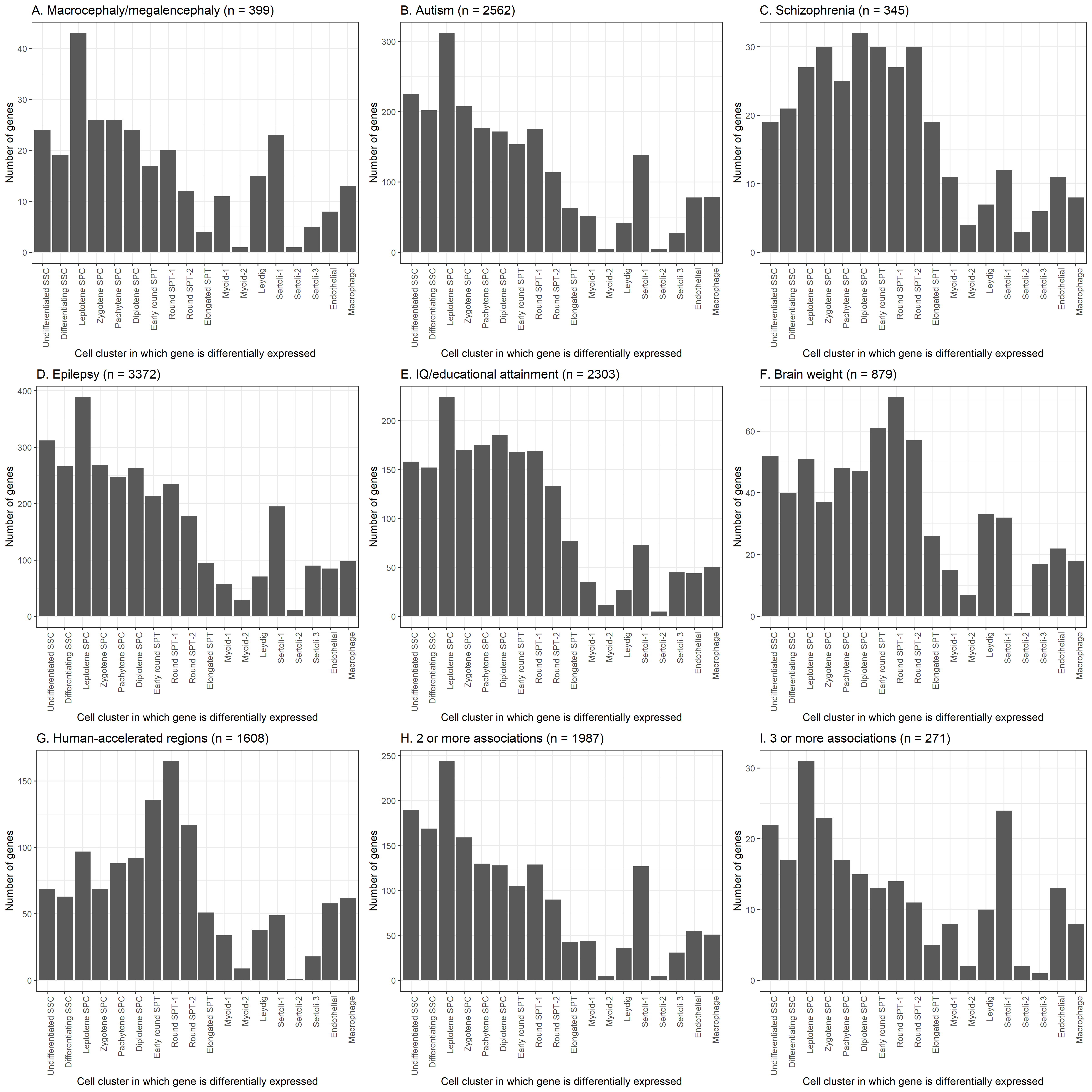

**Supplementary Figure 6. Number of brain-associated genes differentially expressed, at the transcript level, in one or more of cell clusters from the ‘Salehi 2023’ single-cell expression atlas.**

The ‘Salehi 2023’ expression atlas, its respective annotations, and the differential expression analysis results used for this figure, are detailed in **Supplementary Table 4**. The brain-associated gene sets are detailed in **Supplementary Table 1**. Panels A to G show the distribution of differentially expressed genes for seven different categories of phenotype, whereas panels H and I require that each gene is associated with at least 2 or 3 different categories, respectively, not including the two ‘indirect’ phenotypes of IQ/attainment and human-acceleration.

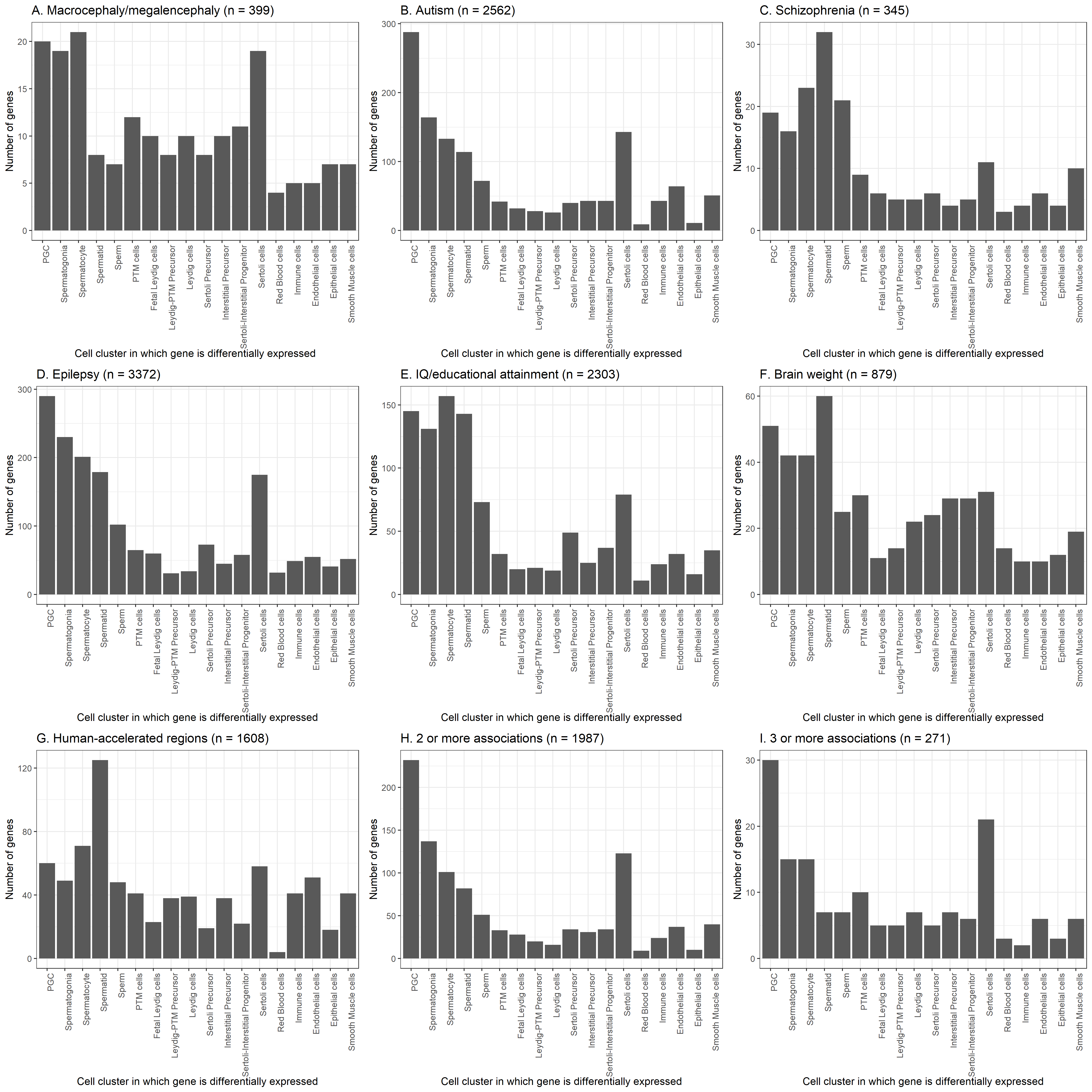

**Supplementary Figure 7. Number of brain-associated genes differentially expressed, at the transcript level, in one or more of cell clusters from the ‘Wang 2025 (all cells)’ single-cell expression atlas.**

The ‘Wang 2025 (all cells)’ expression atlas, its respective annotations, and the differential expression analysis results used for this figure, are detailed in **Supplementary Table 4**. The brain-associated gene sets are detailed in **Supplementary Table 1**. Panels A to G show the distribution of differentially expressed genes for seven different categories of phenotype, whereas panels H and I require that each gene is associated with at least 2 or 3 different categories, respectively, not including the two ‘indirect’ phenotypes of IQ/attainment and human-acceleration.

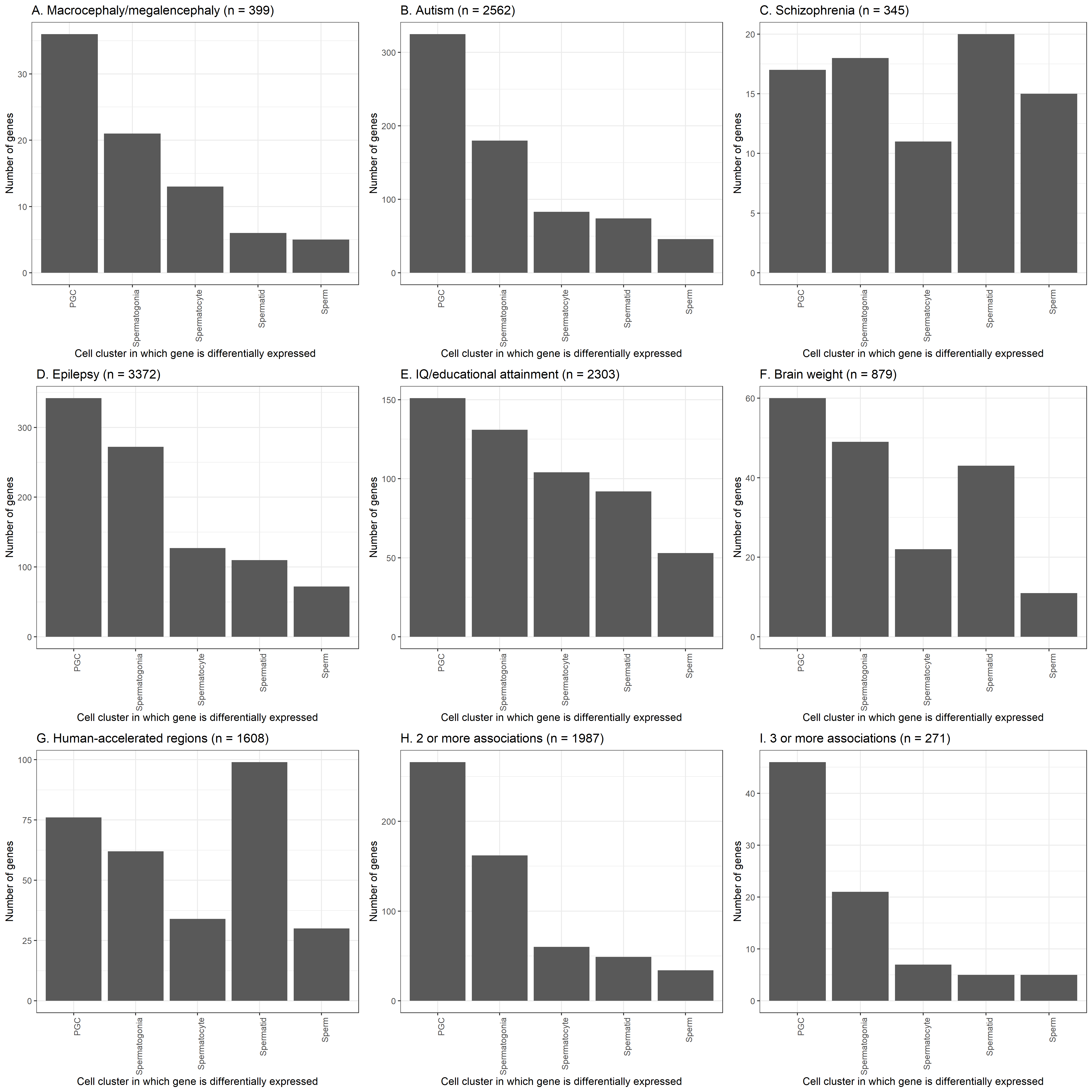

**Supplementary Figure 8. Number of brain-associated genes differentially expressed, at the transcript level, in one or more of cell clusters from the ‘Wang 2025 (germ cells)’ single-cell expression atlas.**

The ‘Wang 2025 (germ cells)’ expression atlas, its respective annotations, and the differential expression analysis results used for this figure, are detailed in **Supplementary Table 4**. The brain-associated gene sets are detailed in **Supplementary Table 1**. Panels A to G show the distribution of differentially expressed genes for seven different categories of phenotype, whereas panels H and I require that each gene is associated with at least 2 or 3 different categories, respectively, not including the two ‘indirect’ phenotypes of IQ/attainment and human-acceleration.

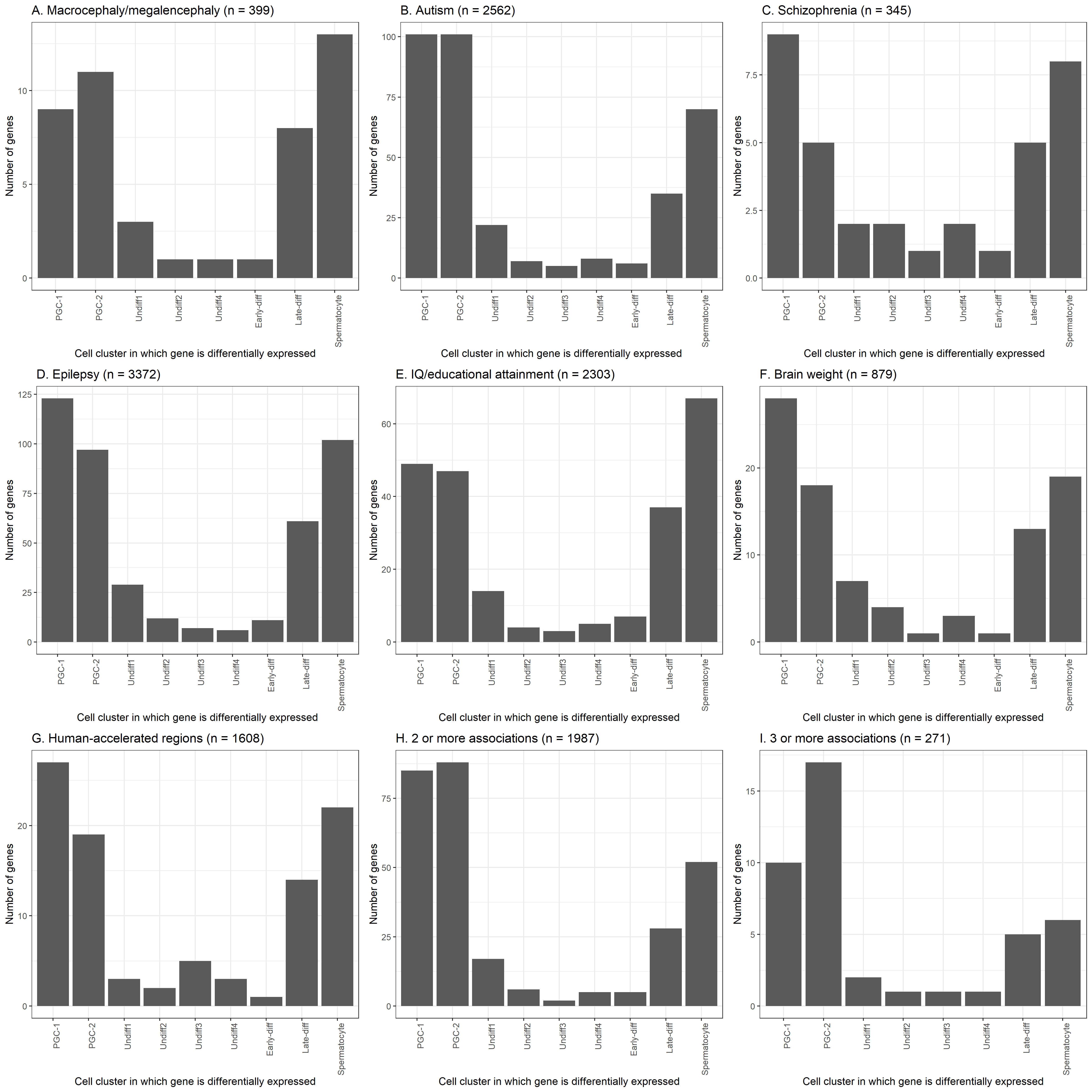

**Supplementary Figure 9. Number of brain-associated genes differentially expressed, at the transcript level, in one or more of cell clusters from the ‘Wang 2025 (germ cells – part 1)’ single-cell expression atlas.**

The ‘Wang 2025 (germ cells – part 1)’ expression atlas, its respective annotations, and the differential expression analysis results used for this figure, are detailed in **Supplementary Table 4**. The brain-associated gene sets are detailed in **Supplementary Table 1**. Panels A to G show the distribution of differentially expressed genes for seven different categories of phenotype, whereas panels H and I require that each gene is associated with at least 2 or 3 different categories, respectively, not including the two ‘indirect’ phenotypes of IQ/attainment and human-acceleration.

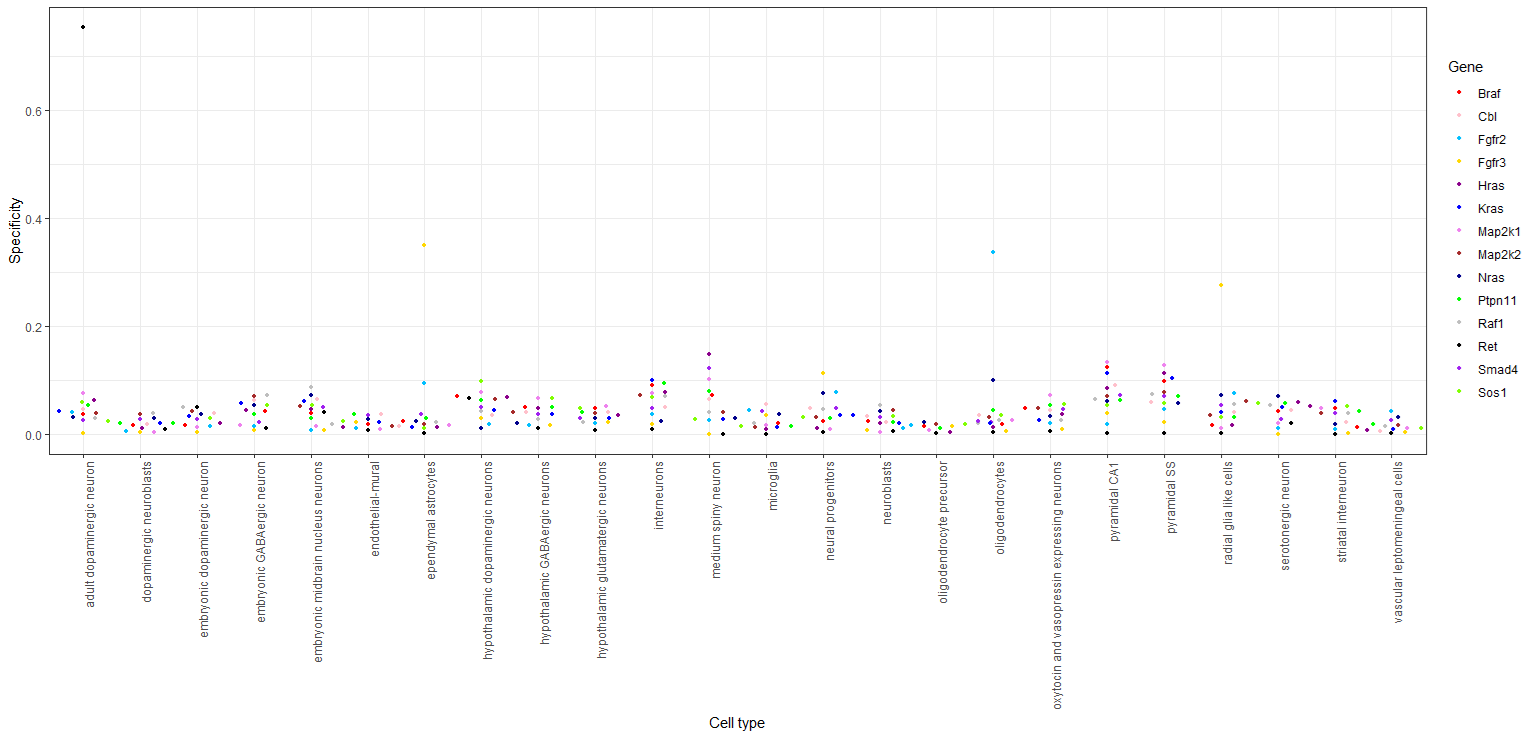

**Supplementary Figure 10. Beeswarm (violin scatter) plot of the cellular specificity, in the mouse brain, of 14 genes known to harbour selfish spermatogonial mutations.**

Cellular specificity values, from the Karolinska superset ^122^, represent the proportion of the total expression of a gene attributable to one of 24 brain cell types (a specificity of 0 means that the gene is not expressed in that cell type and a value of 1 that it is only expressed in that cell type). The set of 14 genes known to harbour one or more selfish spermatogonial mutations have been identified in humans ^1,2^ although for the purpose of visualisation here, specificity data from the one-to-one mouse orthologue is used.

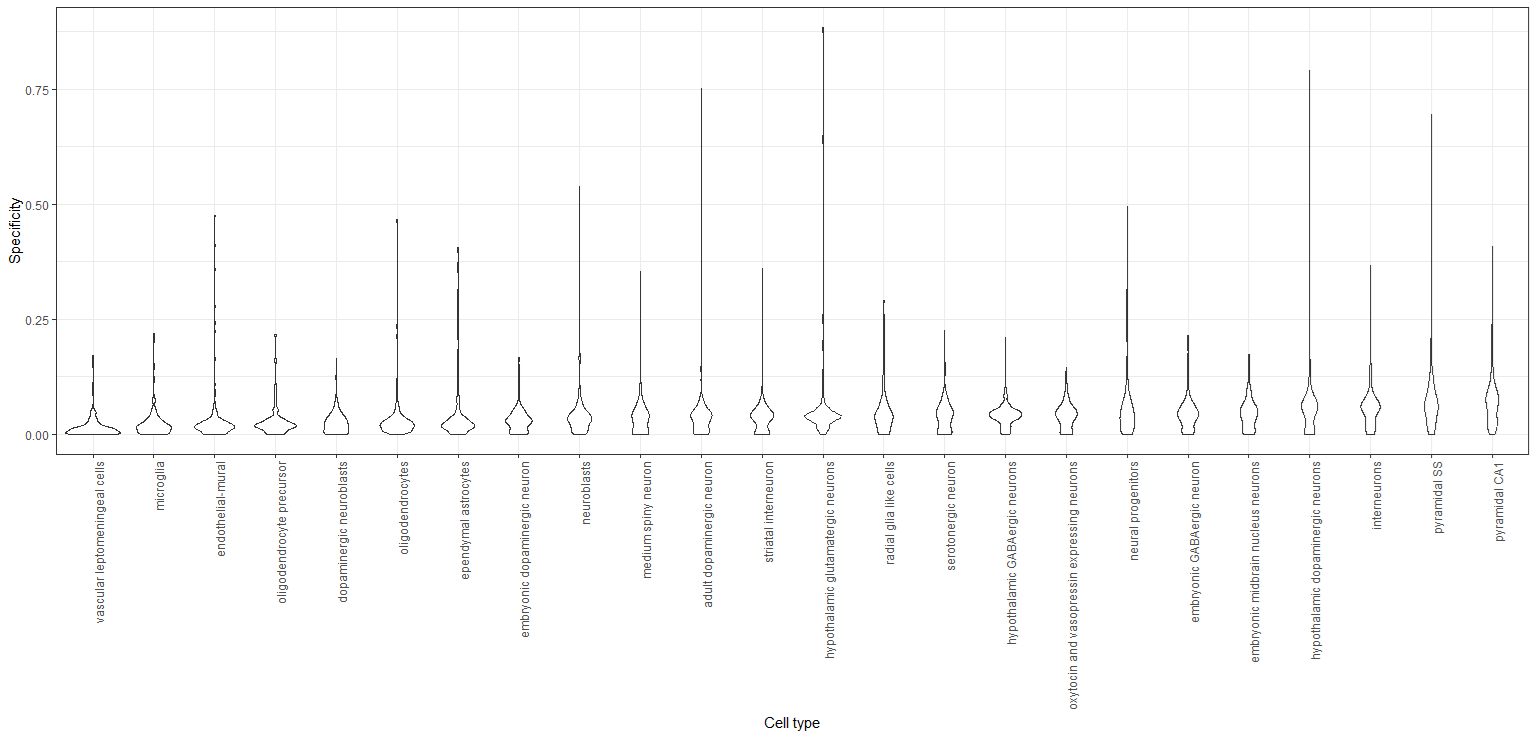

**Supplementary Figure 11. Violin plot of the cellular specificity of 218 spermatogonia and spermatogenesis-associated genes in the mouse brain.**

Cellular specificity values, from the Karolinska superset ^122^, represent the proportion of the total expression of a gene attributable to one of 24 brain cell types (a specificity of 0 means that the gene is not expressed in that cell type and a value of 1 that it is only expressed in that cell type). Violins are sorted from left to right in ascending order of median specificity. The genes used for this plot are listed in **Supplementary Table 5** and drawn from previous studies ^106,107,109,118,119^. The genes in this table are human and for the purpose of visualisation here, the one-to-one mouse orthologue (if extant) is used.

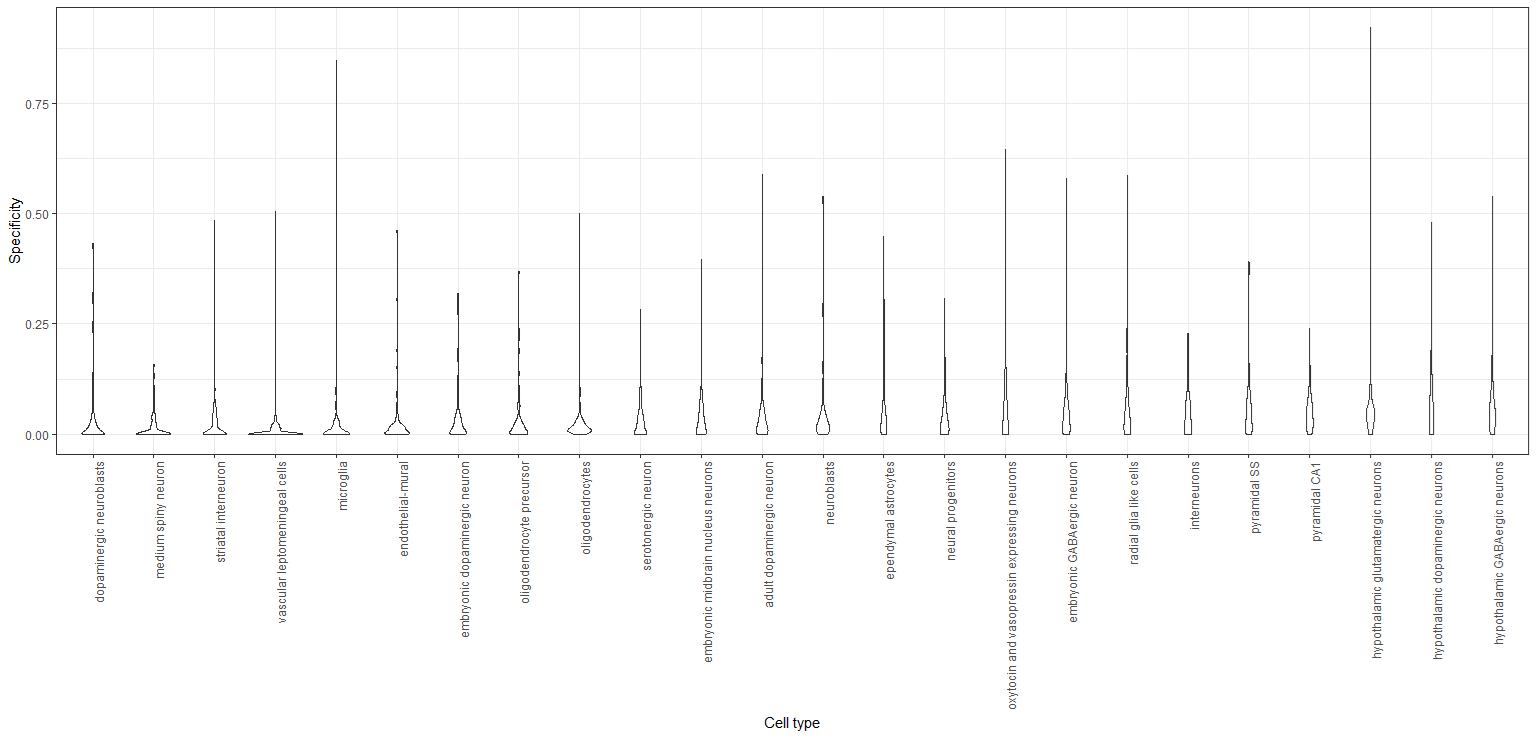

**Supplementary Figure 12. Violin plot of the cellular specificity of 154 male infertility-associated genes in the mouse brain.**

Cellular specificity values, from the Karolinska superset ^122^, represent the proportion of the total expression of a gene attributable to one of 24 brain cell types (a specificity of 0 means that the gene is not expressed in that cell type and a value of 1 that it is only expressed in that cell type). Violins are sorted from left to right in ascending order of median specificity. The genes used for this plot are listed in **Supplementary Table 5** and drawn from two previous systematic reviews ^120,121^. The genes in this table are human and for the purpose of visualisation here, the one-to-one mouse orthologue (if extant) is used.

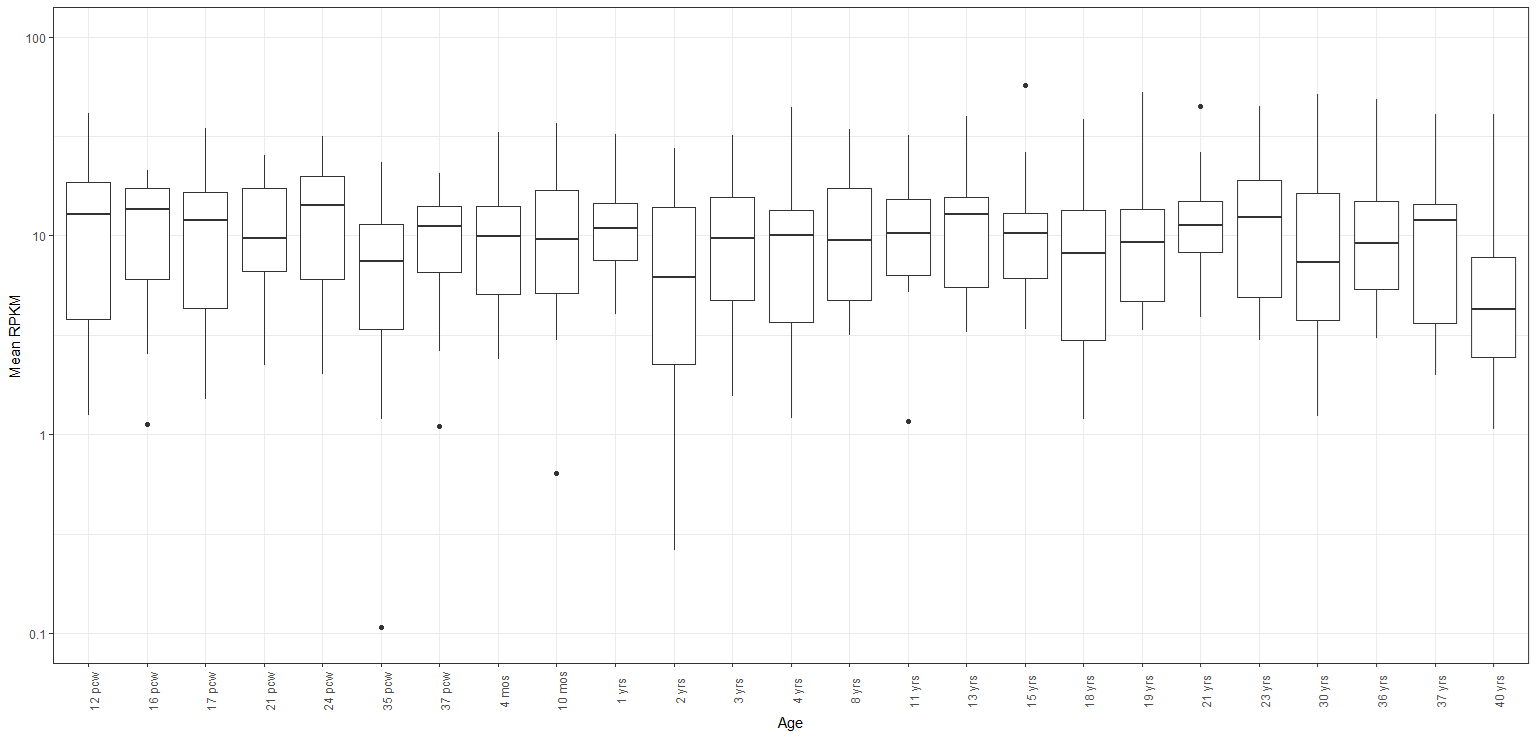

**Supplementary Figure 13. Boxplot of the expression in the cerebellar cortex over time of 14 genes known to harbour selfish spermatogonial mutations, using the Allen BrainSpan Atlas.**

The set of 14 genes known to harbour one or more selfish spermatogonial mutations have been described in previous studies ^1,2^.

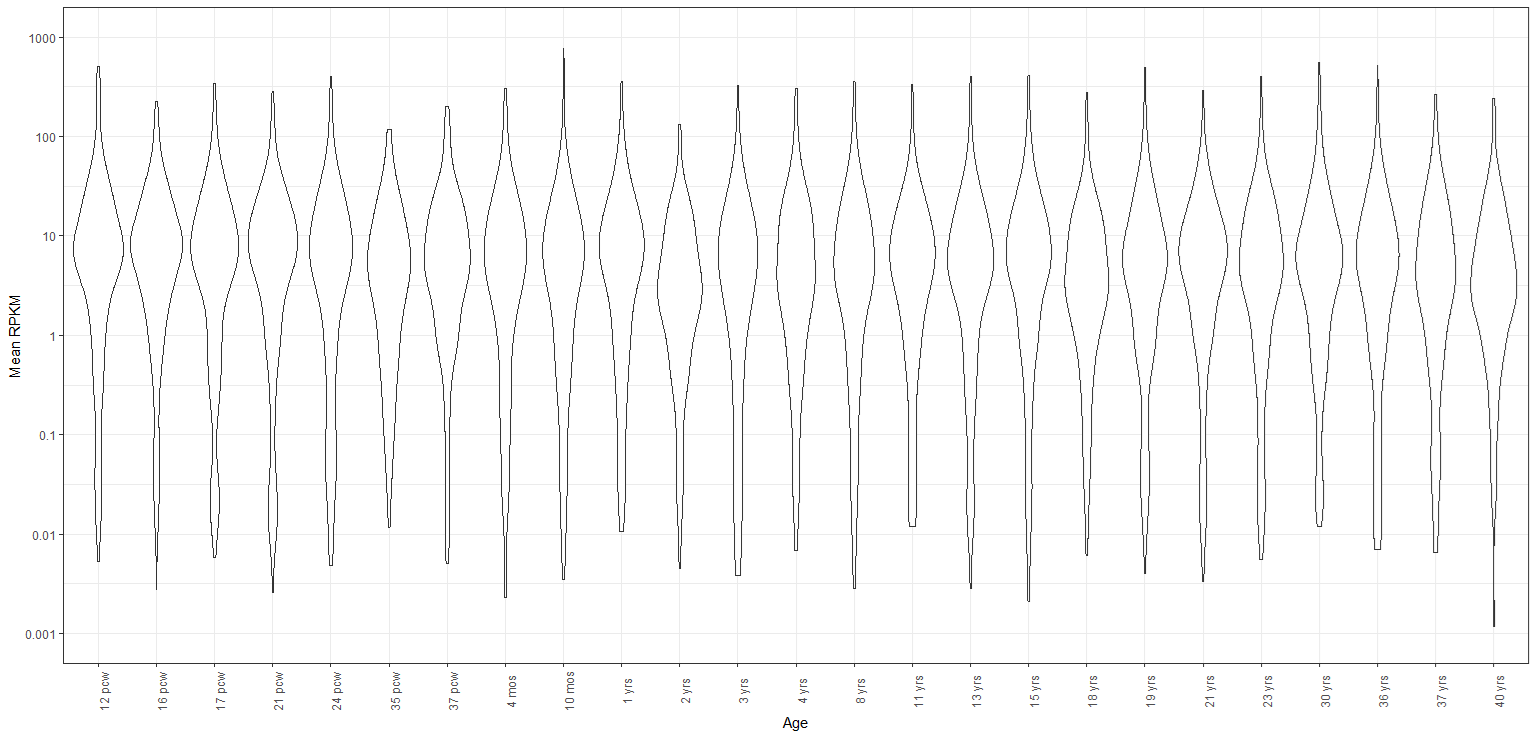

**Supplementary Figure 14. Violin plot of the expression in the cerebellar cortex over time of 218 spermatogonia and spermatogenesis-associated genes, using the Allen BrainSpan Atlas.**

The genes used for this plot are listed in **Supplementary Table 5** and drawn from previous studies ^106,107,109,118,119^.

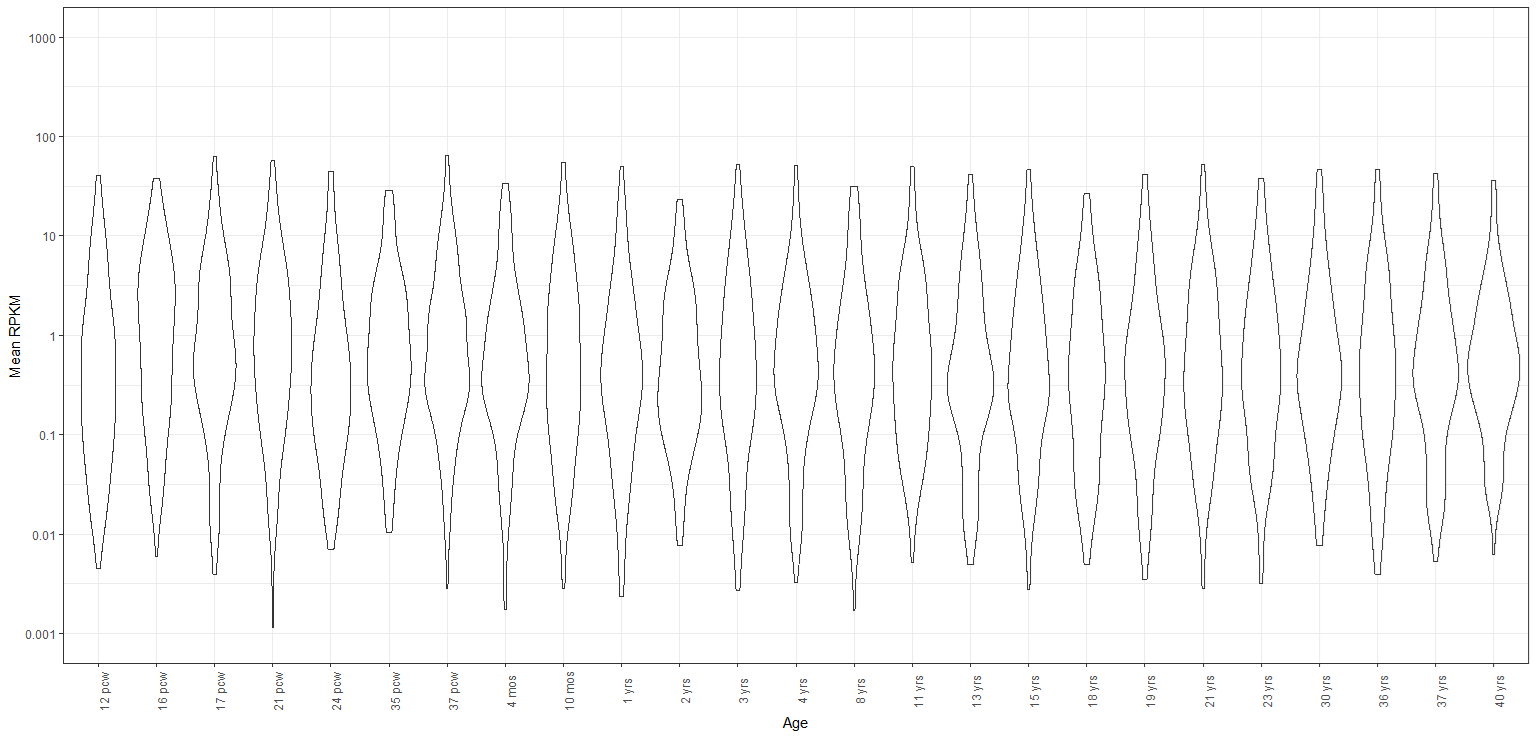

**Supplementary Figure 15. Violin plot of the expression in the cerebellar cortex over time of 154 male infertility-associated genes, using the Allen BrainSpan Atlas.**

The genes used for this plot are listed in **Supplementary Table 5** and drawn from two previous systematic reviews ^120,121^.

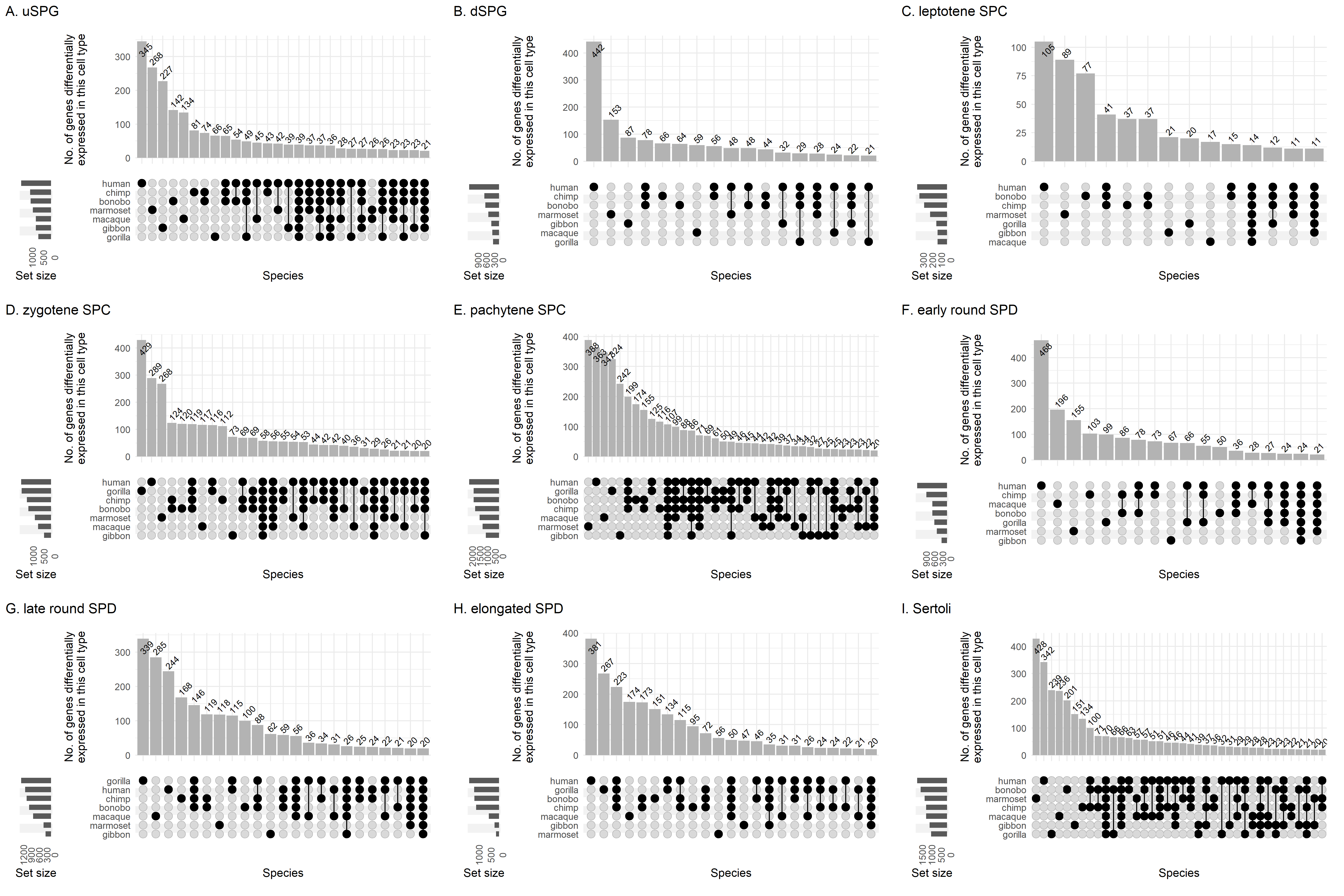
**Supplementary Figure 16. Degree of overlap between marker genes for each of 9 testicular cell types in each of 7 primate species.**

This figure represents a re-analysis of data from ^219^. To compare gene names across species, we only plotted those that could be assigned to an orthogroup, defined as a set of genes whereby every member had a high-confidence one-to-one orthologue with at least one other member. Orthogroups are defined in **Supplementary Table 8** with their differential expression in each testicular cell type per species detailed in **Supplementary Table 9**. uSPG, undifferentiated spermatogonia; dSPG, differentiated spermatogonia; SPC, spermatocyte; SPD, spermatid.

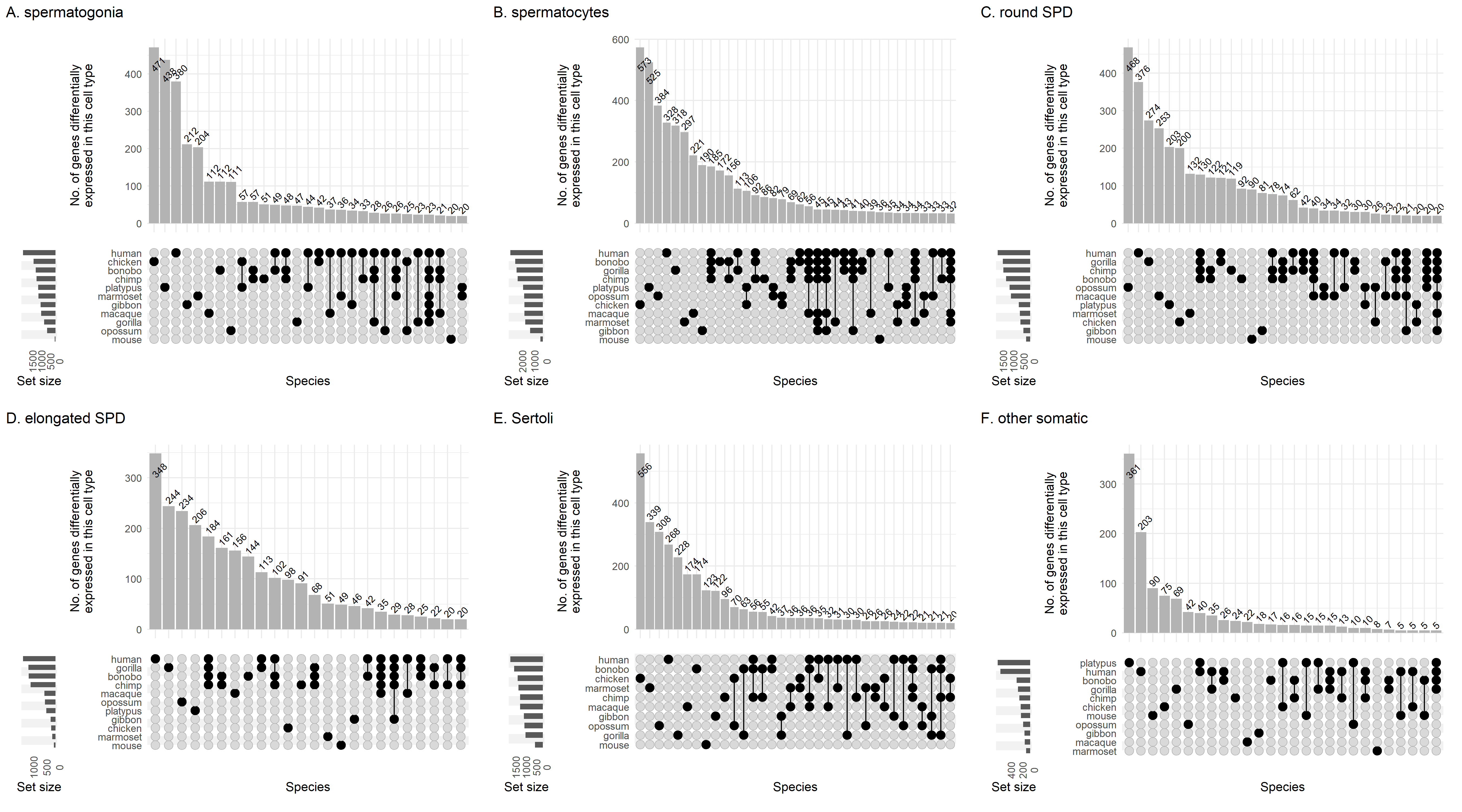

**Supplementary Figure 17. Degree of overlap between marker genes for each of 6 testicular cell types in each of 11 animal species.**

This figure represents a re-analysis of data from ^219^. To compare gene names across species, we only plotted those that could be assigned to an orthogroup, defined as a set of genes whereby every member had a high-confidence one-to-one orthologue with at least one other member. Orthogroups are defined in **Supplementary Table 8** with their differential expression in each testicular cell type per species detailed in **Supplementary Table 9**. SPD, spermatid.

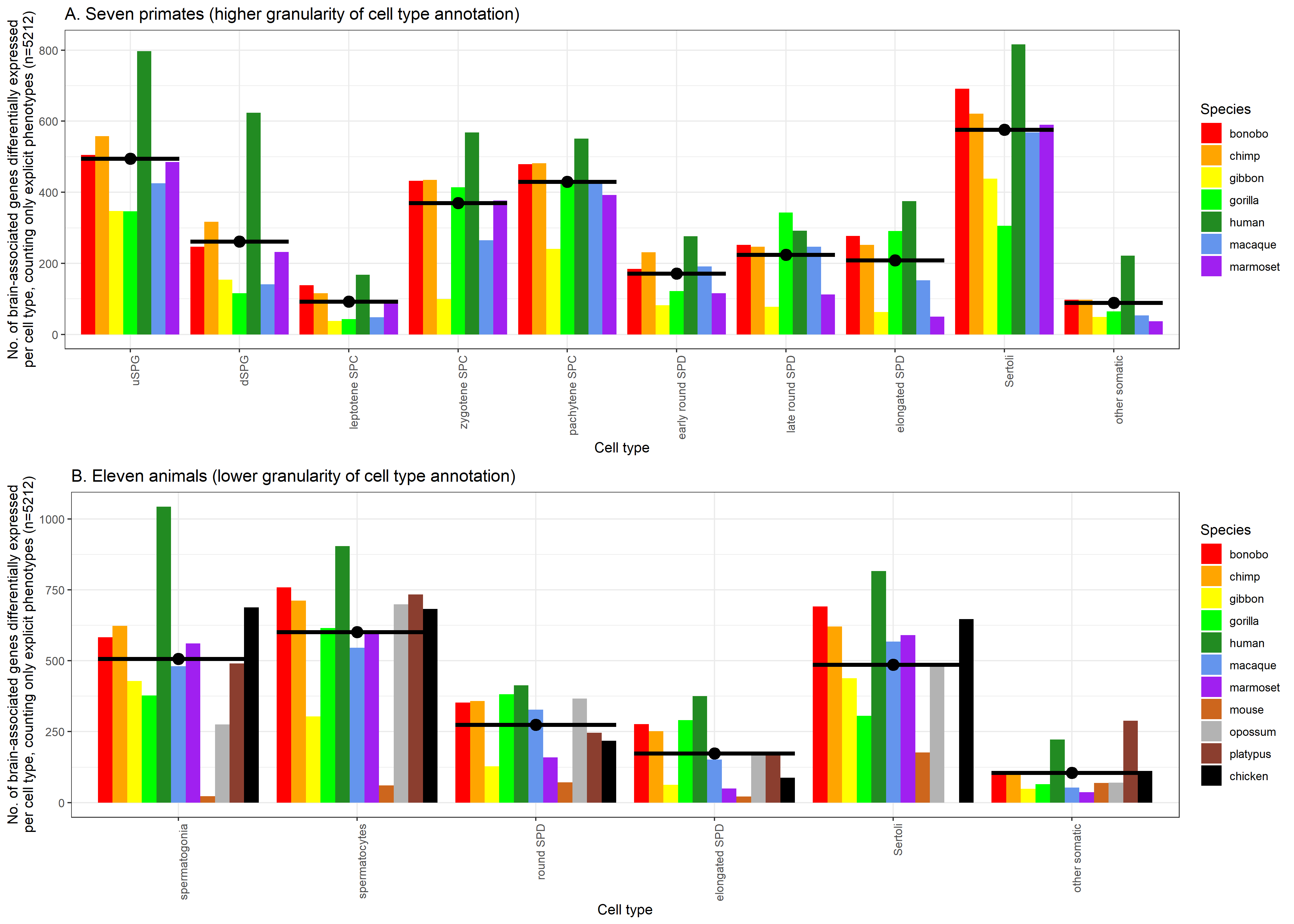
**Supplementary Figure 18. Number of brain-associated genes that are also testicular cell type markers in (A) 7 primate, and (B) 11 animal species, using an inclusive set of 5212 brain-associated genes.**

This figure represents a re-analysis of data from ^219^. To compare gene names across species, we only plotted those that could be assigned to an orthogroup, defined as a set of genes whereby every member had a high-confidence one-to-one orthologue with at least one other member. Orthogroups are defined in **Supplementary Table 8** with their differential expression in each testicular cell type per species detailed in **Supplementary Table 9**. The contents of the brain-associated gene set are detailed in **Supplementary Table 1**. Black bars for each group denote the mean number of genes across species. uSPG, undifferentiated spermatogonia; dSPG, differentiated spermatogonia; SPC, spermatocyte; SPD, spermatid.

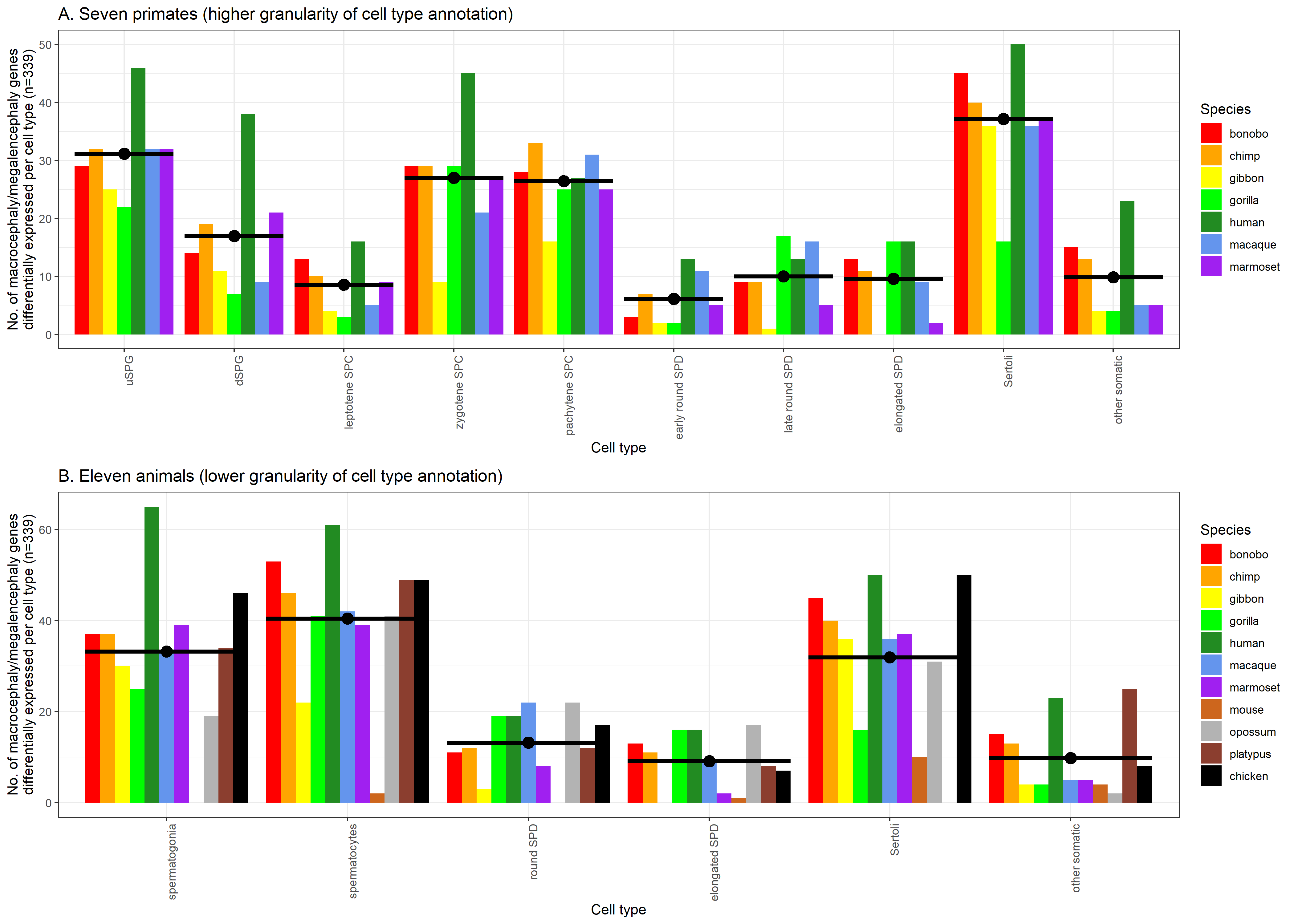

**Supplementary Figure 19. Number of macrocephaly/megalencephaly-associated genes that are also testicular cell type markers in (A) 7 primate, and (B) 11 animal species.**

This figure represents a re-analysis of data from ^219^. To compare gene names across species, we only plotted those that could be assigned to an orthogroup, defined as a set of genes whereby every member had a high-confidence one-to-one orthologue with at least one other member. Orthogroups are defined in **Supplementary Table 8** with their differential expression in each testicular cell type per species detailed in **Supplementary Table 9**. The contents of the gene set are detailed in **Supplementary Table 1**. Black bars for each group denote the mean number of genes across species. uSPG, undifferentiated spermatogonia; dSPG, differentiated spermatogonia; SPC, spermatocyte; SPD, spermatid.
